# Designing a high-resolution, LEGO-based microscope for an educational setting

**DOI:** 10.1101/2021.04.11.439311

**Authors:** Bart E. Vos, Emil Betz Blesa, Timo Betz

## Abstract

Microscopy is an essential tool in many fields of science. However, due to their costs and fragility, the usage of microscopes is limited in classroom settings and nearly absent at home. In this article we present the construction of a microscope using LEGO^®^ bricks and low-cost, easily available lenses. We demonstrate that the obtained magnification and resolution are sufficient to resolve micrometer-sized objects and propose a series of experiments that explore various biophysical principles. Finally, a study with students in the age range of 9 to 13 shows that the understanding of microscopy increases significantly after working with the LEGO microscope.

## Introduction

The invention of microscopy in the seventeenth century by Antoni van Leeuwenhoek was the start of an era of research on the micro-world [1]. Although we know by now what the “little animals” are that he observed, the micro- and nano-world is an inexhaustible topic for biophysical research, and for that the (light) microscope is the instrument of choice.

Despite the simplicity of a basic light microscope, the fundamental working principles are beyond everyday intuition for pupils, but also for most adults. While this lack of insight into simple optics might be partially due to the perception of microscopes as complex research instruments, hands-on experience promises to drastically improve the understanding for the working principles of microscopes by any interested person, including children.

Our aim is to introduce a microscope to both individual students and in a classroom setting, as a scientific tool to access the micro-world and to understand the fundamental principles of the optical components of a microscope in a playful and motivating, yet precise approach. By basing the design on LEGO, we aim to make the microscope modular, cheap and inspiring. The LEGO brick system provides a low entry level for children, as it is a common toy found in most homes. The modular design, flexibility and high level of sophistication of the different building parts make it an ideal framework to demonstrate even complex instruments with simple means. Indeed, LEGO has been used before to demonstrate the working principle of scientific instruments, such as a conceptual AFM microscope [2] and a Watt-balance [3], or in actual scientific setups [4]. Furthermore, this article is in line with other initiatives to create high-performance, low-cost microscopes. While a previous version of a LEGO-microscope, called the Legoscope, was successful in generating high quality images, it still required an objective and custom (3D-printed) parts, hindering its capacity to be used a simple way to demonstrate the principles of a microscope. Another initiative used very simple, but smart designs to generate a 1$-foldable microscope (foldscope) [5], where the low adaptability of the microscope and the image quality are a trade-off for the extraordinary low price. Finally, affordable, robust tools such as pocket microscopes and clip-ons for smartphones also allow students hands-on access to microscopy.

In this article we present the design of a fully functional microscope made entirely out of LEGO. The only non-LEGO components are the two optical parts, which can be purchased off-the-shelf at approximately €4 each. We quantify the optical performance of the microscope and find a sub-micrometer resolution. We show how, thanks to the flexible design, the magnification of a microscope can be tuned by combining different lenses and adjusting their distance, thus providing a simple tool for teaching fundamental optics and simple biophysical concepts. Furthermore, the simple and robust design allows long-term imaging without drift of the focus of the microscope.

We design a number of experiments that can be carried out using this LEGO-microscope and easily accessible materials, while exploring various biophysical themes. Finally, we demonstrate that this LEGO-microscope can be used in an educational setting by letting a group of 9-to 13-year-old students build the microscope and demonstrate their progress in understanding of microscopy with a questionnaire. To ease the usage of the LEGO-microscope, we designed a step-by-step workflow for a school setting or for helping parents and children to discover biophysics and microscopy in an autonomous way. All materials, including parts lists, building plans, workflows and project suggestions are found in the Supplementary Information and in [6].

## Scientific and Pedagogical Background

Microscopy is a fundamental tool of modern science that is used on a daily basis to study and understand biological and biophysical questions. In sharp contrast to its importance, general layman understanding of the fundamental principles remains superficial, and microscopes are often seen as complex instruments, exclusively usable in a science context. Although the working principles of a microscope are taught in high-school settings, the actual usage of microscopes in the lecture setting is often not targeting an understanding of its principles, but simply using ready-made microscopes.

Here we aim to teach in a playful way the magnifying properties of a single lens and a series of two lenses. This is effectively an understanding of the lens equation:

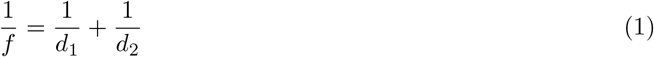

with *f* the focal length of a lens and *d*_1_ and *d*_2_ the distance between the lens and the object or the observer, respectively. Magnification occurs when the ratio 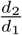 is greater than 1. We demonstrate that even children from 9 to 13 years of age can intuitively grasp this concept of optical systems. This knowledge is brought into practice with the aid of a widespread and brick-based modular toy system, called LEGO. The flexibility and the already present experience of using LEGO to build complex assemblies lies at the core of the LEGO-based workflow. We combine two lenses with adequate sample illumination and a system to precisely control the position of the lenses, which are the essential components of an optical microscope. During the building process, the difficulties in object positioning, lens adjustment and illumination become obvious, and are solved in a simple way.

Besides an understanding of optics, we suggest a series of experiments that can be conducted with the home-built microscope and readily available samples. These experiments demonstrate some fundamental biophysical concepts, such as the drastic effects of osmotic pressure on plant cells. Furthermore, the motion of easily-obtainable microswimmers can be observed, where the fundamental problem faced by swimming at low Reynolds number may be integrated in the experimental description.

To make the work with the microscope an efficient experience, we provide a detailed work plan that can be followed at the individual but also at the class level. As we provide all information as freely accessible and extensible GitHub repositories [6], we hope that the instructions might be translated to further languages by the community. We have already provided versions in English, German and Dutch.

## Materials and methods

### Microscope assembly

We designed the microscope by taking advantage of the modular and flexible characteristics of the LEGO brick system. The design of the microscope is shown in Fig. 1a-b. The building plans, including a list of components, can be found in Supplementary File 1 and Supplementary File 2. The main components are the illumination part, the objective holder, including the objectives, and the ocular part.

**Fig 1.**
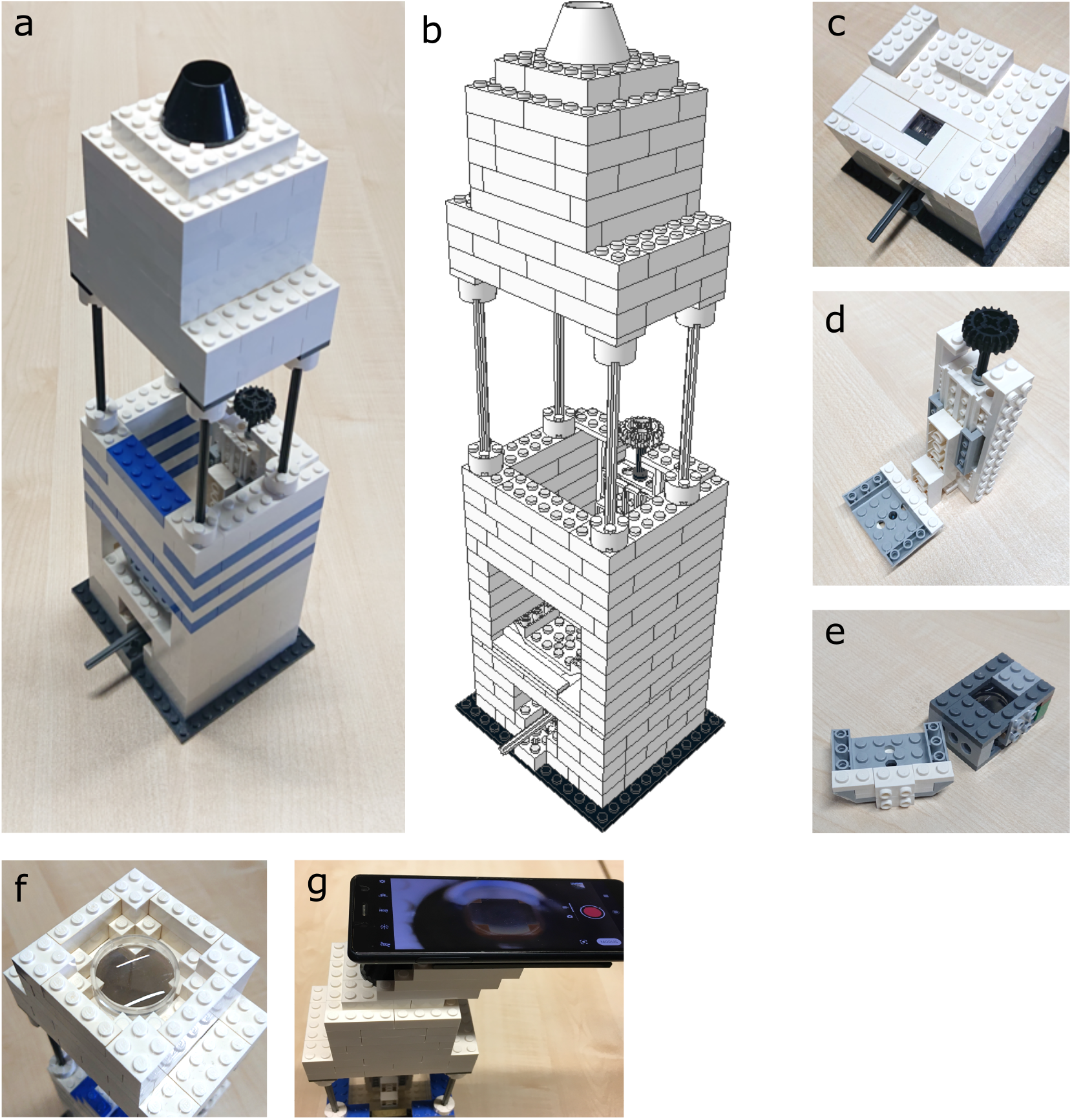
Design of the LEGO-microscope. **(a-b)** A photograph and a schematic representation of the microscope, **(c)** the LED that illuminates the sample from below, **(d)** the threaded system that adjusts the focus of the microscope by moving the objective, **(e)** two objectives, containing a replacement smartphone lens with a 3.85 mm focal distance (left) and a glass lens with a 26.5 mm focal distance (right), **(f)** the second lens, consisting of two acrylic lenses, in its holder just below the eye piece, **(g)** by adapting the eyepiece, a smartphone can be used as camera.

#### The illumination

is among the simplest parts of the microscope (Fig. 1c). In case of a group work to construct the full microscope, this part can also be built by young children that are less used to complex building instructions. The illumination itself is a special LEGO brick integrating an orange LED. It is straightforward to replace the orange LED with a different color. A short video demonstrating the opening and replacement of the LED is given in Supplementary Movie 1. As the light source is positioned close to the sample, no special condenser is included, which helps reducing the optical parts to a minimum. Due to the absence of a condenser, it is possible that the illumination with the lower magnification objective becomes slightly nonuniform. To avoid this, a diffuser can be made using thin sheet of paper that is placed between the LED and the sample.

#### Objective holder

To allow precise focus adjustment even with a high-magnification objective, an accurate and smooth positioning with sub-millimeter precision is required (Fig. 1d). As LEGO is not designed to generate such precise motion, we had to push the limits of the brick system. This was possible by combining a gear rack with a gear worm screw which results in approximately 3 mm of travel for every full rotation. This was sufficient for the final adjustment of the focus, even when looking at samples of only several μm in size. While the objective holder is the smallest of all individual parts of the microscope, it is the most demanding one with respect to building skills. Hence, building this part should either be done by older and more experienced children, or the building should be supervised by an adult to prevent frustration in an early phase of constructing the microscope. Once built, the gear system will move the objective in the vertical direction upon rotating the control wheel (black wheel in Fig. 1d). The precision and repeatability of this threaded system suffices for both objectives presented below.

#### The ocular

is a key element of the microscope, which is in the proposed classroom or exploration setting built first and then used as a classical magnifying glass. Although the overall construction of the ocular remains quite simplistic, the positioning of the lenses is still demanding. Although the lenses used here are cheap yet powerful, plastic lenses have the drawback that mishandling will rapidly lead to scratches on the surface. Users need to be careful when handling the lenses, since scratches on the surface disturb the image quality.

#### Optics

The only non-LEGO components in the building plans are the lenses in the objectives and the ocular. The designs for two different objectives are included (Fig. 1d), with a higher and a lower magnification. However, other lenses or even lens systems can be easily integrated. However, it is important that such adapted objectives position the optical elements in the same position, to keep the alignment of the microscope. For the high-magnification objective (Fig. 1d, left), the lens of a replacement iPhone5 camera module is used. Such plastic lenses that are highly specialized by a non-spherical shape have outstanding optical performance compared to glass or polymer beads and are suitable for high resolution microscopy that approaches the diffraction limit [7], while also being available at a low price. The lens used here has a reported focal length of 3.85 mm, leading to a numerical aperture of 0.54 as discussed in the Results section. After carefully peeling the lens off the chip, it is attached to a LEGO brick using transparent tape and a glass cover slip. Detailed photographs of this procedure are shown in Supplementary File 1.

For the low-magnification objective (Fig. 1e, right), a glass lens with a focal length of 26.5 mm and a numerical aperture of 0.29 was used, which was built into a holder that shares the attachment brick and also has the same optical center as the high-magnification objective.

The ocular consists of two acrylic lenses with a focal length of 106 mm each, with one flat and one convex side. Placing the two lenses with their flat sides on top of each other, the focal length is reduced to 53 mm (see Fig. 1f).

#### The sample holder

is a surface of flat LEGO bricks. This surface can be covered with rough tape to reduce the risk of accidentally moving the microscope slide. Furthermore, advanced students are encouraged to design a stage to accurately position the microscope slide. Either the gear rack and worm screws of the objective holder, or a translation stage such as presented in [2], can be used as a starting point.

Table 1 summarizes the list of non-LEGO components used in the assembly of this microscope, including their costs in euro at the time of writing. A smartphone is not included but recommended for image acquisition. In case no LEGO parts are already available the typical costs of the full set of parts is also listed.

**Table 1.** List of components used in the assembly of the microscope. The cost in euro is given at the time of writing and does not include shipping.

| Part name | Price (€) |
| --- | --- |
| iPhone 5 replacement camera | 4 |
| Acrylic lens (2x)<br>(diameter 34.5 mm, $f = 106$ mm) | 3 |
| Glass lens<br>(diameter 18.0 mm, $f = 26.5$ mm) | 5 |
| Set of microscopy slides (50) and cover slips (100) | 10 |
| All LEGO parts | 80 |

### Sample preparation

#### Salt crystals

To create sodium chloride crystals, a 1 molar sodium chloride solution is prepared by dissolving 5.8 gram of kitchen salt in 100 mL lukewarm water. A drop of the solution is placed on a microscope slide and left at room temperature for approximately an hour, during which the water evaporated and salt crystals formed.

#### Onion cell monolayer

A monolayer of red onion cells is obtained by cutting a red onion with a sharp knife, and carefully peeling off a thin slice using a pair of tweezers. This layer is then spread out on a microscope slide and covered with a drop of water and a cover slip. The cover slip is sealed off on two opposing sides with nail polish and left open on the two other opposing sides for the in- and efflux of liquid. Supplementary Movie 2, Supplementary Movie 3 and Supplementary Movie 4 show the preparation procedure.

#### Microswimmers

A selection of water-based micro-organisms such as amoeba, paramecia and water fleas are obtained by collecting water from a local pond in a plastic bottle. Since water fleas feed on bacteria and algae, some grass was added as a source of nutrients for bacterial growth. The mixture was then left for one week at room temperature, so that the water fleas could multiply. Afterwards, a few drops were placed on a microscope slide, which was covered with a cover slip and sealed off with nail polish. Since harmful bacteria can also grow in the bottle, strict hygiene must be followed, and careful washing of hands after contact with the water is important. Alternatively, paramecia and water fleas can be bought from pet shops.

#### Artemia

Eggs are obtained in powder form from a pet shop and put in water to revitalise them. After 2-3 days the organisms have hatched and a drop of water with the Artemia is placed on a microscope slide which is covered with a glass cover slip.

### Workflow for classroom and individual exploration

To support the usage of the LEGO-microscope and to optimize the learning experience, we designed a workflow plan. Such workflows can be included either in a classroom environment by separating a class in several groups, where 2-3 pupils work on a single LEGO-microscope set, where the project is split into different sets. Alternatively, the workflow can also be followed in an individual setting. Since the help of an experienced adult is not guaranteed, the workflow includes help sections, to ensure that certain steps are successful before the next steps are initialized. This is important to avoid frustration, in case some questions or tasks turn out to be difficult. The full workflow, translated in several languages, is found in Supplementary File 3 and in [6].

## Results and Discussion

The microscope can be used both for direct observations by eye or images can be recorded with a smartphone camera. For the images in this article, a Sony Xperia XZ2 Compact smartphone was used. Its rear camera has a chip of 5057 × 3796 pixels with a pixel size of approximately 1.2 *μ*m. To record images or movies, the Android camera app was used. In all experiments, the autofocus setting was used.

The following subsections quantify the specifications of the microscope. Although this is important to assess the capabilities of the microscope, this might be beyond the interest of middle school students.

### Measuring the magnification

To start the characterization of the LEGO microscope, its magnification is experimentally determined. To achieve that, images are taken of objects with known length, with and without lenses in the microscope (Fig. 2). The magnification is then given by the ratio of the respective millimeter-to-pixel ratios:

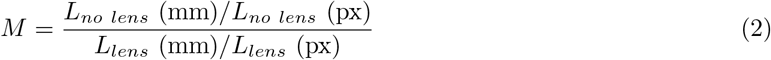

where *L_no lens_* and *L_lens_* are the length of the structure in an image without and with lenses, respectively, measured in units of millimeters (mm) and pixels (px). The millimeter-to-pixel ratio was obtained from squared paper with 5 mm squares for the microscope without lenses, and a 1951 USAF Resolution Test Targets (Thorlabs, Dachau, Germany) was used to obtain the millimeter-to-pixel ratio when the lenses were in place. For the high-magnification objective we find *M* = 254×. For the low-magnification objective we find *M* = 27×.

**Fig 2.**
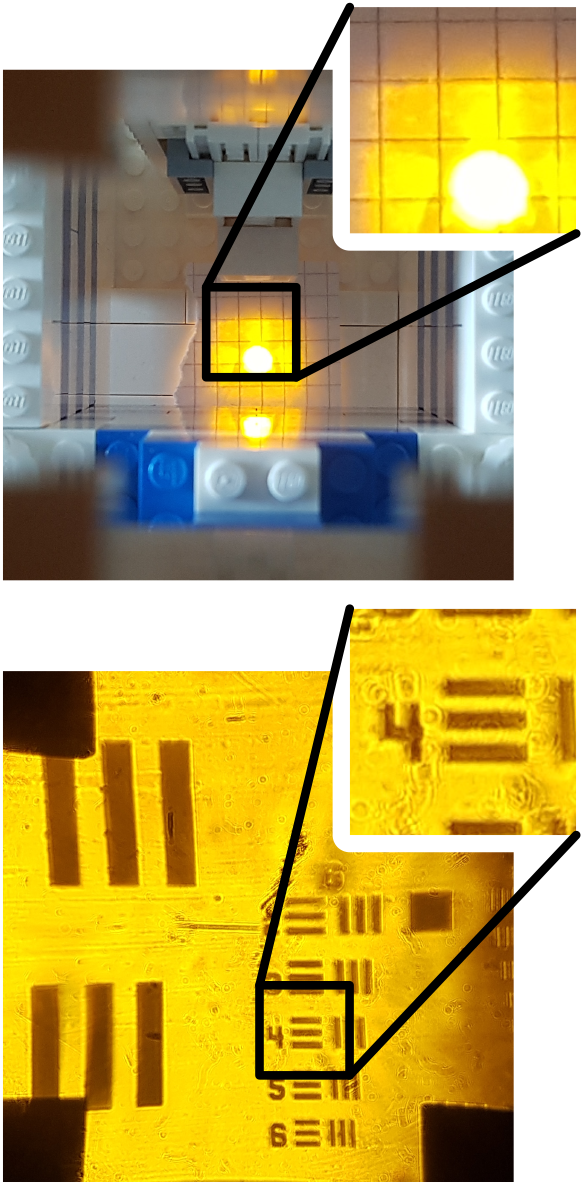
Experimentally determining the magnification of the microscope. **Top:** Imaging of squared paper without lenses in place. The width of each square is 5 mm. **Bottom**: Imaging of the USAF Resolution target with the high-magnification objective and a *f* = 53 mm lens near the eye piece. The length of the bar next to the number four is 27.6 *μ*m. Note that for a correct determination of the magnification, the (digital) zoom of the camera has to be the same for both images.

### Calculating the magnification

The magnification of the microscope can also be calculated based on the distances between the object, lenses and the observer. In Fig. 3 the light path of the microscope is schematically drawn. The magnification *M* of the first lens is simply given by the ratio of the lens to the intermediate image distance *d_intermediate_* and the lens to object distance *d_object_*:

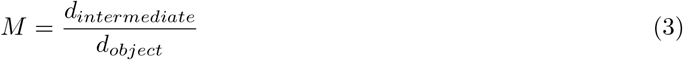

**Fig 3.**
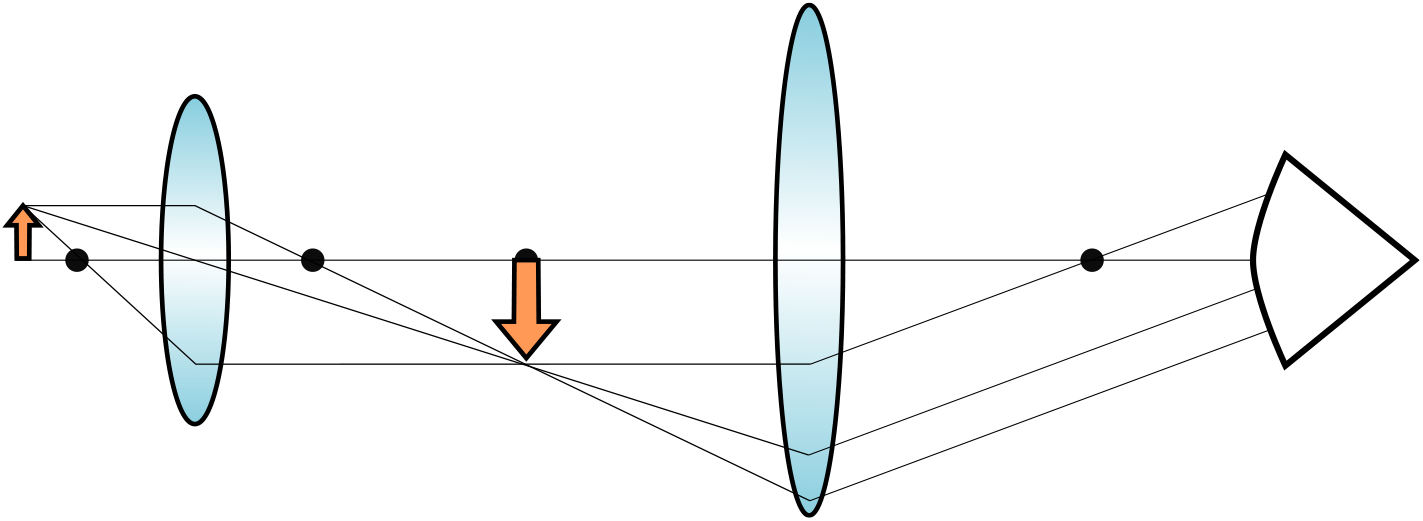
Schematic overview of the light path in the microscope. The object (here depicted as an arrow) forms an inverted intermediate image in the focus of the second lens. The second lens then sends collimated light to the observer.

The distance between the object and the second lens was measured with a ruler and found to be 240 mm. Since the intermediate image is located in the focus of the second lens, the object-to-intermediate-image-distance *d_object_* + *d_intermediate_* = 240 — 53 = 187 mm. To determine *d_object_* and *d_intermediate_* individually, we have to use the lens formula (Eq. 1), which can also be expressed as:

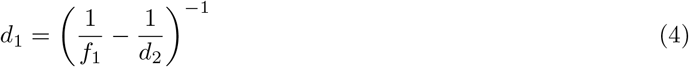

Since the sum *d_intermediate_* + *d_object_* is known, *d*_2_ can be substituted:

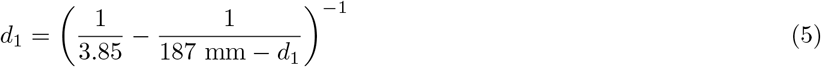

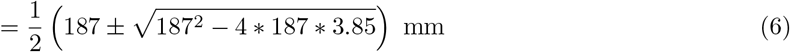

which shows that *d_object_* = 3.9 mm and *d_intermediate_* = 183 mm. When substituting these numbers into Eq. 3, the magnification of the smartphone lens is 47×. This magnification is then further enlarged by the second lens, which magnifies it by a factor of the near distance to the eye (around 250 mm for adults [8]), divided by the focal length of the second lens, here 53 mm. That makes the total magnification *M* = 220 ×, in good agreement with the value that was experimentally determined.

The total magnification using the low-magnification objective with the glass lens is determined in a similar way, substituting *f* = 26.5 mm in Eq. 5 for the focal length. In this case, *d_object_* = 32 mm and *d_intermediate_* = 155 mm, leading to a total magnification of *M* = 23×, again in good agreement with the value that was experimentally determined.

### Quantifying the resolution

To quantify the optical performance of the microscope, the width of a sharp black-white transition is measured. To this end, a 1951 USAF Resolution Test Targets (Thorlabs, Dachau, Germany) was used as shown in Fig 4a-b. The intensity profile along the red lines in Fig 4a-b is shown in Fig 4c-d, where the three black bars can be clearly recognized as dips in the intensity profile. In Fig 4e-f, a close-up of the intensity profile of a single bar is shown. This is a convolution of a step-function with the Point-Spread Function (PSF) of the microscope. The Point-Spread Function of a microscope describes how a point-like object is imaged by a microscope. It depicts a fundamental limit of a microscope: when the PSFs of two objects overlap, they can no longer be optically resolved as different objects, but are imaged as one object. We approximate the shape of the PSF with a normal distribution, so that the intensity profile can be fitted to a cumulative distribution function of a normal distribution:

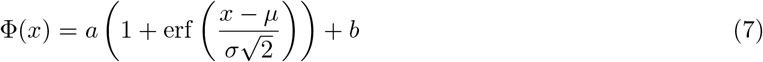

where *σ* is the deviation, *μ* the center of the normal distribution, *a* and *b* a prefactor and offset, respectively, and erf the error function. The Full-Width Half-Maximum (FWHM) of the normal distribution is then given by 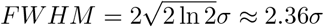. Finally, since the width of the lines in the resolution test target is known, the FWHM can be converted from pixels to *μ*m. From the high-magnification objective (panels a, c, e), we find *FWHM* = 0.78 ± 0.12 *μ*m (*n* = 6). For the low-magnification objective (panels b, d, f), we find *FWHM* = 4.3 ± 0.9 *μ*m (*n* = 6).

**Fig 4.**
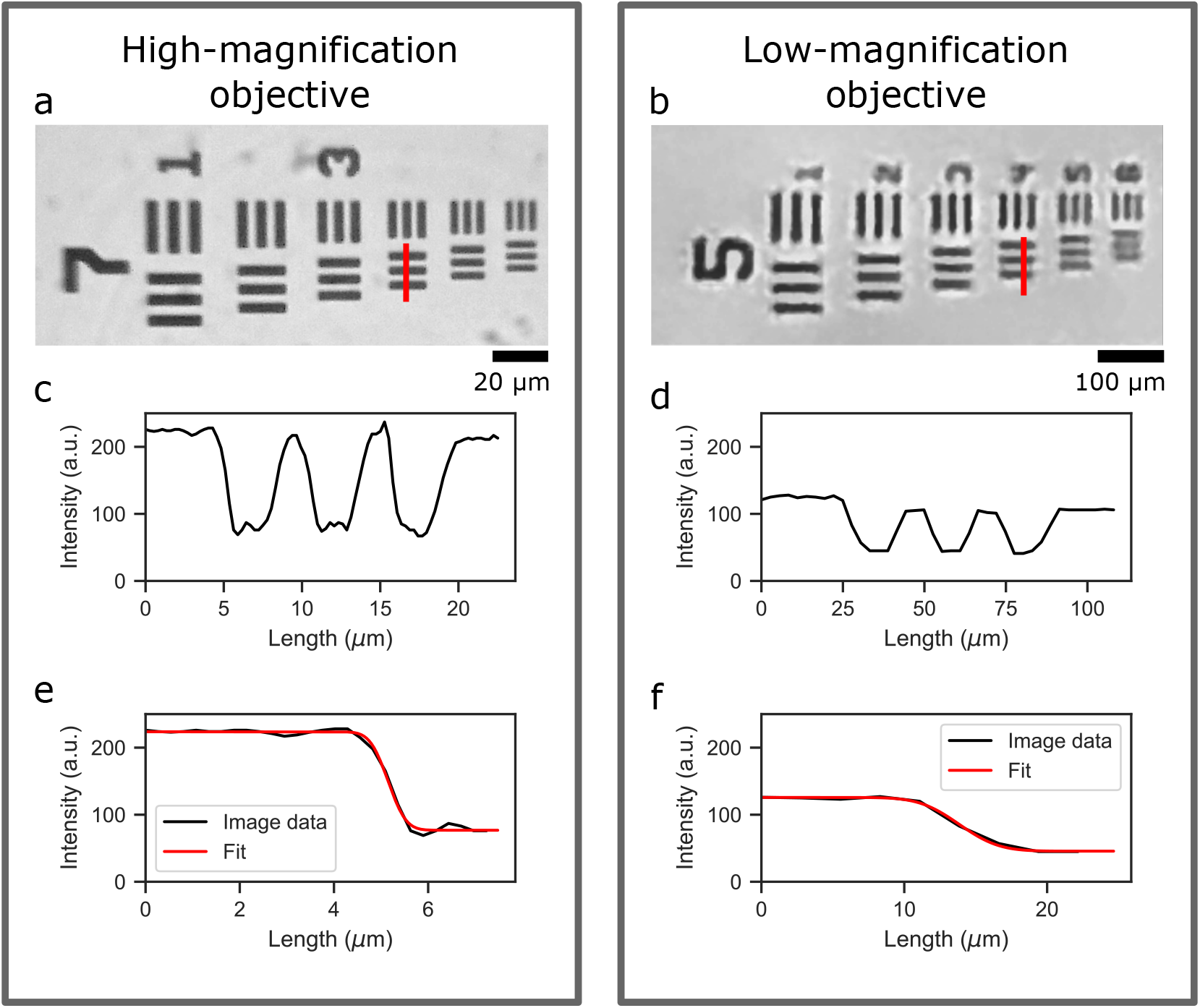
Quantifying the resolution of the microscope with a high-magnification and a low-magnification objective. **(a-b)** Using the known width of the lines in the USAF Resolution Test Target, the conversion factor from pixels to micrometers can be calculated. In these panels, the width of bars crossed by the red lines are 2.76 *μ*m and 11.05 *μ*m respectively, and the corresponding intensity profiles along the lines are shown directly below. **(c-d)** Intensity profiles along the the red lines, using the pixel-to-micrometer conversion factor obtained from the calibration slide, **(e-f)** intensity profile along a single bar (black), which has been fitted (red) with a cumulative distribution function of a normal distribution.

To compare this result to the maximal obtainable resolution, the Abbe diffraction limit is estimated for both objectives. The Abbe diffraction limit [9] states that two spots can be optically separated if they are separated by a distance larger than *d*, given by:

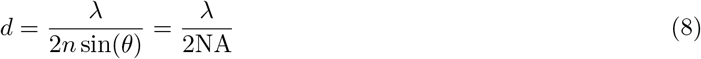

where λ is the wavelength of light, *n* the refractive index of the medium through which the light travels and *θ* the maximum angle in which that light can still be collected by the lens. The numerical aperture NA is defined as NA = *n* sin(*θ*). For the LEGO-microscope, *n* =1 since the sample and the objective are separated by air.

The angle *θ* that is needed to calculate the numerical aperture is obtained by taking the arctangent of the lens radius *r_lens_* divided by the focal length *f*,

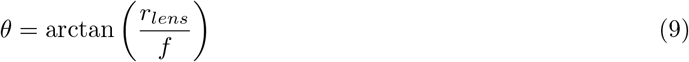

For the high-magnification objective, we find a numerical aperture of 0.54, which, combined with an average wavelength of light of approximately λ = 550 nm, leads to a resolution of *d* = 0.51 *μ*m, which is only slightly smaller than the measured 0.78 *μ*m that was estimated based on the intensity profile in Fig. 4e. For the low-magnification objective, the radius of the aperture of the LEGO brick that holds the lens needs to be used, rather that the radius of the actual lens. We find a numerical aperture of 0.29, which leads to a resolution limit of *d* = 0.95 *μ*m, a factor 4.5 smaller than what was experimentally estimated based on the intensity profile in Fig. 4f. The poorer performance of the low-magnification objective compared to the high-magnification objective is likely due to the fact that a single lens is used in the low-magnification objective rather than the composite lens system of the smartphone camera lens used in the high-magnification objective. Indeed, a smartphone camera lens is designed to provide good image quality for low and high angles of incidence, here we show that this lens functions close to the theoretically possible limit.

As an alternative to calculating the numerical aperture from the lens radius and its focal length, the numerical aperture is experimentally determined in Supplementary Figure 1. The measured value of the NA was comparable with the calculated value.

## Use in educational setting

The goal of this project is to provide an accessible, hands-on microscope that can be used in an educational setting. In a preliminary attempt to quantify the effectiveness of the LEGO microscope as a learning tool, 8 students in the age range of 9 to 13 were tested before and after following the provided workflow instructions. The tests were identical, and contained 5 questions on the subject of microscopy as well as 5 general questions related to natural sciences that served as a benchmark, see Supplementary File 4. Although the subject of the microscopy questions was discussed in the workflow instructions, the answers were not included and had to be obtained by working through the different steps.

The workflow instructions were organized in a 13-step procedure during which the students built the microscope step-by-step. The manual explores various aspects of the microscope. The workflow consists of the following basic steps:

1. **Building of the three main parts of the microscope.** The relevant parts should be built individually. At this point, the students are not necessarily aware that the final outcome will be a microscope. Objectives are not yet added in this stage. However, the lens of the ocular is already mounted.
2. **Exploration of the different parts.** The students are asked to play with the different parts and to guess what they can be used for.
3. **Focus on the ocular.** The students are asked to closely inspect the ocular. They should find out that it can be used as a magnifying glass. They should realize that because of the closed construction, typically not enough light is delivered to the sample.
4. **Combination of light source and ocular.** To overcome the problem of low light, the students should realize that using a light source, built in a previous step, produces a better view of the magnified sample.
5. **Exploration of individual samples.** The students are asked to inspect arbitrary samples with the magnifying property of the ocular.
6. **Interruption to recapitulate experiences.** The students are asked to collect the main experiences gathered so far. This ensures that the important lessons to work on the following steps have been learned. In a classroom setting this should be done together with all groups.
7. **Identifying possible ways to increase magnification.** Besides using stronger lenses, the students should figure out that two magnifying glasses might improve the total magnification. If this step was not suggested by the students, it will be introduced either by the teacher or by the help in the manual.
8. **Introduction of the objectives.** First, the students are exposed to the low-magnification objective. They should try to use both the ocular and the second lens to have an improved magnification. As this step requires very precise alignment, students will mostly fail, thus introducing the requirement of stable mounting. Several helping strategies are included for this step.
9. **Interruption to collect experiences.** The idea of fixing the lens position is discussed or introduced in the help section of the manual.
10. **Combination of all parts.** In this step, the students actually combine all parts to get a working microscope. Now they are also told that they have successfully constructed a LEGO microscope.
11. **Exploration of samples with the microscope.** Samples are explored, as suggested in the sample part of this paper or provided by the teacher.
12. **High-magnification objective.** To improve the magnification, the second objective is introduced at this stage. Due to the lens appearance, the similarity with a magnifying glass is lost. However, after integration in the microscope, micrometer-sized objects can be visualized.
13. **Inspection of micrometer-sized samples.** In the final step, the students are asked to inspect samples from the micro-world. Biophysical effects, such as osmotic pressure and the motion of micro-swimmers, are introduced here.

After having finished this workflow, the second test was taken, asking the same questions as the initial test. The general questions serve as a control benchmark, and, as shown in Fig. 5, the fraction of correct answers of this control changes only marginally after the students have worked with the microscope. The fraction of correct answers on the microscopy questions however, almost doubles. Although the group size was relatively small, the effect is significant.

**Fig 5.**
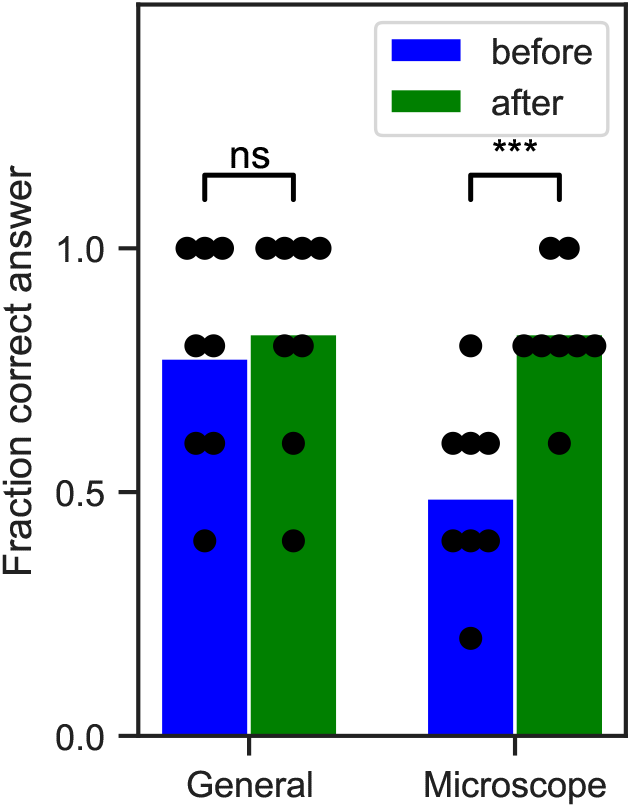
Fraction of correctly answered questions on subjects of natural sciences and microscopy, before and after using the LEGO-microscopy. The bars show the average results of 5 questions on either of the subjects, filled in by 8 students in the age range between 9 and 13, the black dots correspond to the results of individual students. There was no significant (ns) change in the results of the general science questions: from 78 ± 21% to 83 ± 21% correctly answered questions, but a statistically significant improvement (***, *p* < 0.001) was observed for the questions on microscopy, from 50 ± 17% to 83 ± 12% correct answers.

### Example experiments

One aim of the LEGO microscope is to provide children and teenagers an easy and playful access to optics and biophysics. After building the microscope, simple objects of everyday use can be imaged, such as hairs from humans and animals or natural and artificial fibers from clothing. To improve the learning effect, we furthermore propose a series of simple experiments that can be performed at home that allow discovering some of the main biophysical principles. Specifically, we suggest discovering the process of crystallisation, to observe the effect of osmosis on plant cells and to discover the motion of microswimmers.

Among the simplest, yet impressive samples is a salt crystal, see Fig. 6. The shape of a sodium chloride (NaCl) crystal reflects the face-centered cubic lattice formed by the larger chloride ions that are surrounded by the smaller sodium ions. As described in the material section, these crystals can be pre-formed and observed. Alternatively, when a thin film of the salt solution is placed on a microscope slide, the concentration of salt will slowly increase due to evaporation of water, so that the crystallization process itself can be recorded.

**Fig 6.**
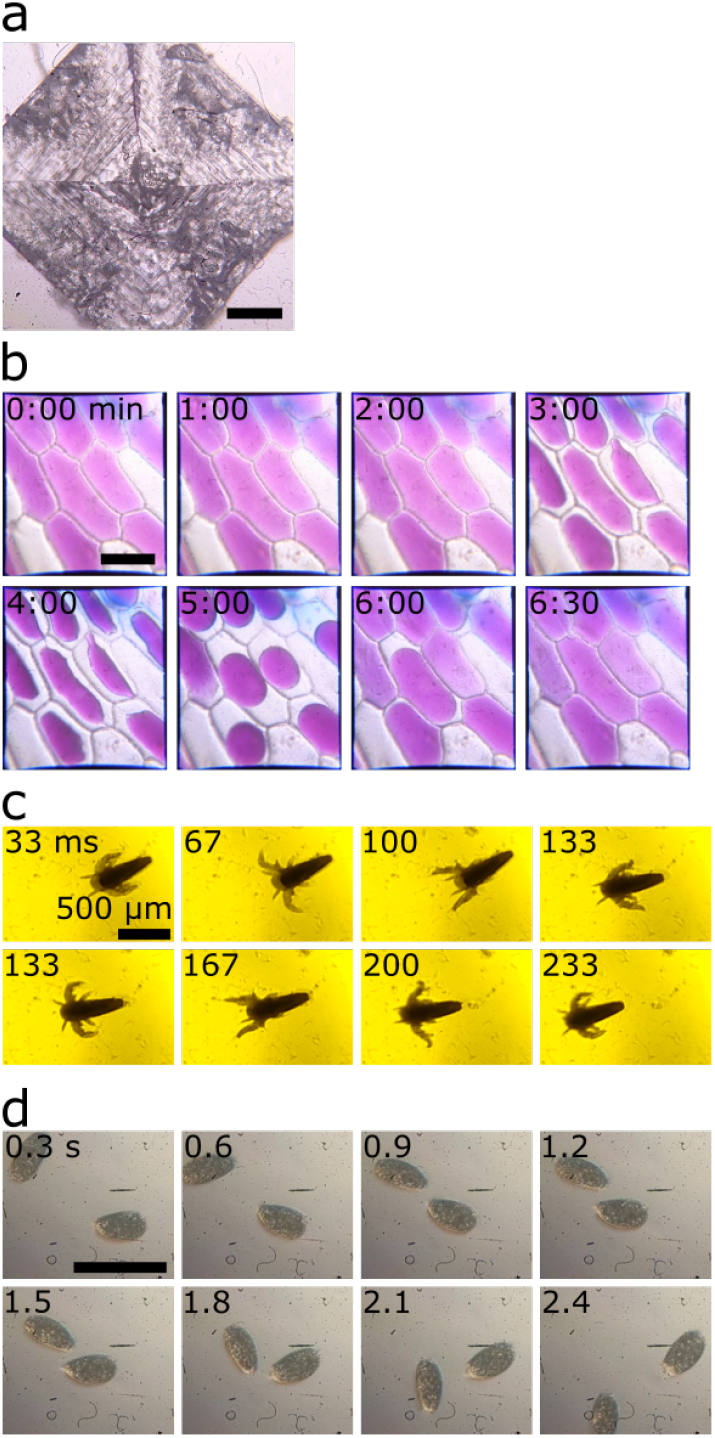
Examples of experiments conducted with the LEGO-microscope. **(a)** Image of a sodium chloride crystal. **(b)** Time lapse of an osmotic shock in red onion cells. After approximately 30 seconds, a 1M NaCl solution is flowed in. Subsequently, water leaves the cells, causing the cell membrane to detach from the cell wall. After approximately 5 minutes, distilled water is flowed in, washing away the 1M NaCl solution, and the cells return to their original volume. **(c)** Time lapse of the movement of an Artemia shrimp in water. **(d)** Time lapse of the movement of two water fleas in water. The scale bars in panels (a,b,d) are 100 *μ*m.

Osmosis is an important biophysical effect that regulates shape and pressure in many biological systems, for example the turgor pressure in plant cells that presses the plasma membrane against the cell wall and gives plant tissue its rigidity. In Fig. 6b, a time-lapse is shown of red onion epidermal cells exposed to an osmotic shock. The full movie is shown in Supplementary Movie 5. At approximately *t* = 30 s, a drop of 1M NaCl solution is placed on one open side of the cover slip which, due to capillary forces and a piece of absorbing paper on the other side of the cover slip, ensured that the cells were rapidly surrounded by the NaCl solution. This creates a large difference in osmotic pressure between the inside and outside of the cell, resulting in a loss of water. As a consequence of the volume reduction, the plasma membrane detaches from the cell wall starting around *t* = 3 min, and the pigment becomes more concentrated. At approximately *t* = 5 min, the NaCl solution is replaced with distilled water, which enters the cells and returns them to their original volume. During the recording, care was taken not to touch the microscope to avoid movement of the microscope slide. No drift in focus was observed during our measurements, avoiding the need to adjust the height of the objective, which in turn would lead to motion of the microscopy.

To discover and image systems that are currently the research focus of many biophysics groups, a series of different microswimmers can be imaged with the LEGO microscope. The propulsion of microswimmers is fundamentally different from macroscopic swimmers since it occurs in a low Reynolds number regime, where friction is the dominating force [10]. Relatively large microswimmers are Artemia shrimps. Their movement is imaged with the low-magnification objective and is shown in Fig. 6c and Supplementary Movie 6. Artemia shrimps can swim relatively fast, using their antennae for regular strokes. In contrast, unicellular organisms apply different strategies for propulsion. These organisms can be grouped in two categories: *flagellates* that use a small number of long flagella for movement, while *ciliates* have a large number of small, hair-like flagella (called cilia) covering the entire organism [11]. Flagellates are typically smaller than ciliates and they are hard to resolve with the LEGO microscope. On the other hand, the movement of ciliates can be imaged well. An example is the propulsion of paramecia, unicellular organisms that are covered with cilia and are commonly found in ponds. Their movement is imaged with the high-magnification objective and is shown in Fig. 6d and Supplementary Movie 7.

## Conclusion

Using LEGO and low-cost, easily available lenses, it is possible to construct a microscope that can resolve micrometer-sized objects, with a resolution that is close to the diffraction limit of light. A series of experiments is suggested, covering different fields of natural sciences, that can be conducted with household ingredients. The modular design of the microscope itself also lets it easily be incorporated in a curriculum on optics. A preliminary study with 8 students in the age range of 9 to 13 shows that the understanding of microscopy increases after working with the LEGO-microscope.

Since the design of the microscope presented here is only one of many possible configurations, customization is highly encouraged.

## Author contributions

BV, EB and TB designed the microscope. BV and TB carried out experiments, analyzed data and wrote the manuscript.

## Acknowledgements

This project was supported by the European Research Council (Consolidator Grant 771201). We also acknowledge teachers from Pascal Gymnasium in Münster for useful discussions and their students for testing the building instructions of the microscope and the workflow and providing helpful feedback.

## Supporting information

**Supplementary File 1.** Step-by-step building instructions for the LEGO-microscope.

**Supplementary File 2.** A list of parts for the LEGO-microscope.

**Supplementary File 3.** A manual to step-by-step explore the LEGO-microscope.

**Supplementary File 4.** A series of questions that were used to test the knowledge on microscopy before and after using the LEGO microscope.

**Supplementary Movie 1.** Instructions for replacing the LED in the LEGO brick to change the color of illumination.

**Supplementary Movie 2.** Instructions for the preparation of red onion epidermal cells for microscopy.

**Supplementary Movie 3.** Instructions for closing the sides of a sample slide for flow experiments.

**Supplementary Movie 4.** Instructions for performing an osmotic shock experiment.

**Supplementary Movie 5.** Red onion epidermal cells exposed to an osmotic shock by the addition of a 1 M sodium chloride solution.

**Supplementary Movie 6.** Movement of Artemia shrimps observed with the low-magnification objective.

**Supplementary Movie 7.** Movement of Paramecia observed with the high-magnification objective.

**Supplementary Figure 1. Experimental determination of the numerical aperture of the high-magnification objective.** A green laser is refracted by the smartphone camera lens and illuminates the backside of a paper box. By measuring the distance from the lens to the box and the diameter of the refracted circle, the maximum angle *θ* under which light left the lens is estimated. The average of multiple measurements results in a numerical aperture of 0.53, close to what was calculated based on the lens diameter and its focal length.

