## Supplementary figures and images for "Designing a high-resolution, LEGO-based microscope for an educational setting"

### Supplementary Figure 1

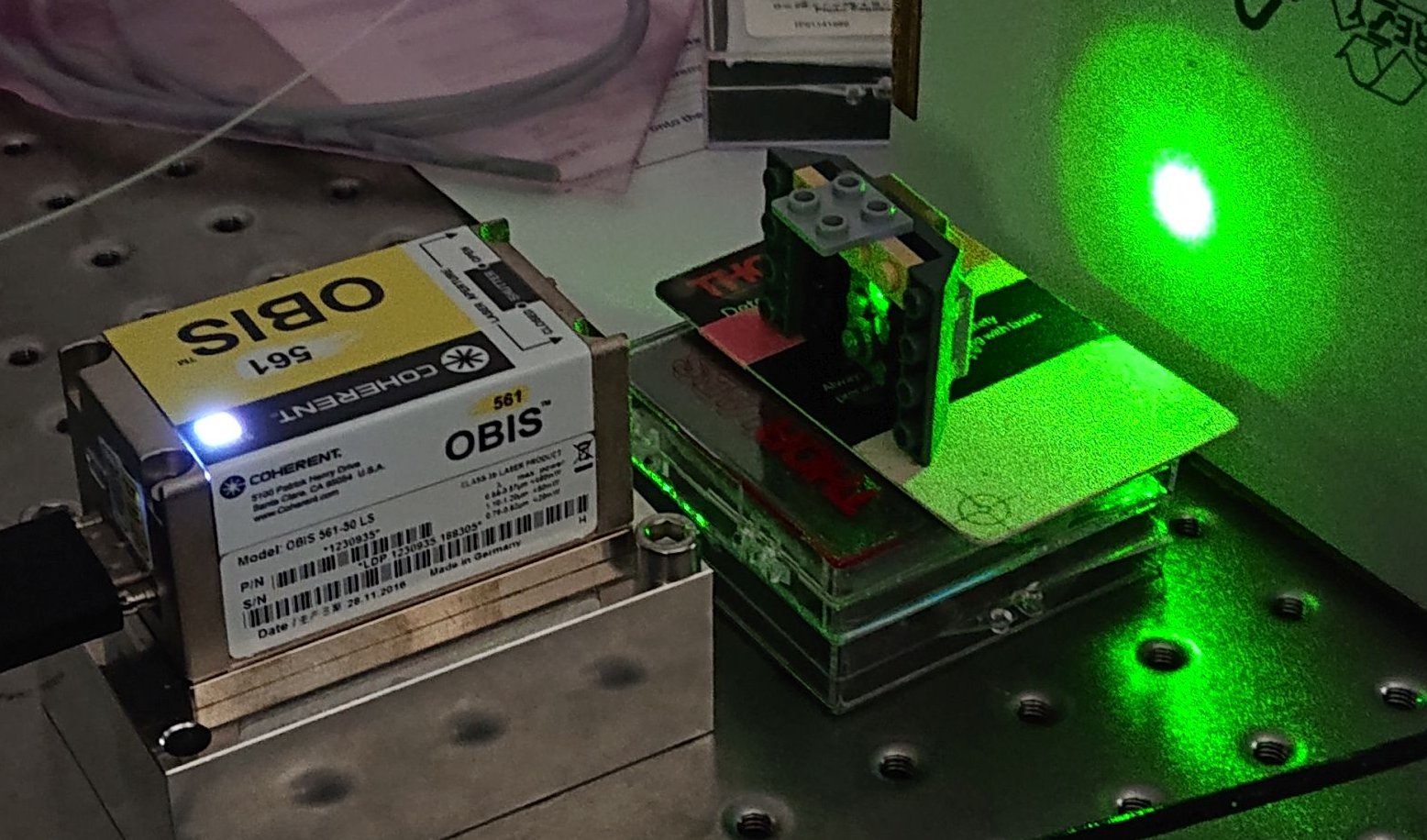
