## Supplementary File 1 for "Designing a high-resolution, LEGO-based microscope for an educational setting"

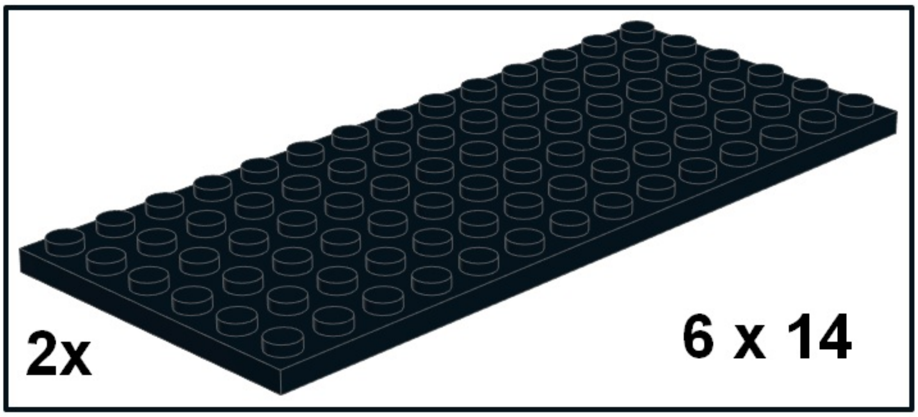


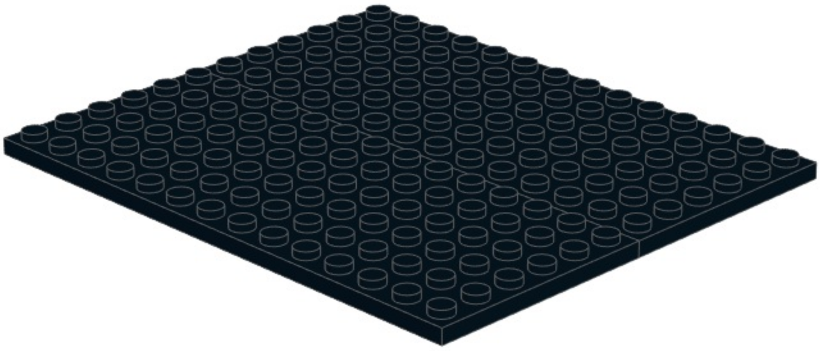

## 2

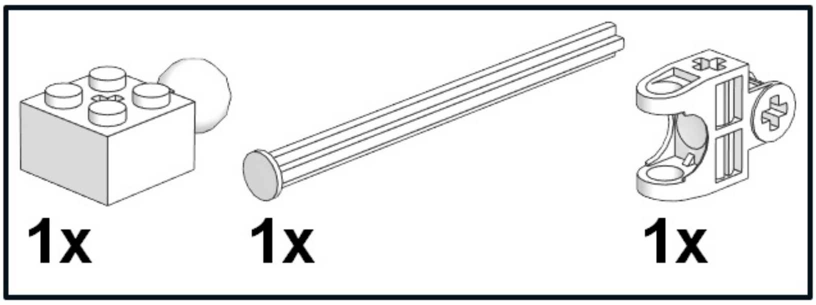


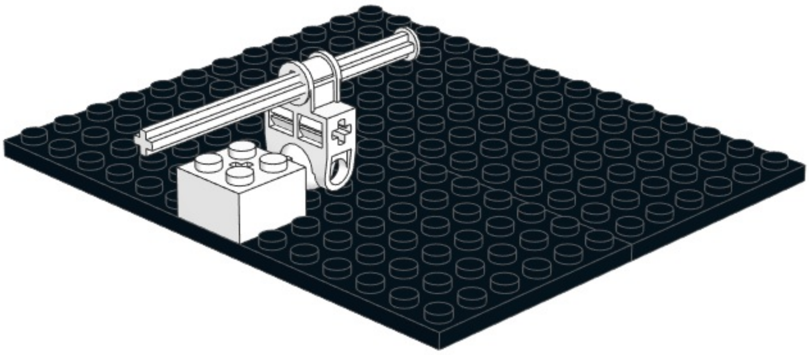

# 3

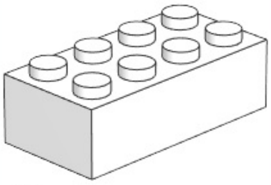

2x

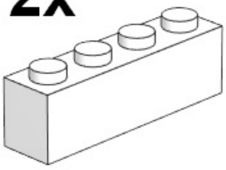

2x

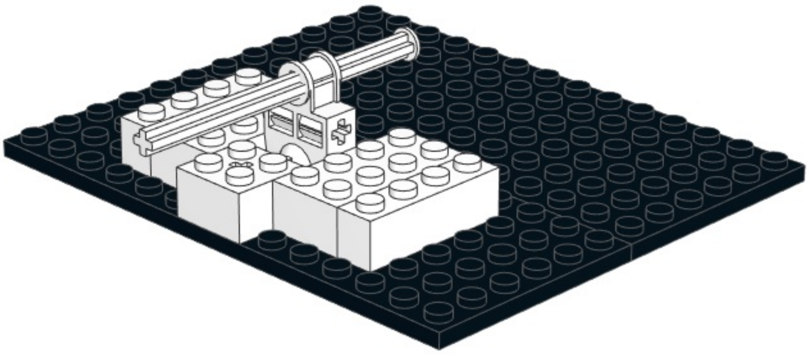

4

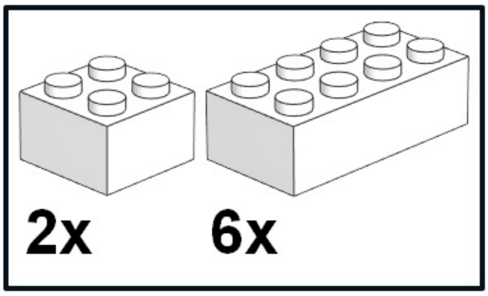


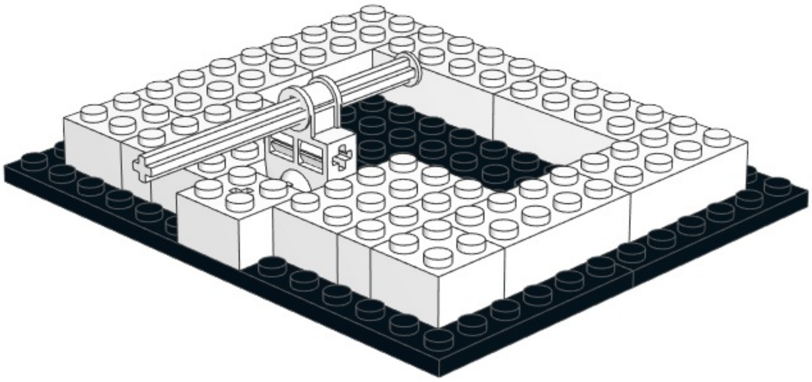

# 5

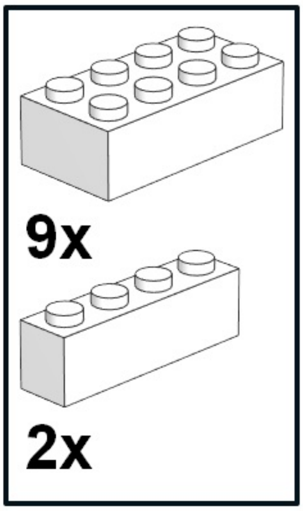


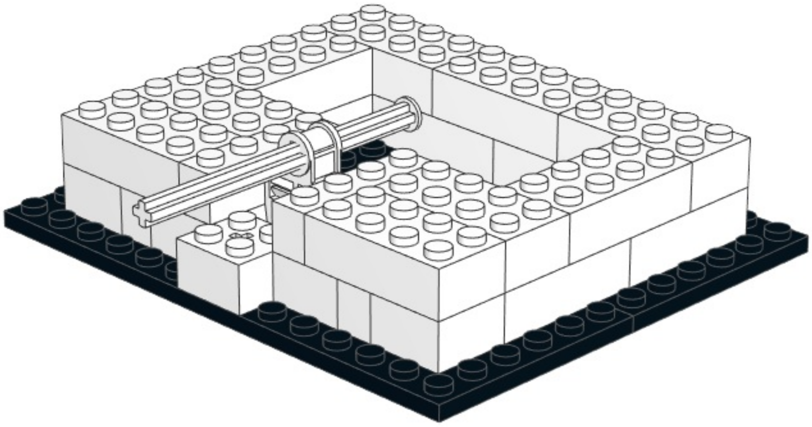

6

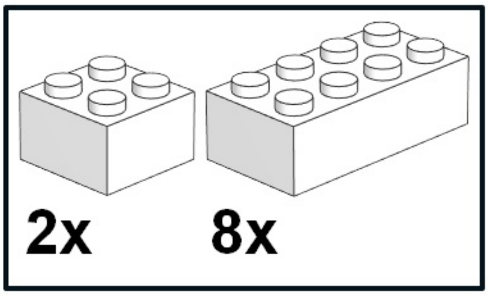


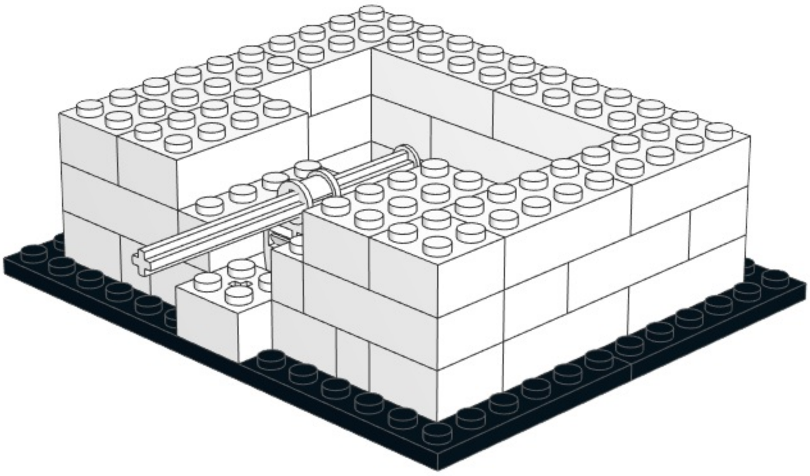

7

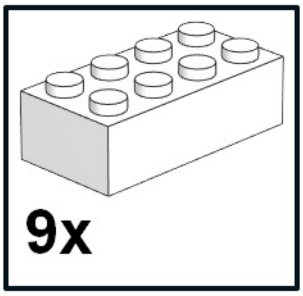


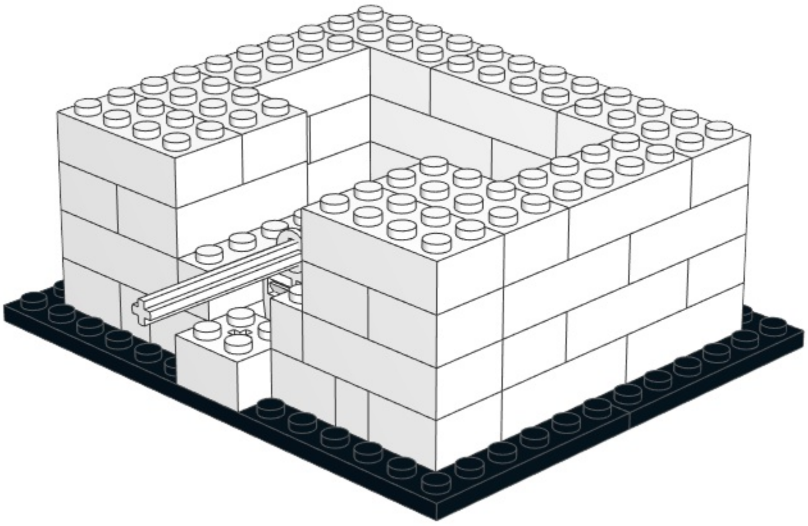

8

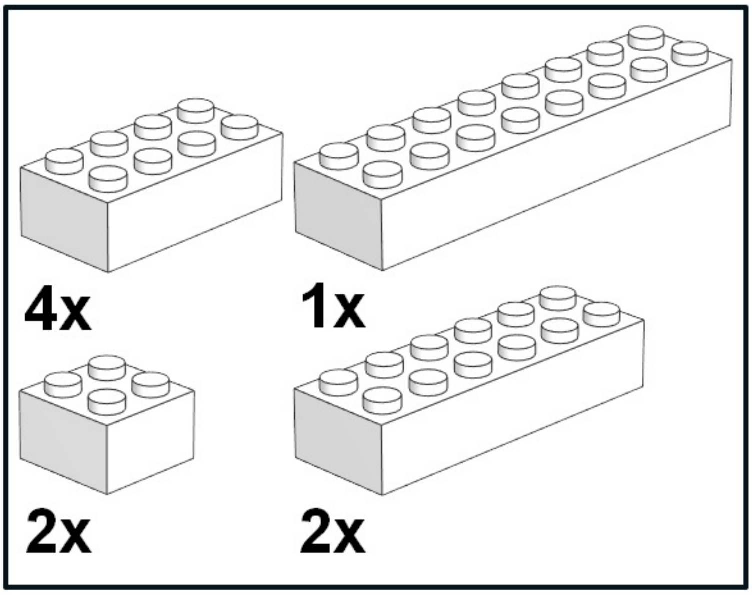


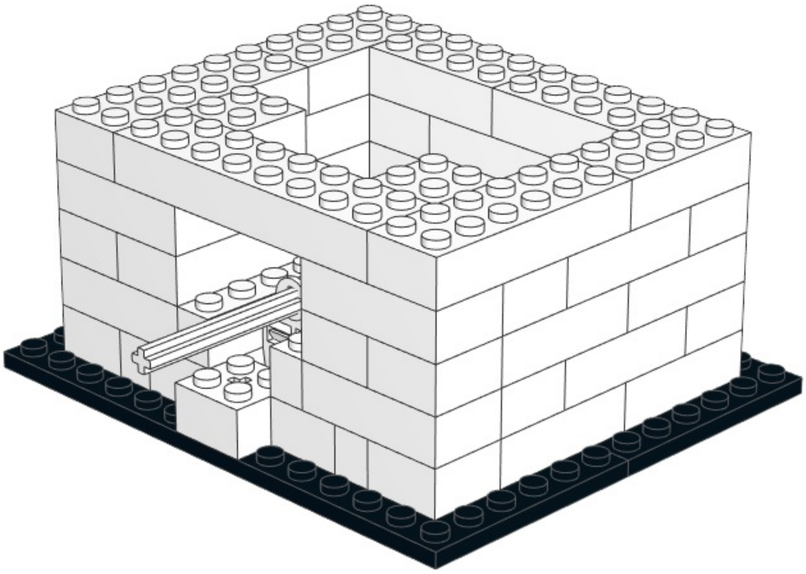

# 9

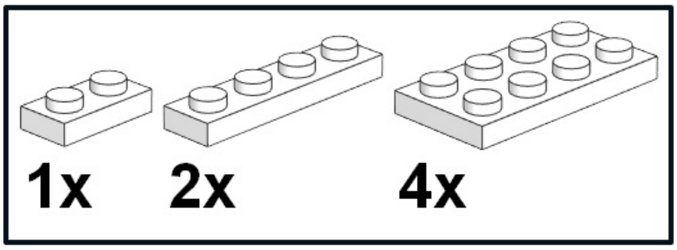


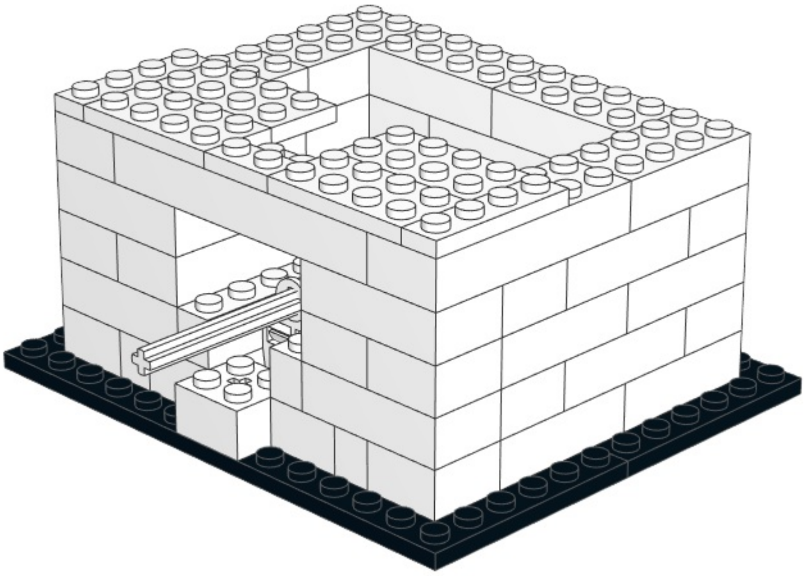

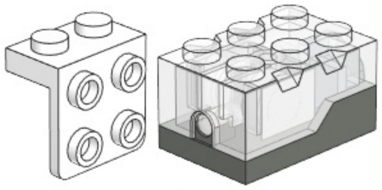

1x 1x

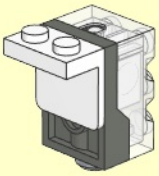

10

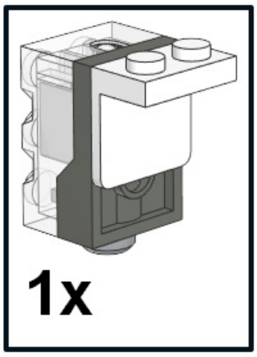


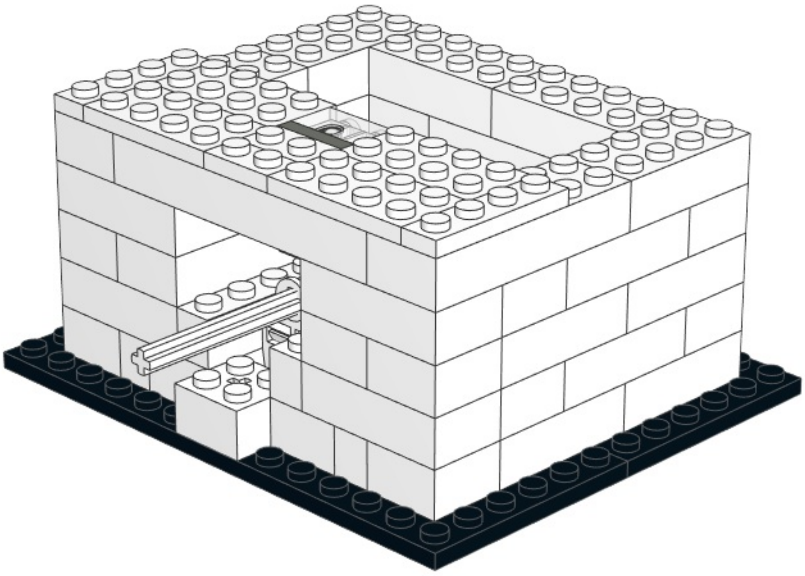

# 11

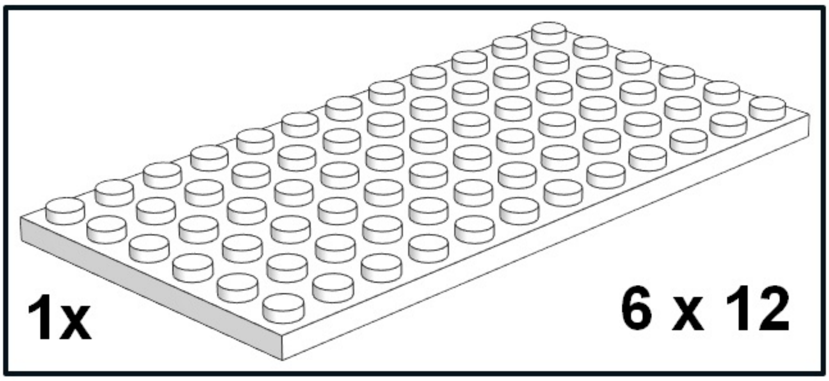


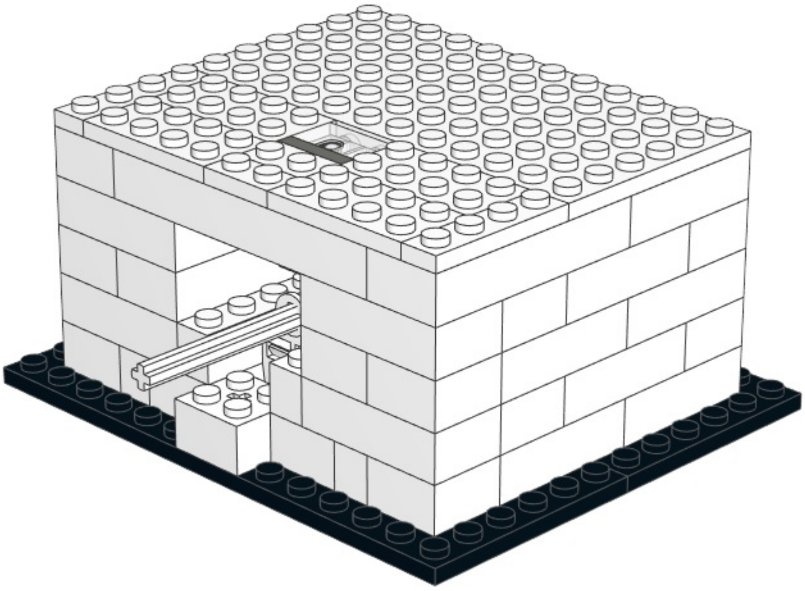

## 12

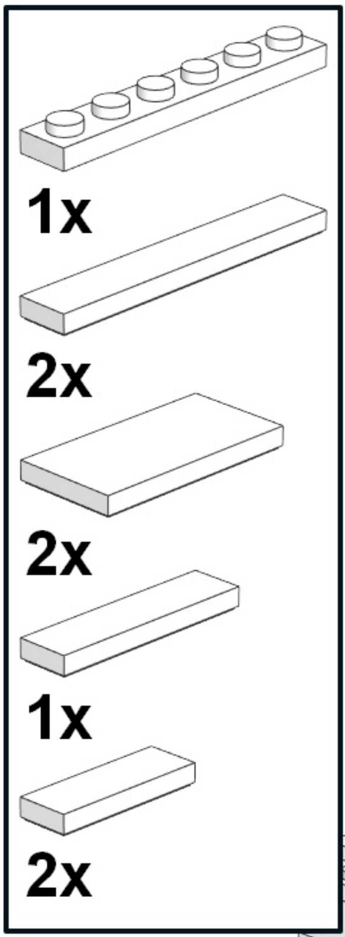


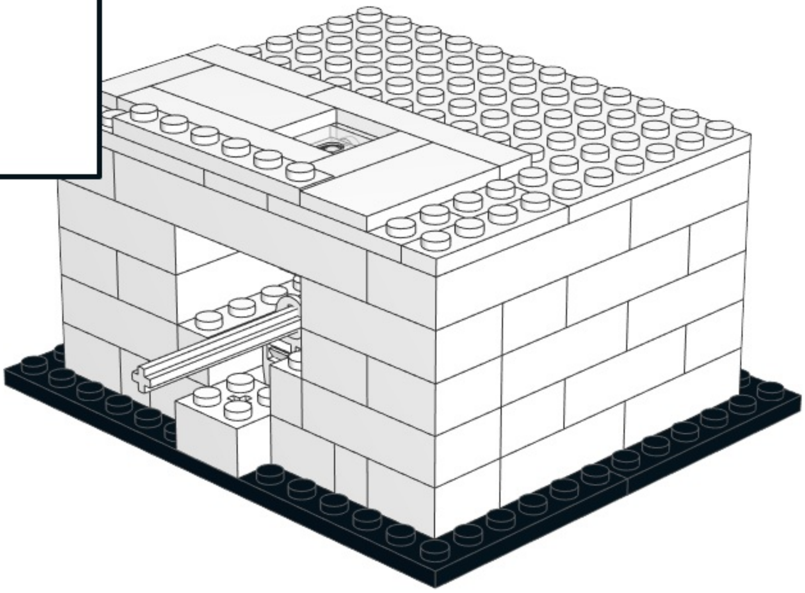

13

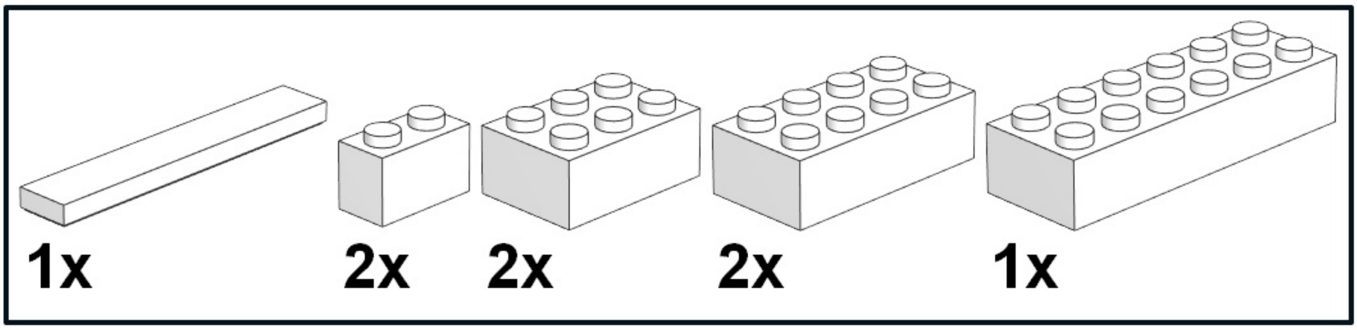


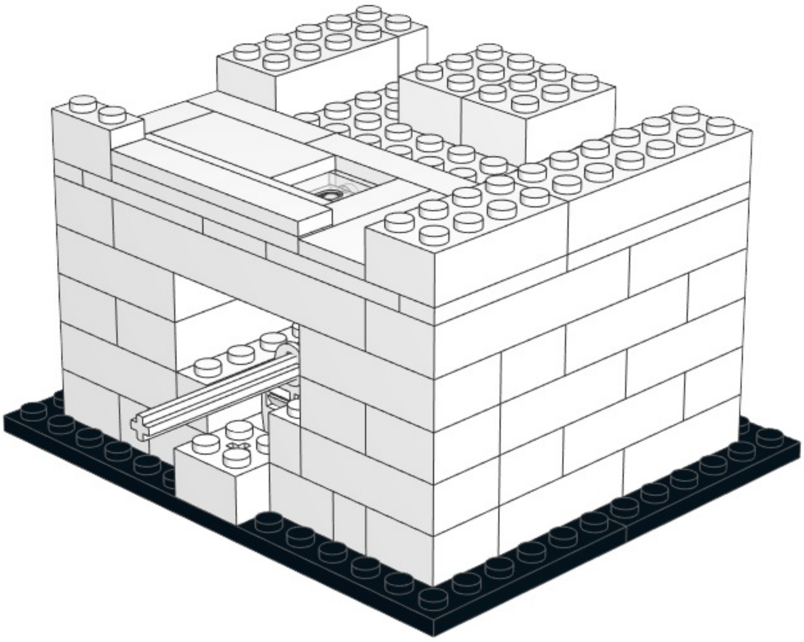

1

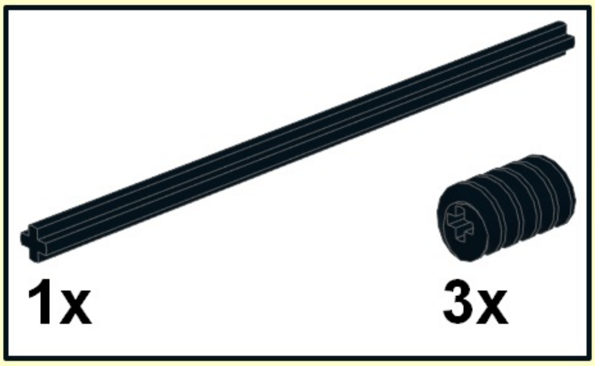


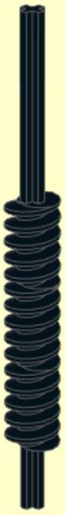

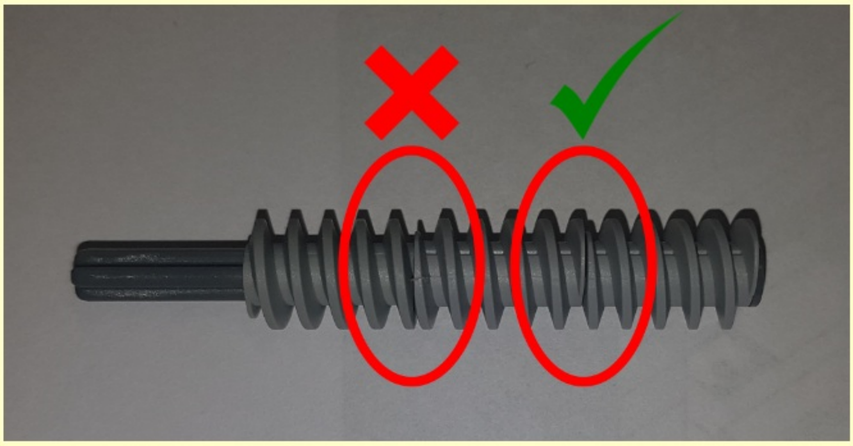

2

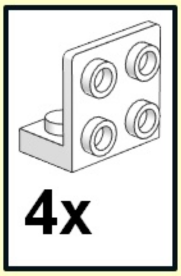


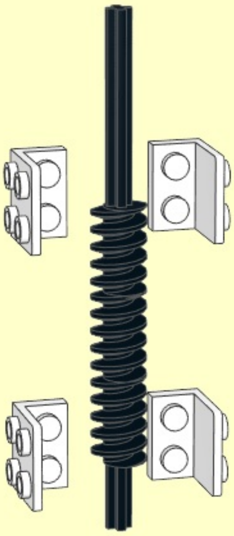

3

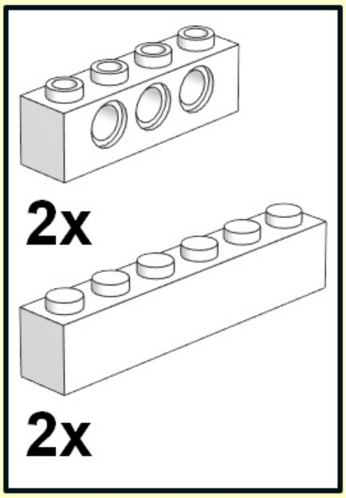


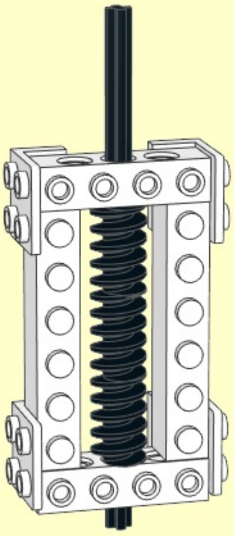

4

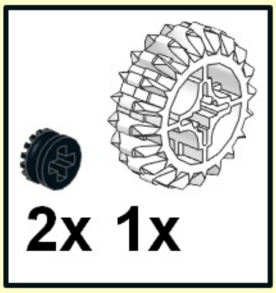


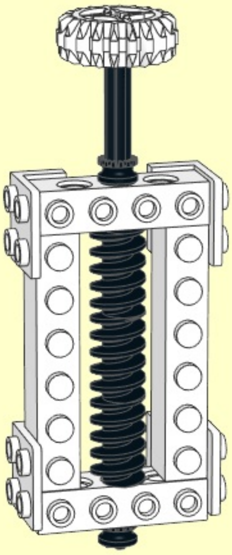

5

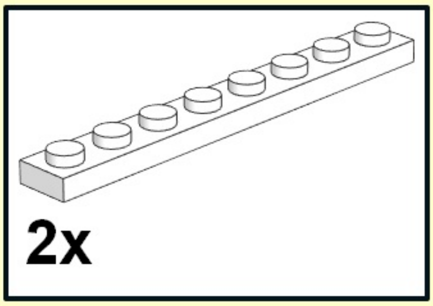


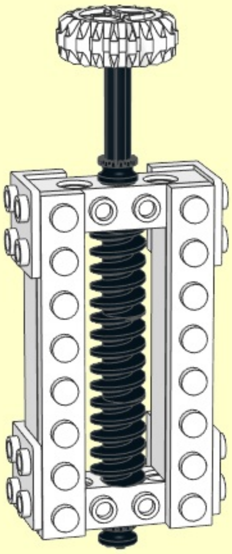

6

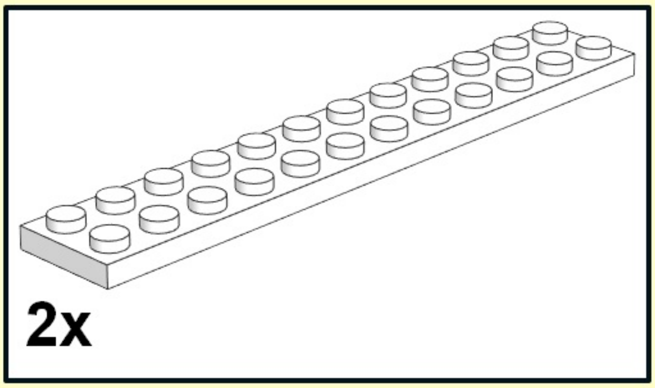


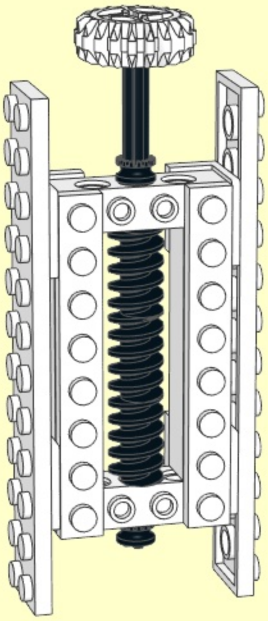

7

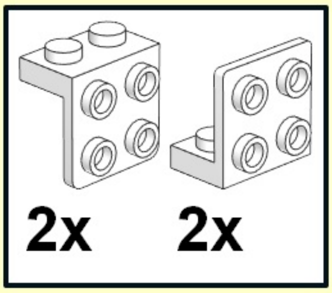


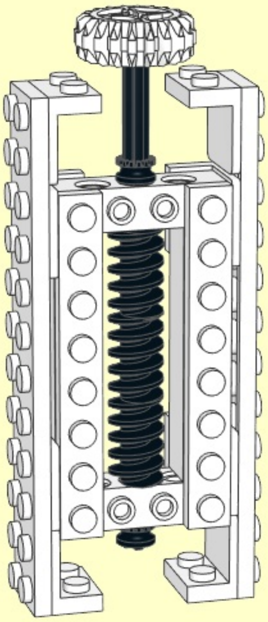

**8**

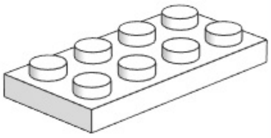

**1x**

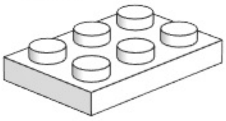

**1x**

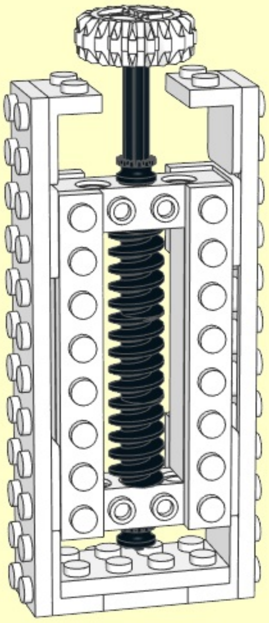

1

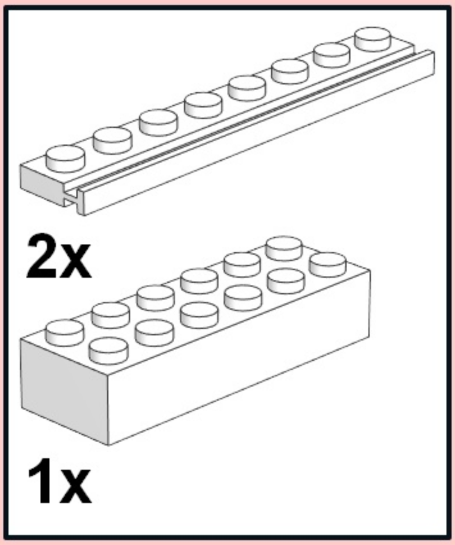


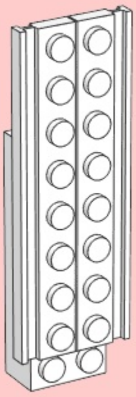

2

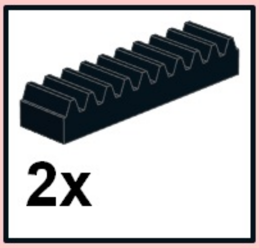


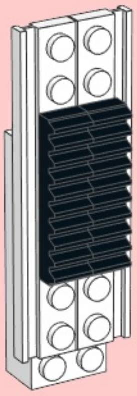

3

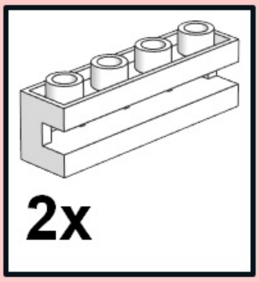


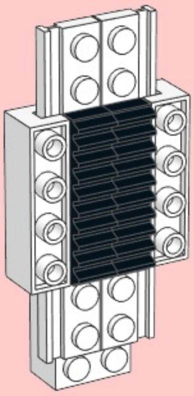

9

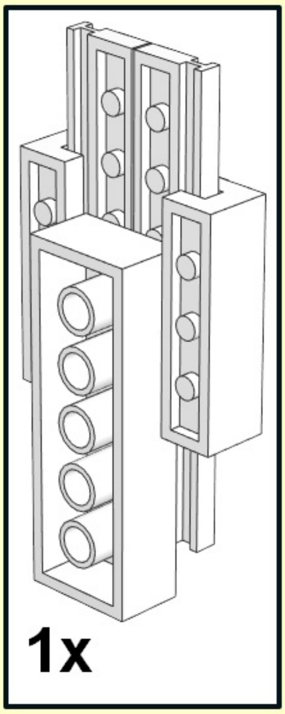


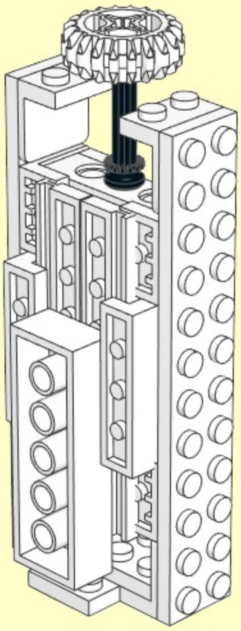

**14**

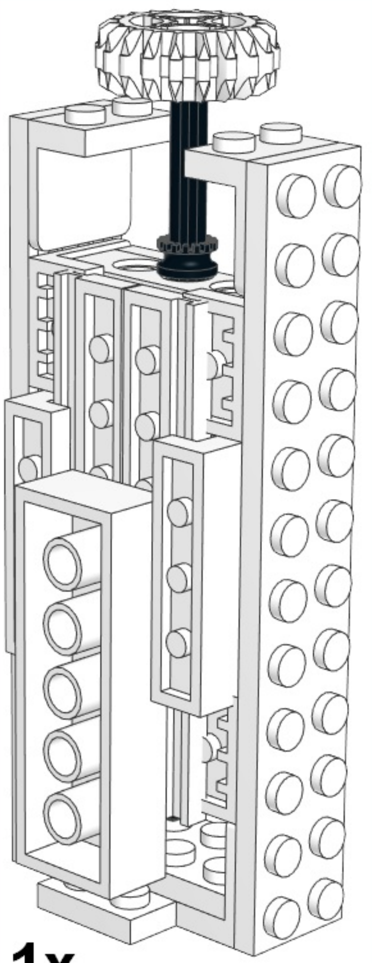

1x

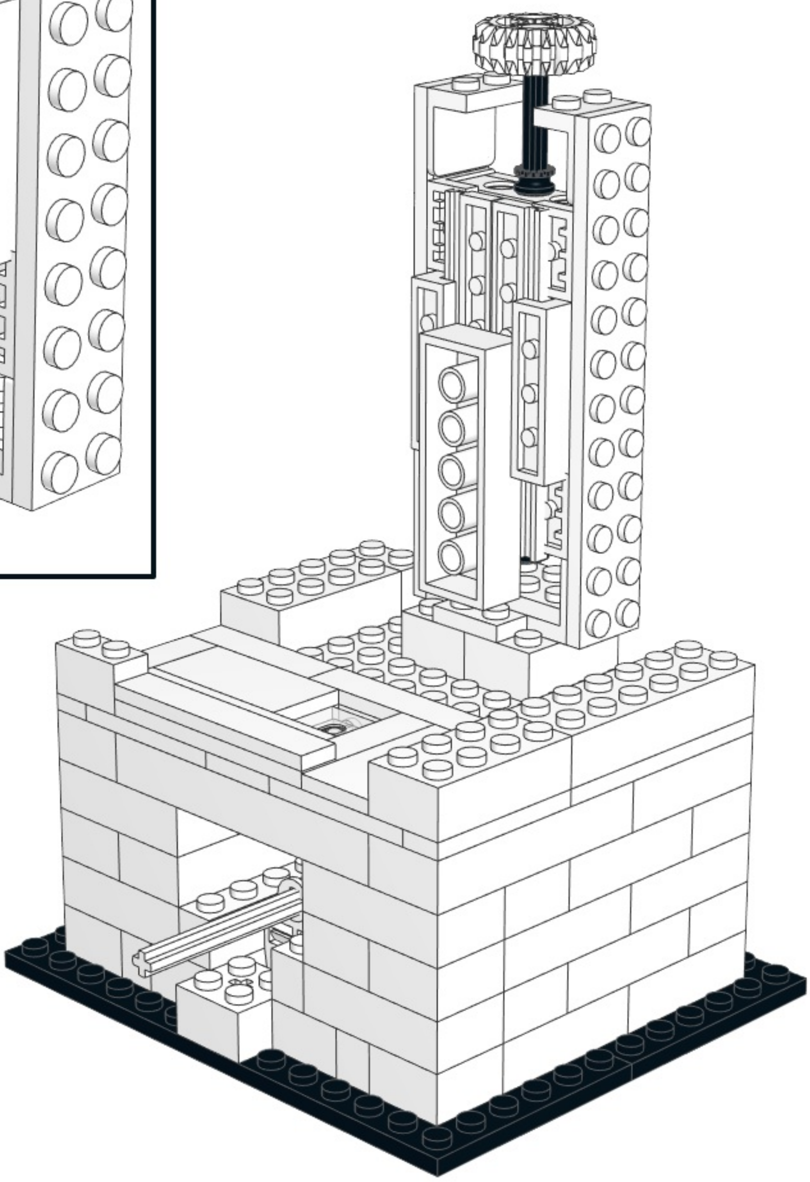

1

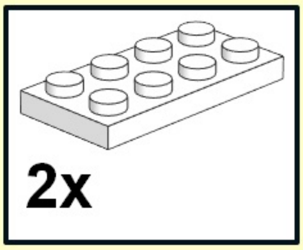


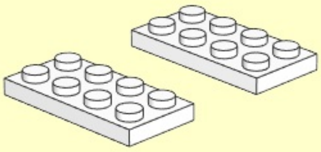

2

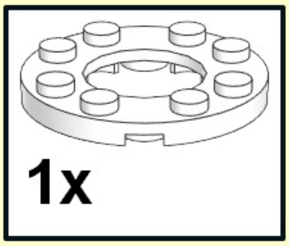


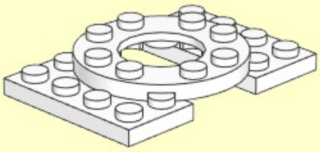

# 3

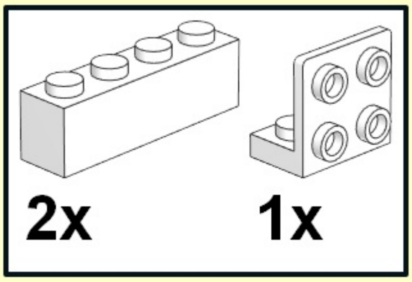


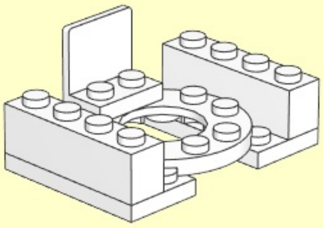

**4**

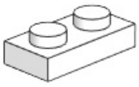

**1x**

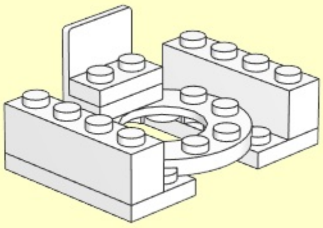

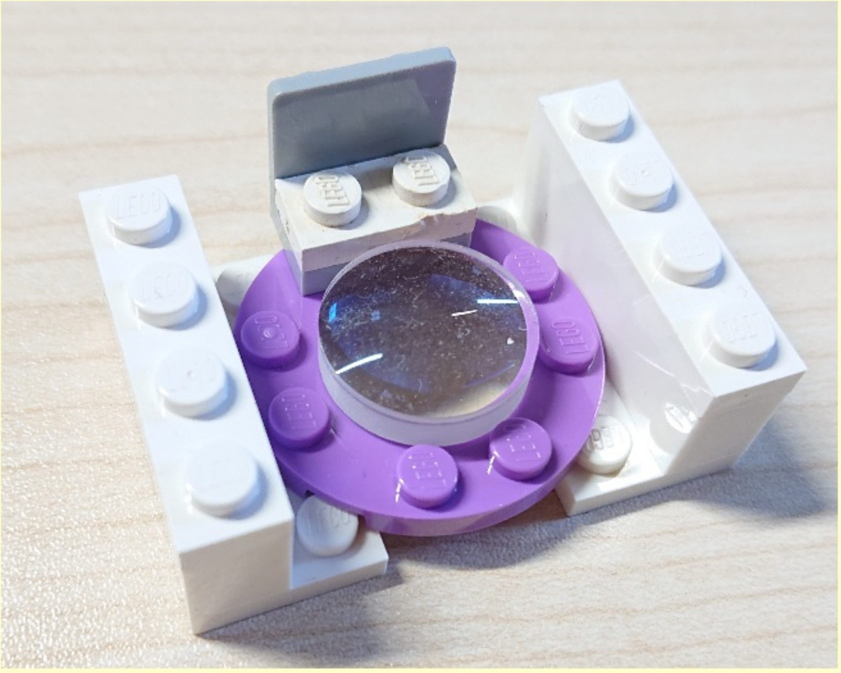

5

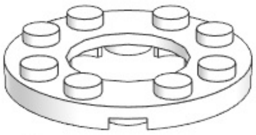

1x

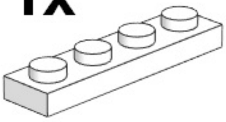

2x

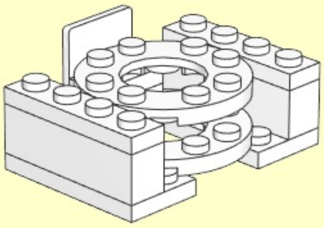

**6**

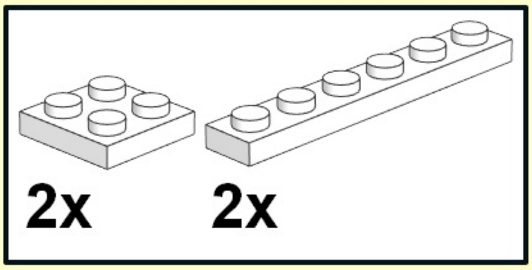


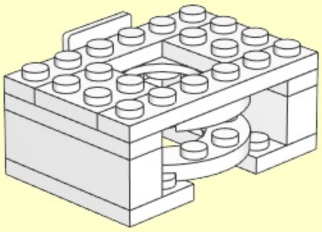

# 7

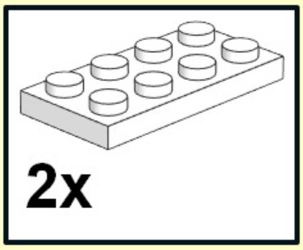


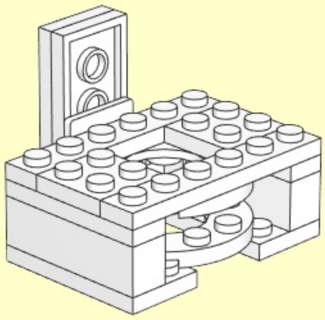

1

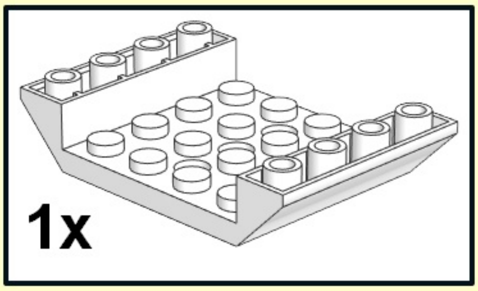


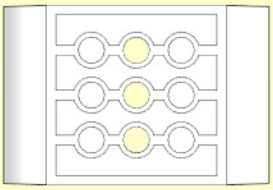

**2**

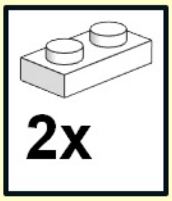


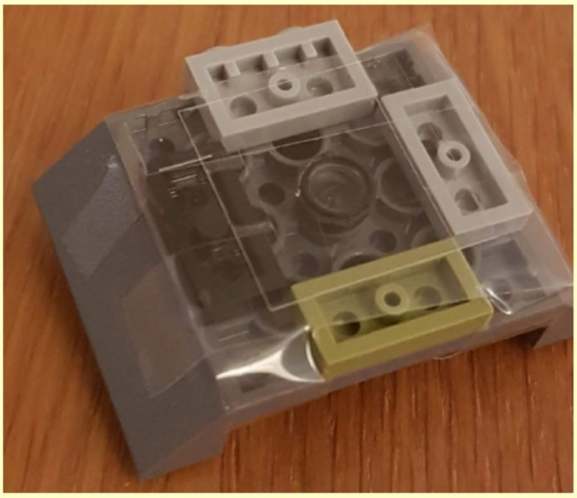

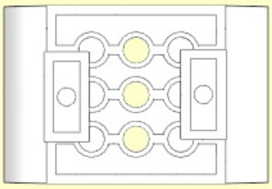

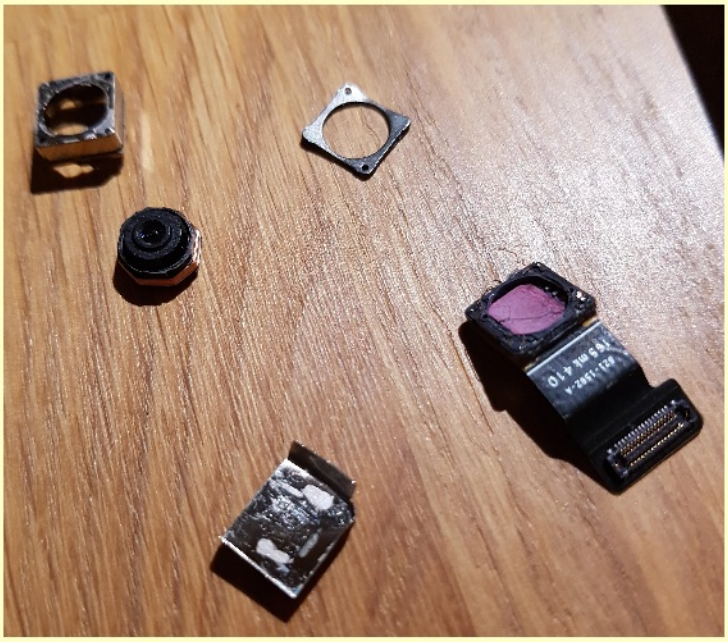

3

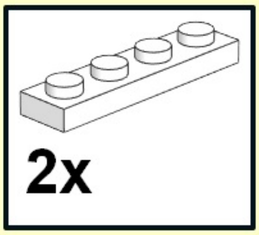


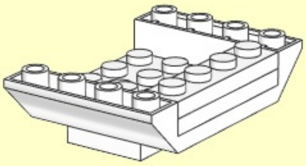

**4**

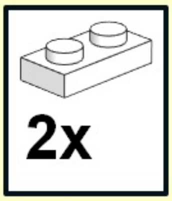


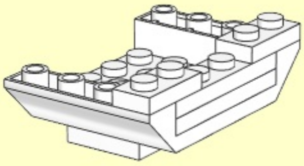

**5**

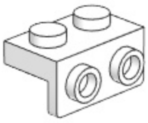

**1x**

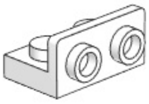

**1x**

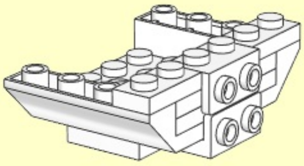

6

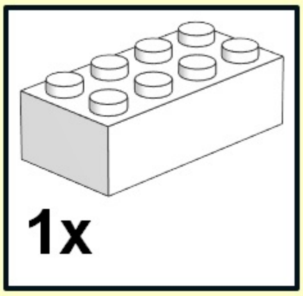


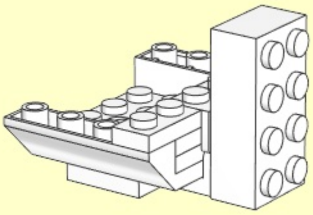

# 15

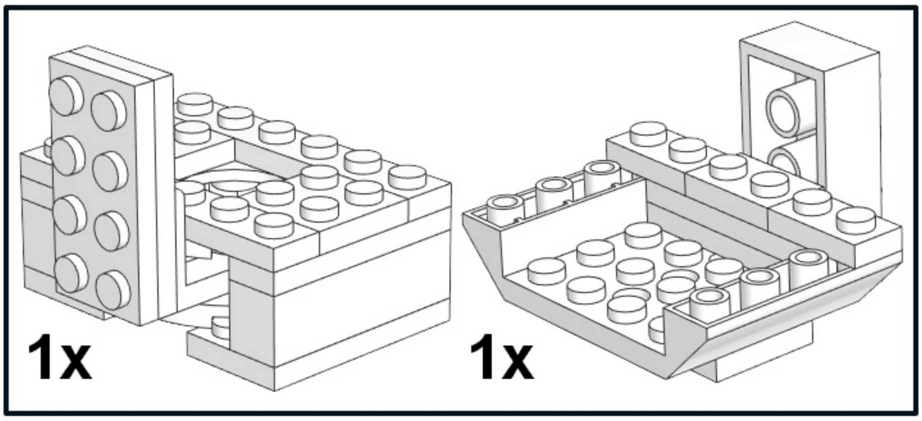


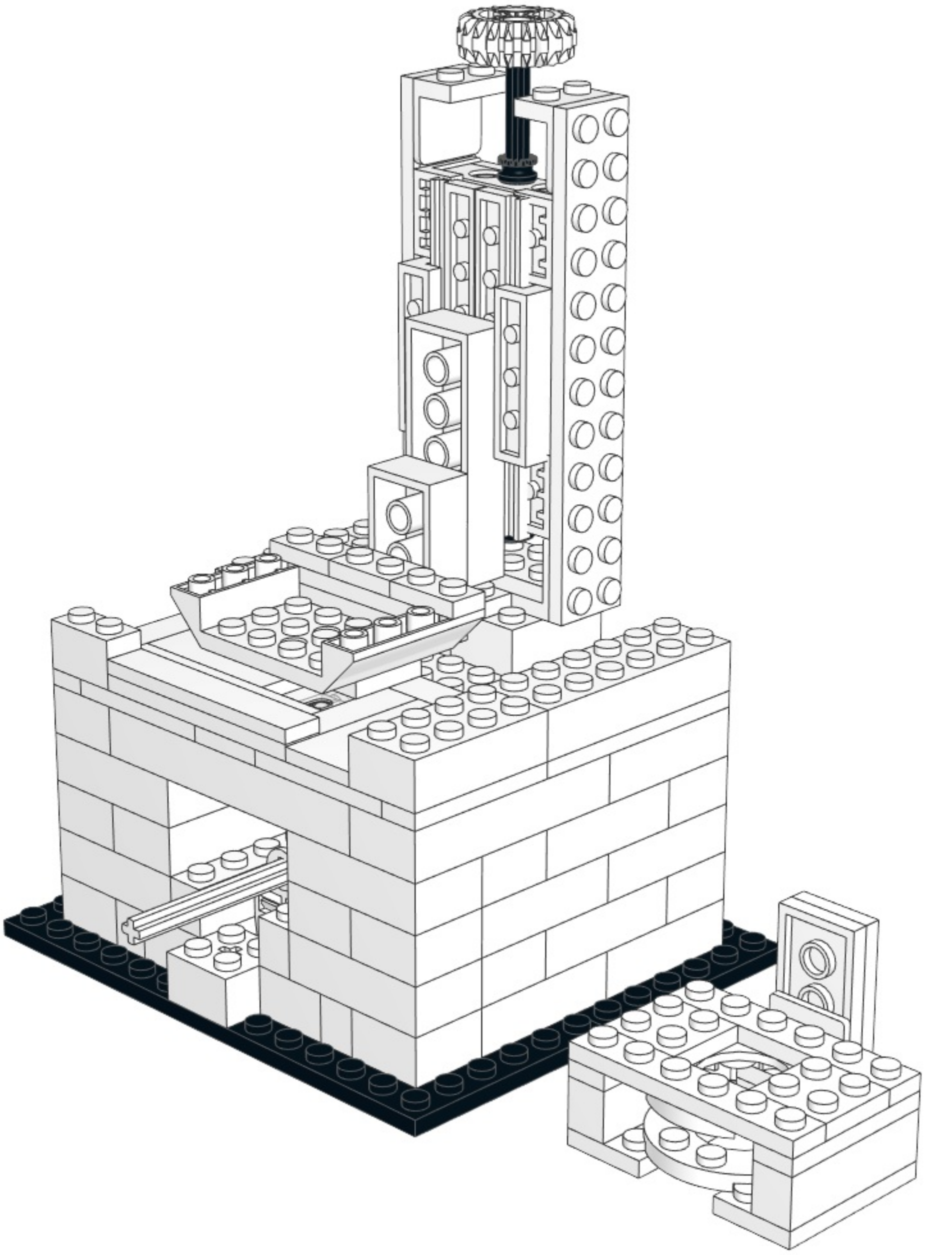

# 16

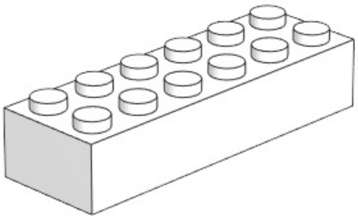

2x

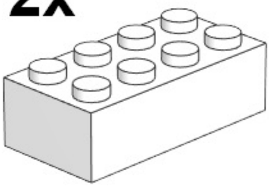

2x

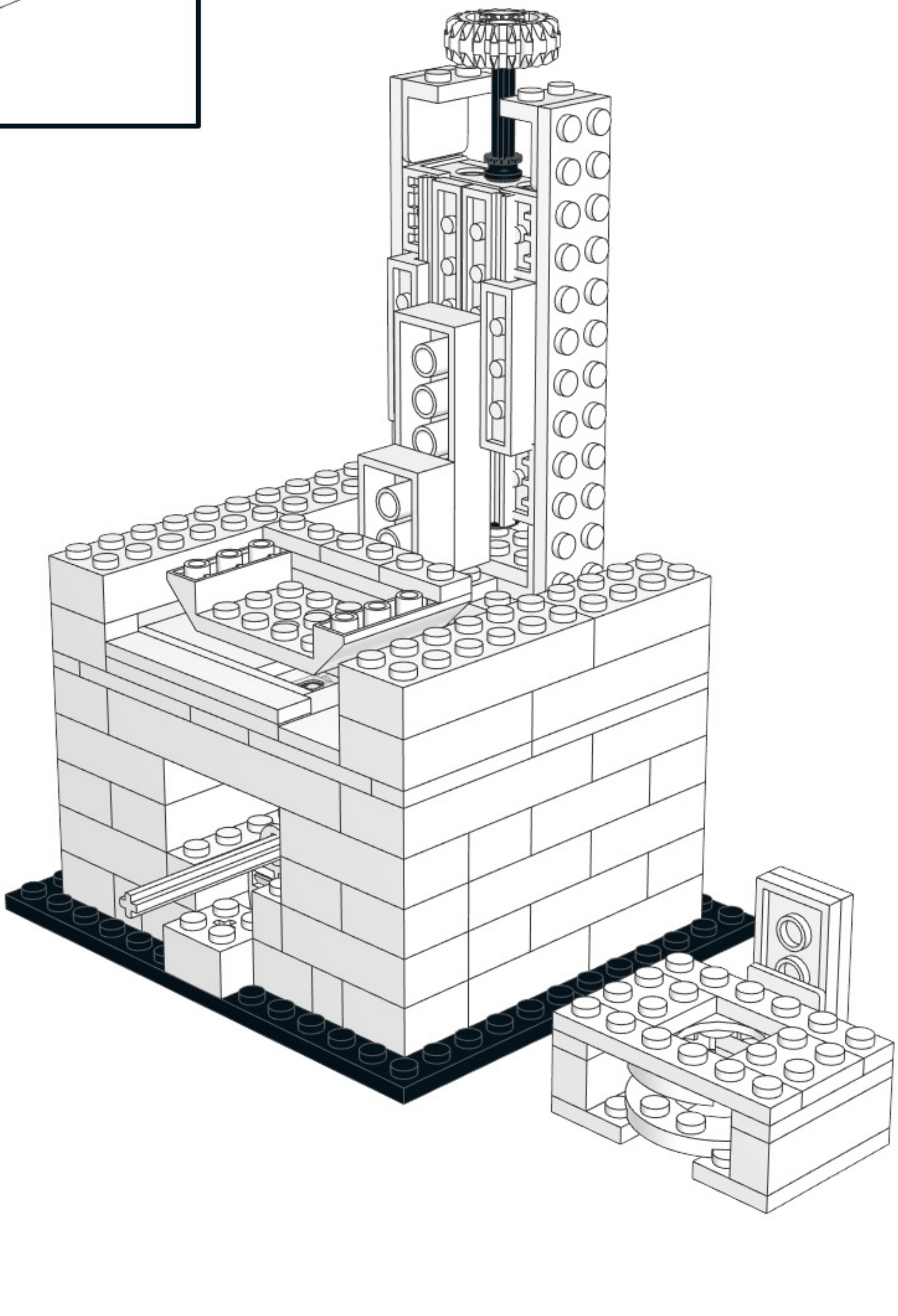

# 17

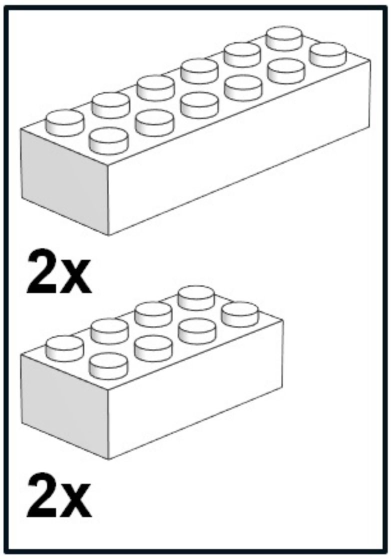


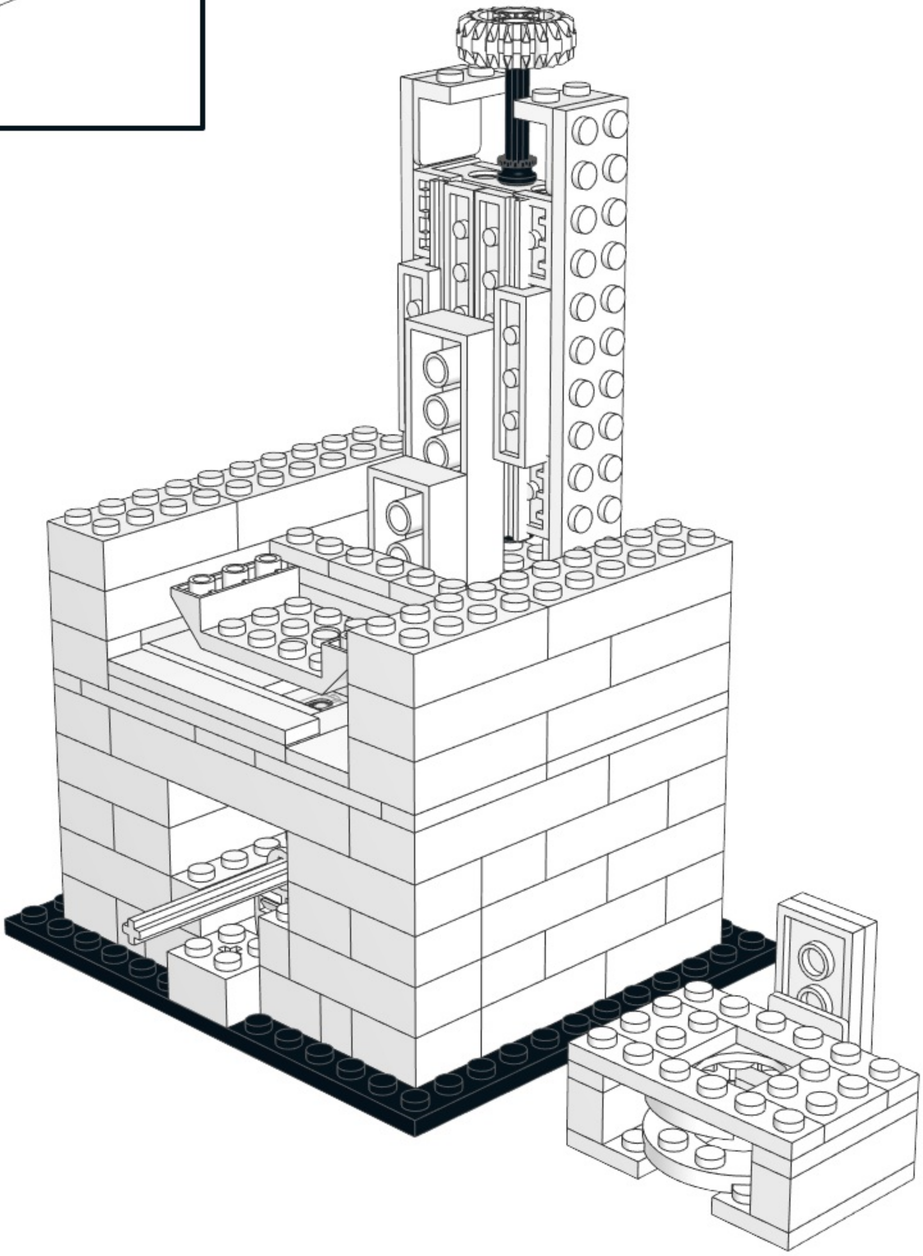

# 18

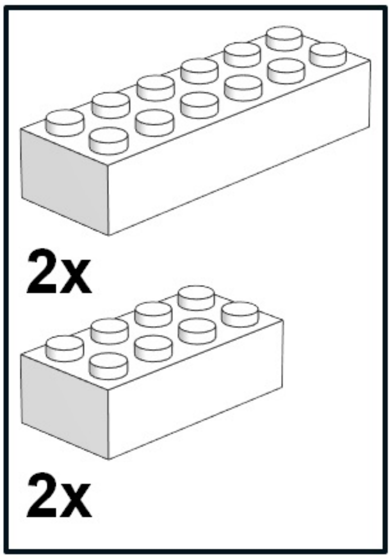


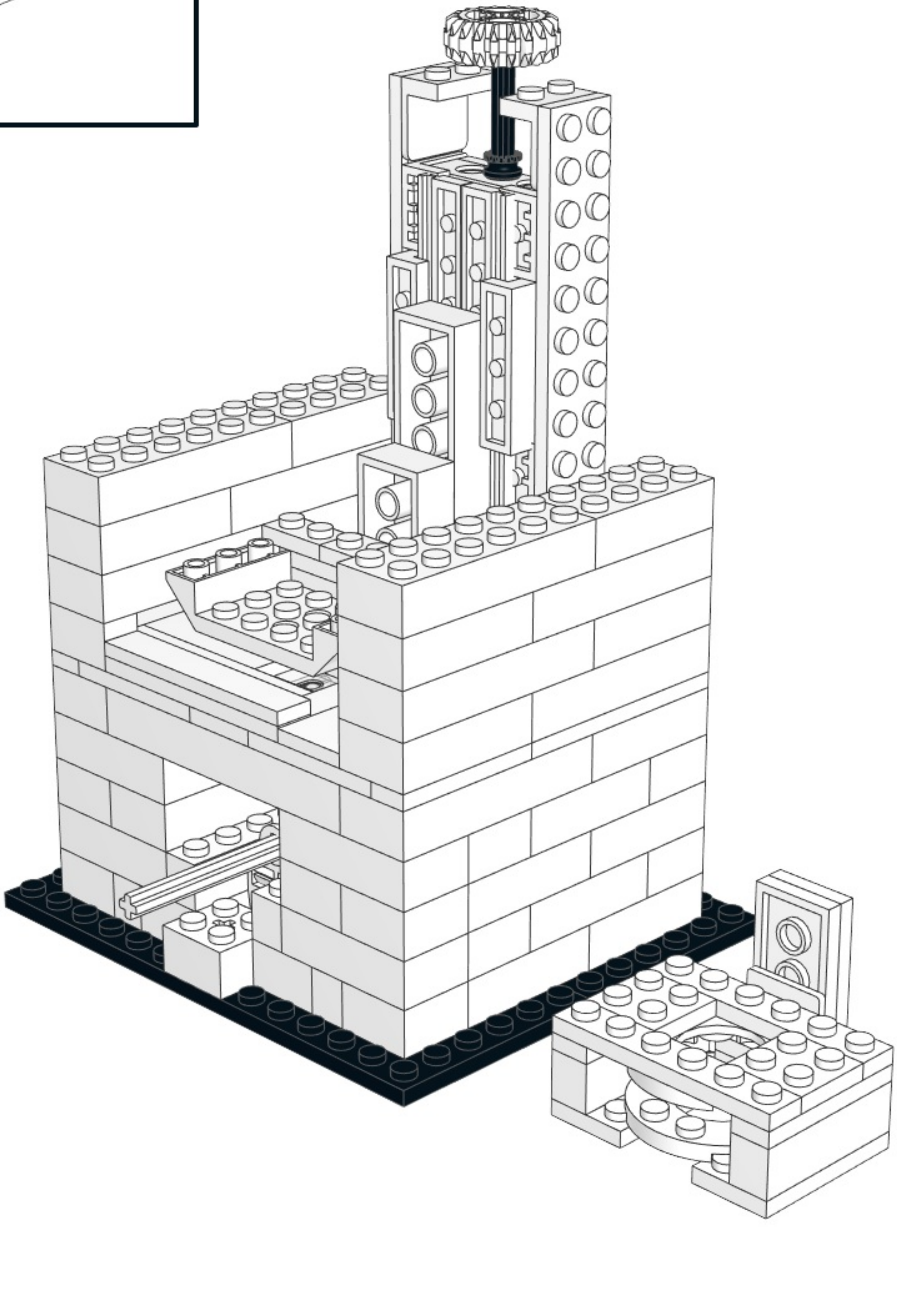

19

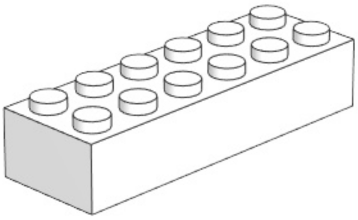

2x

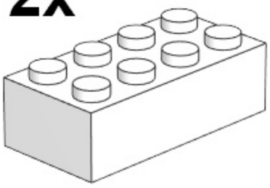

2x

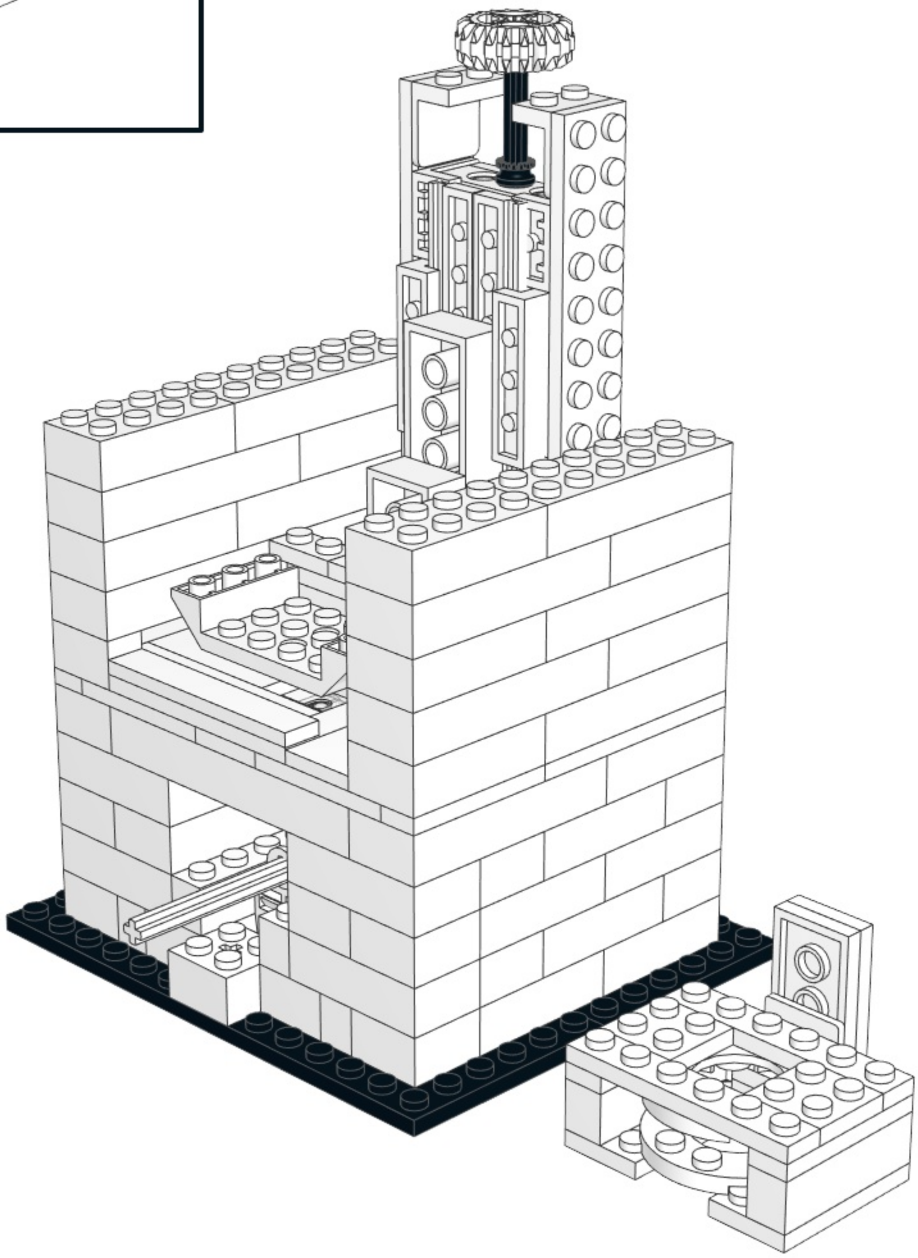

20

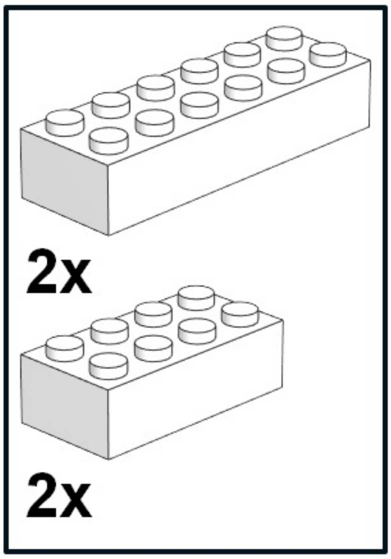


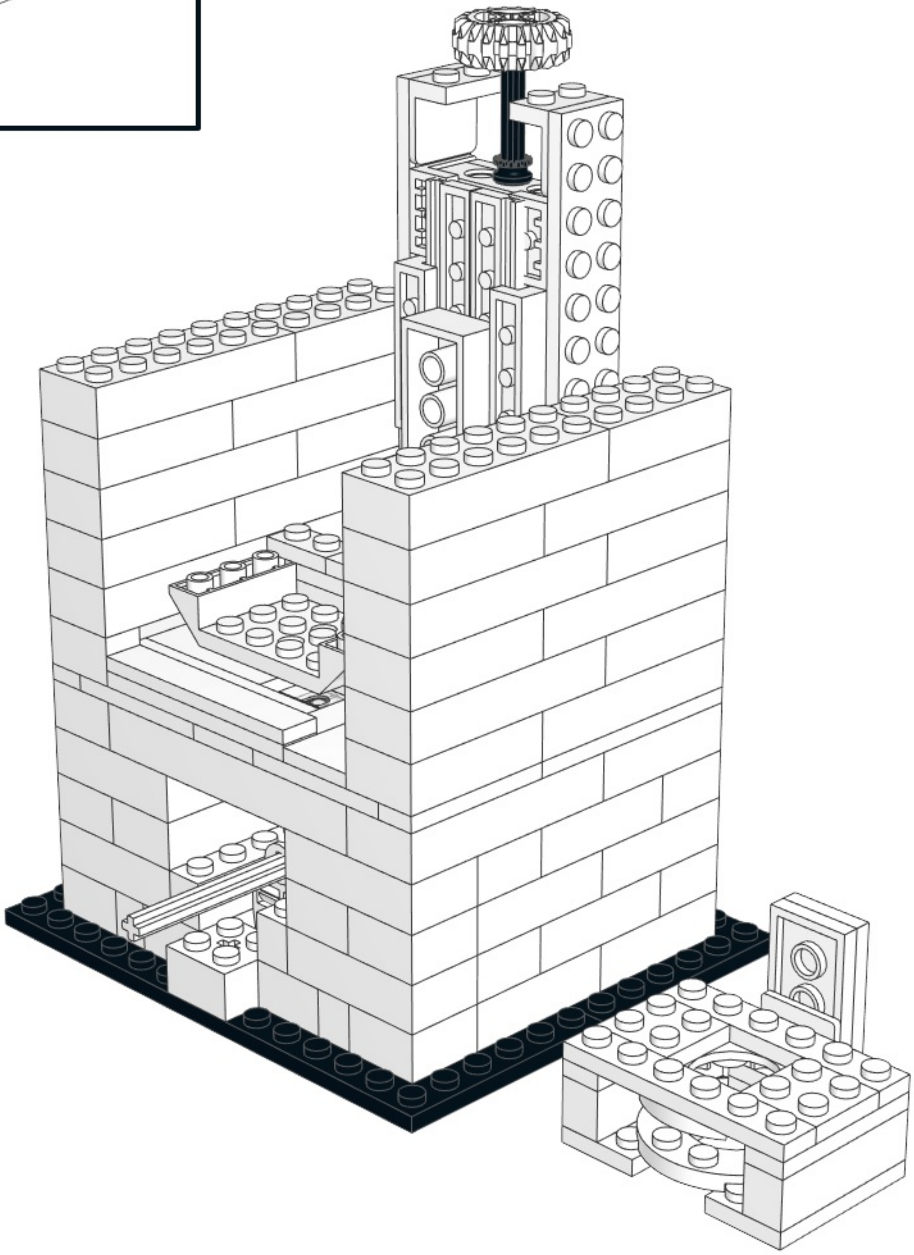

# 21

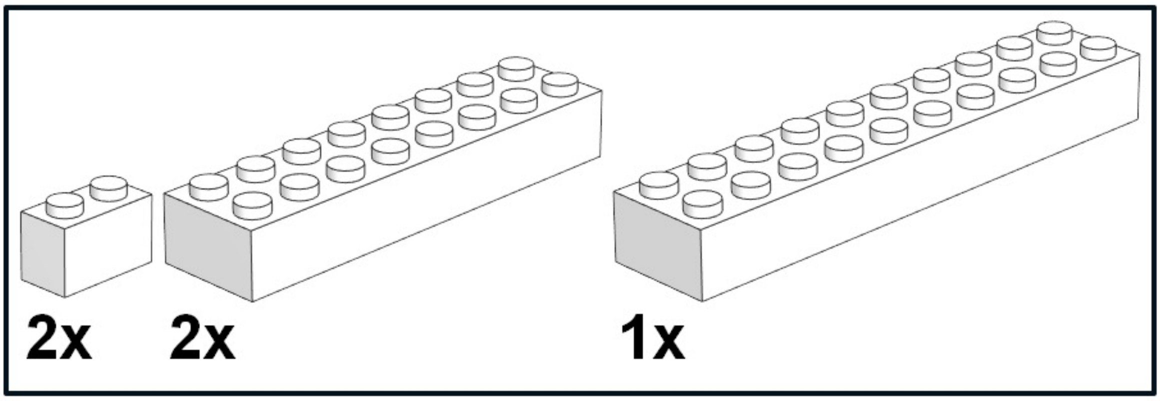


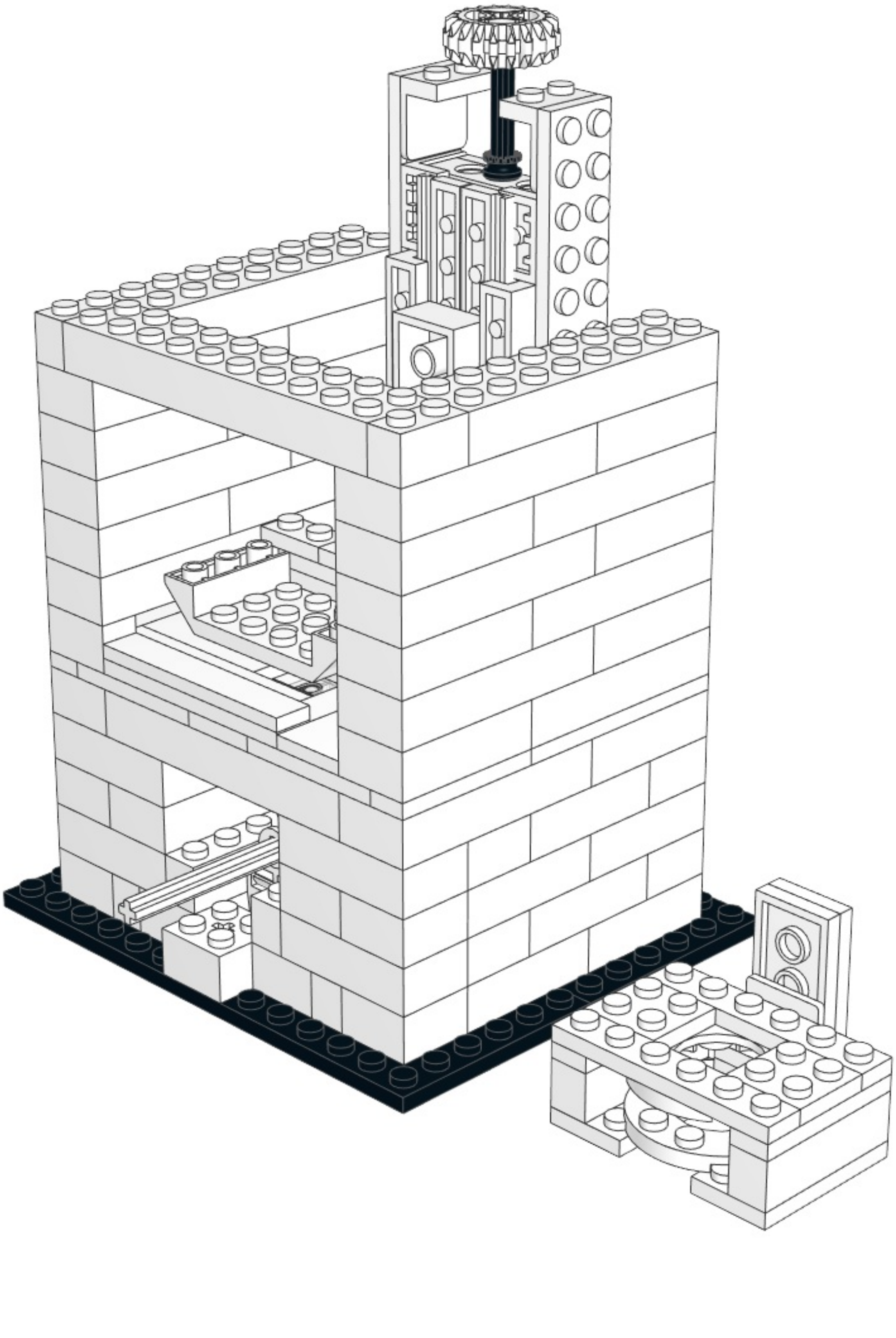

22

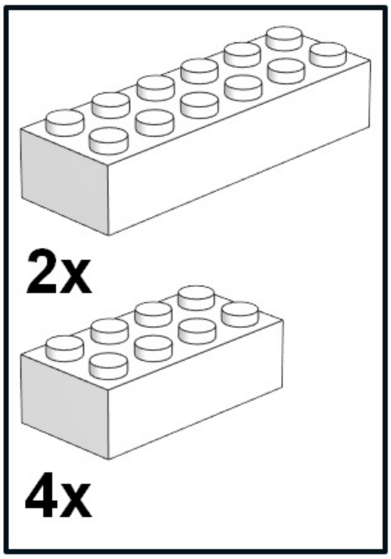


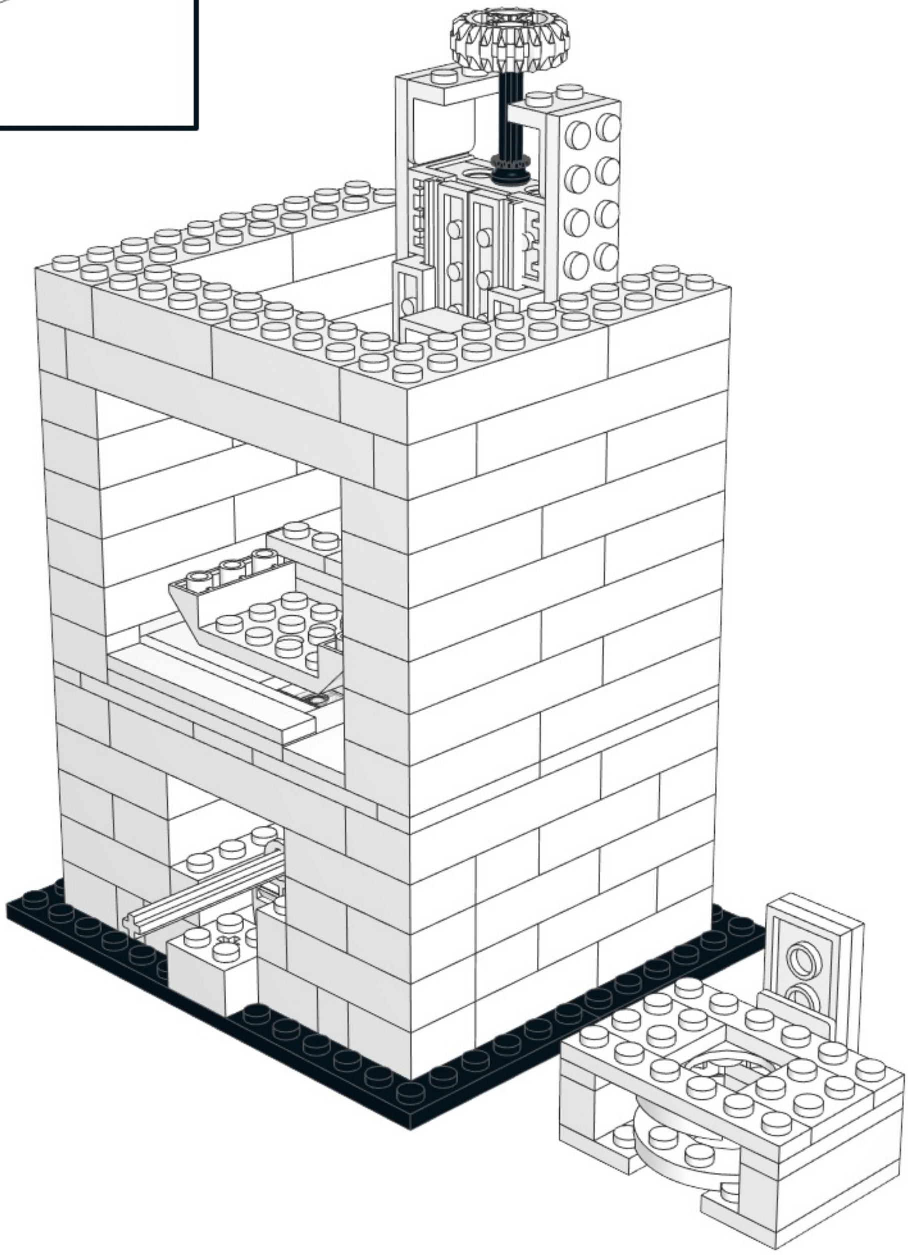

# 23

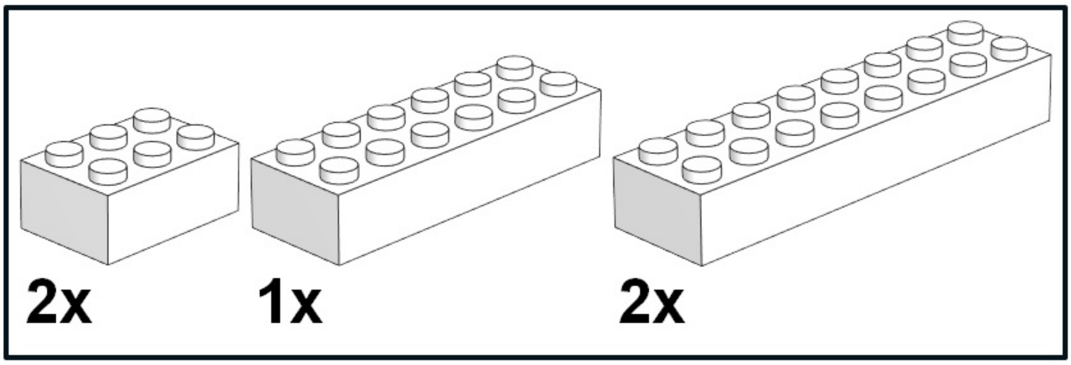


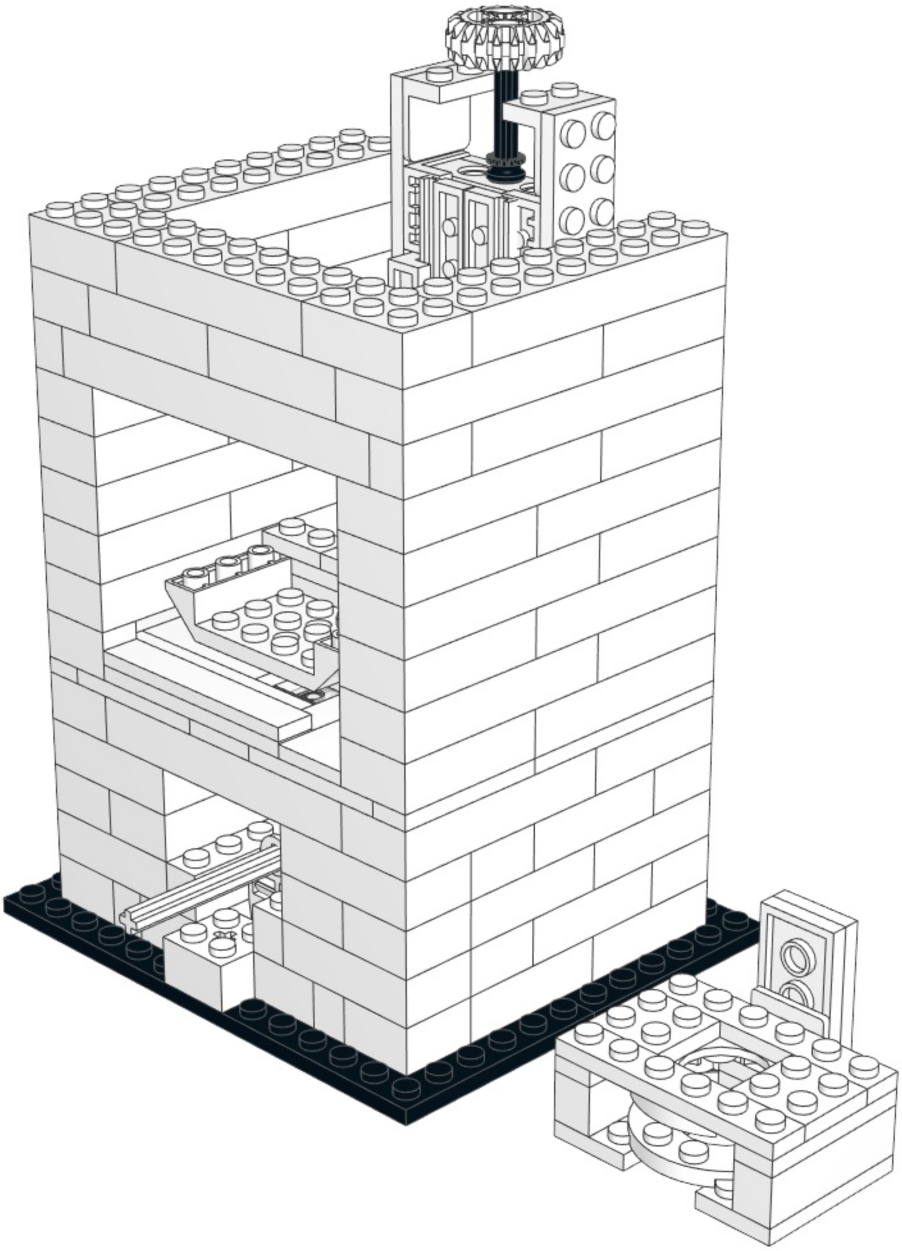

# 24

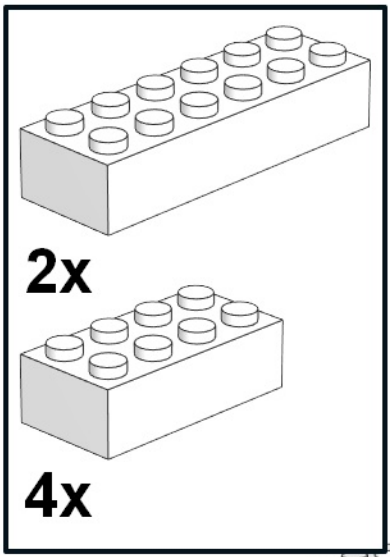


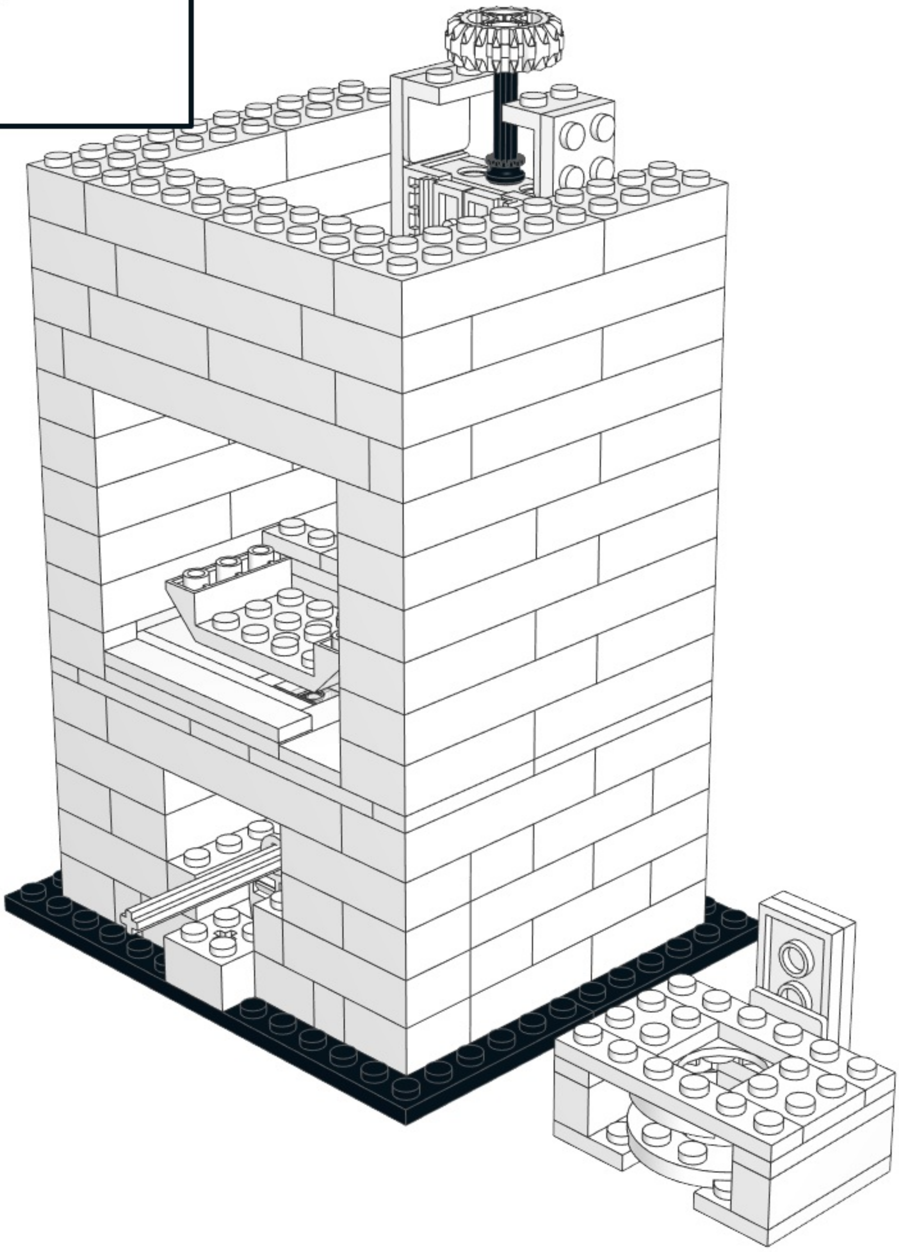

## 25

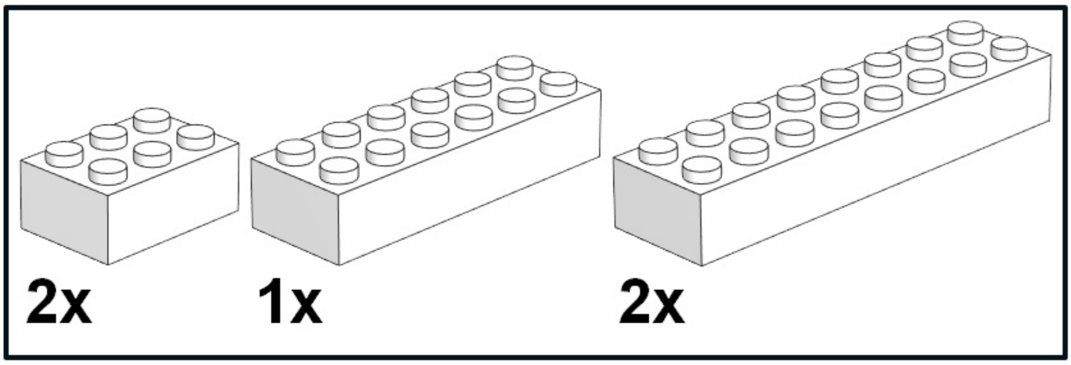


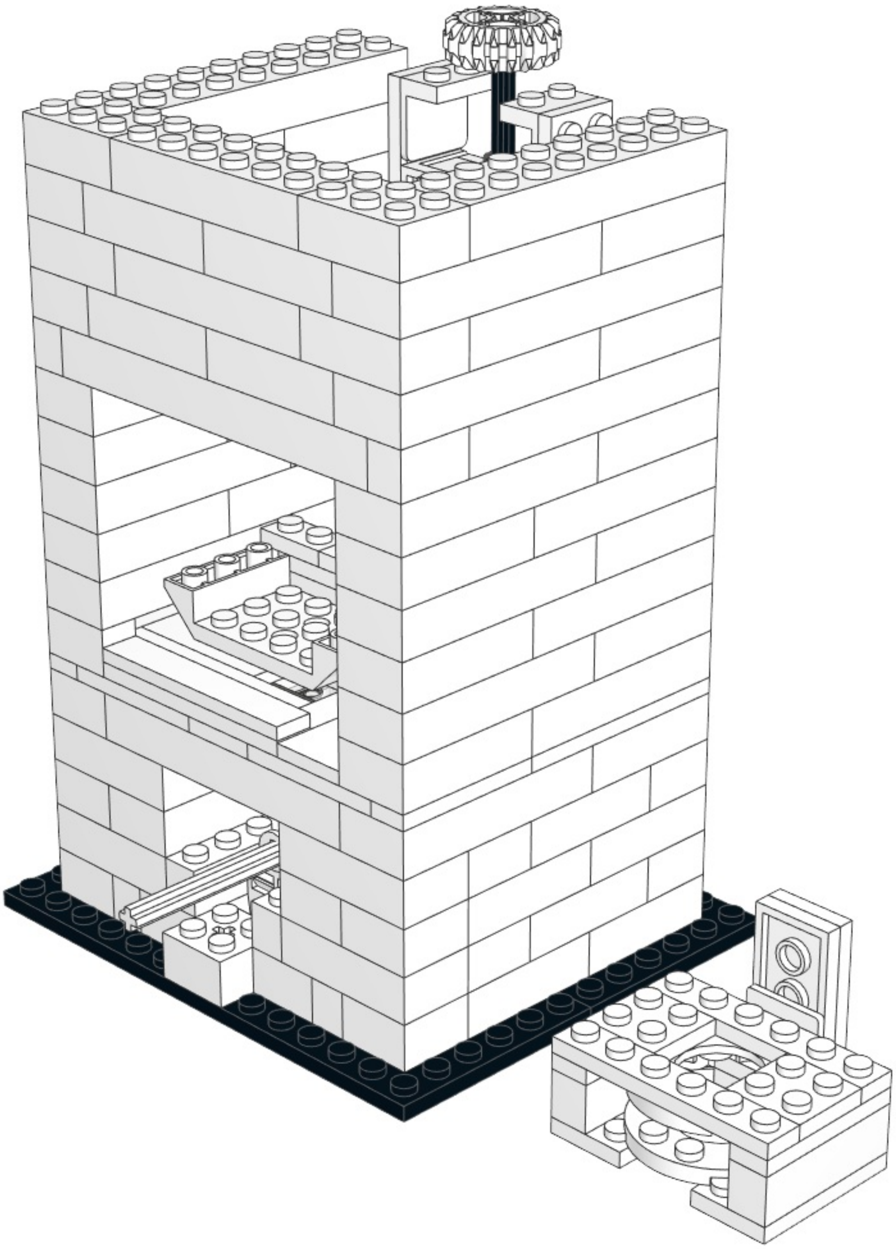

# 26

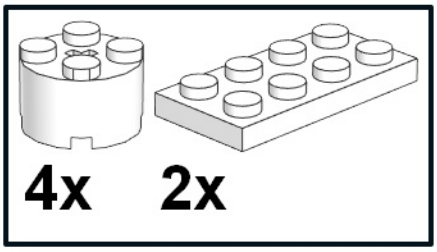


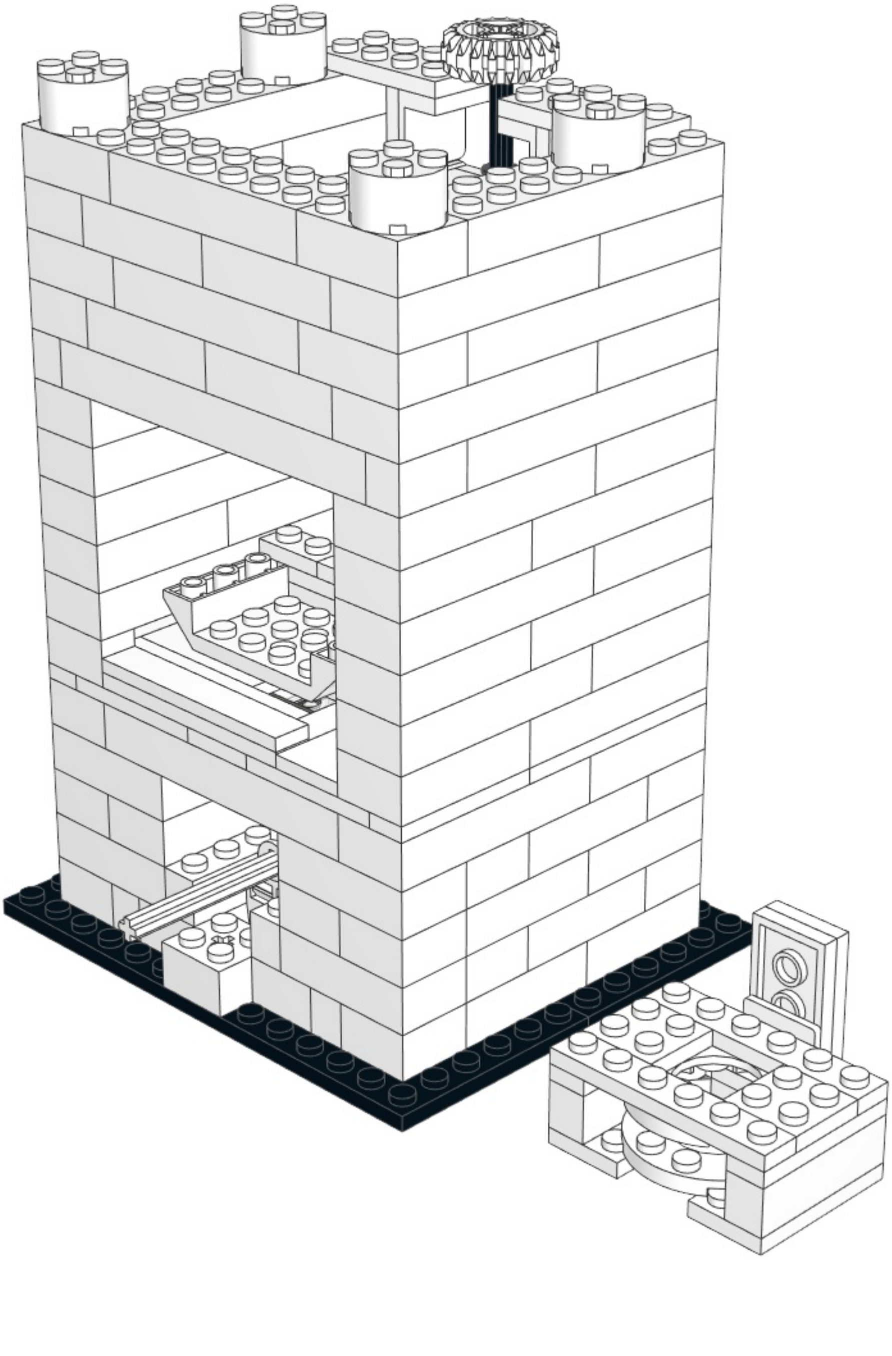

## 27

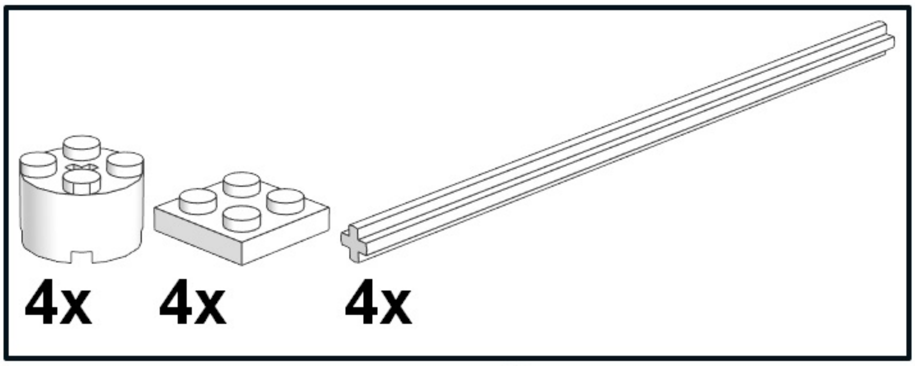


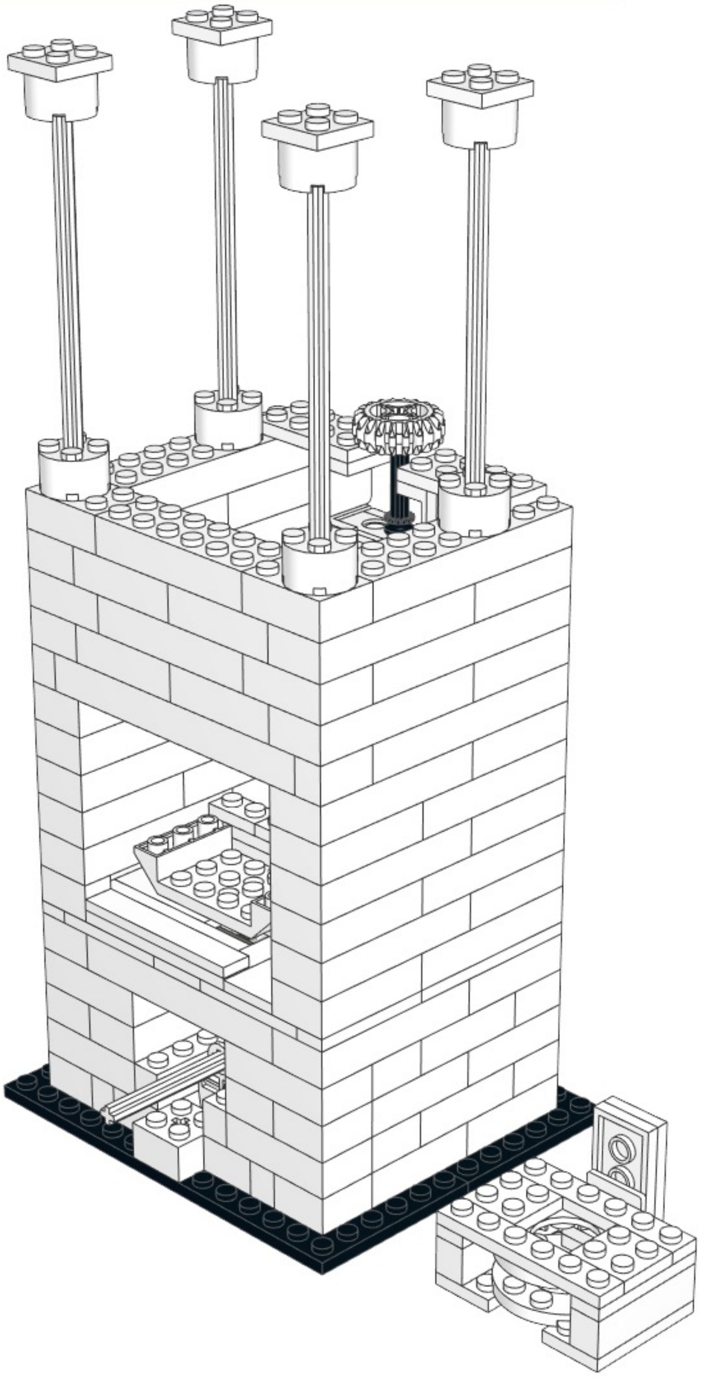

1

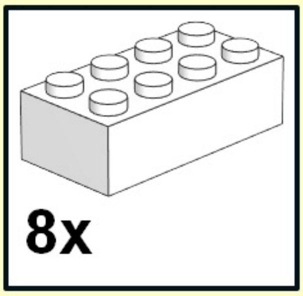


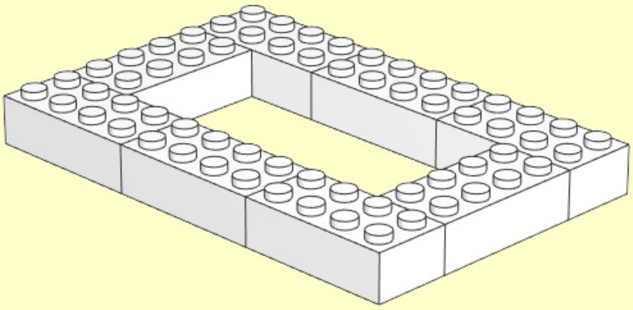

2

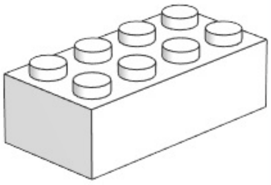

8x

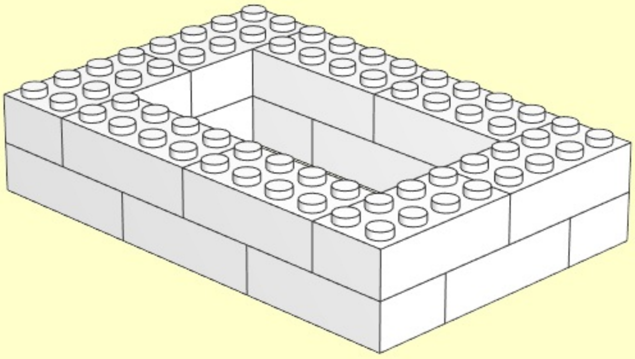

3

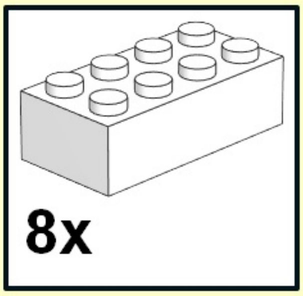


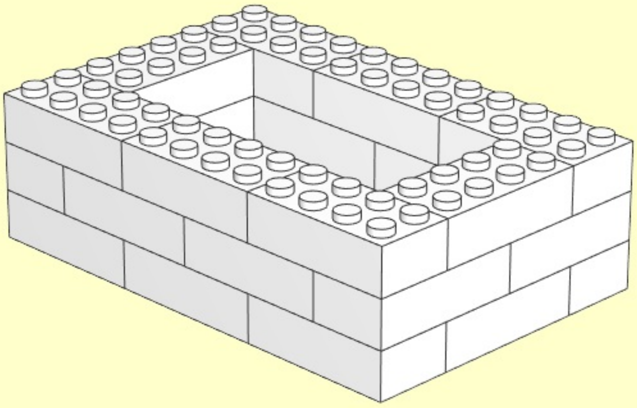

4

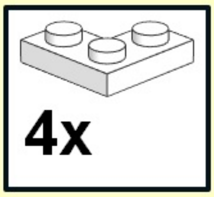


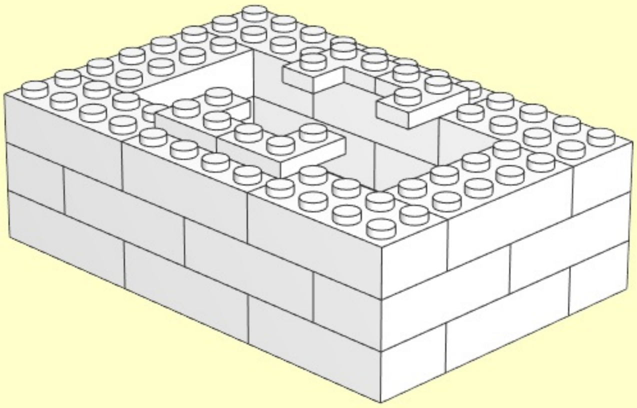

5

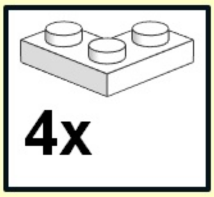


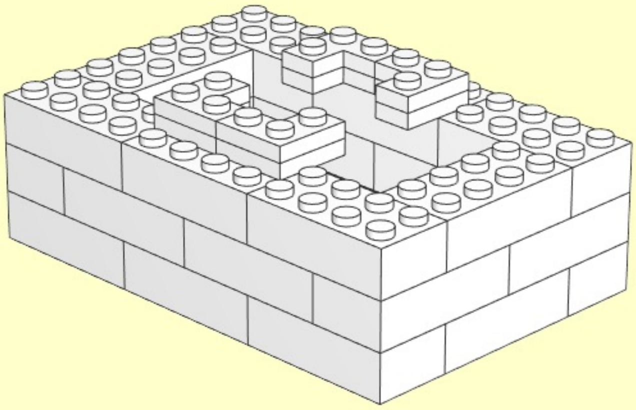

6

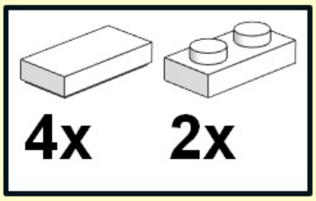


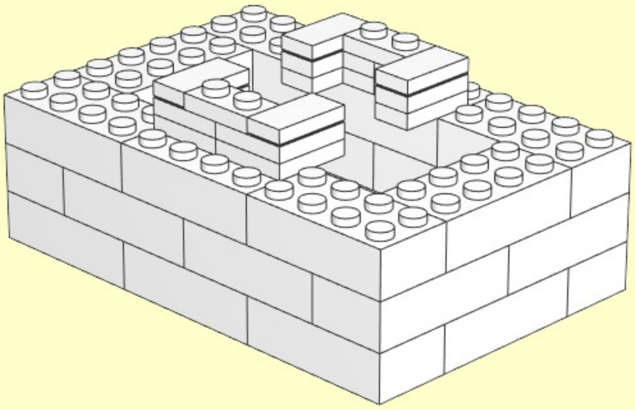

**7**

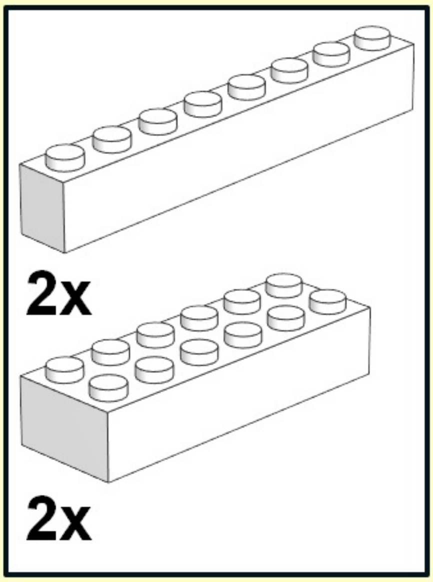


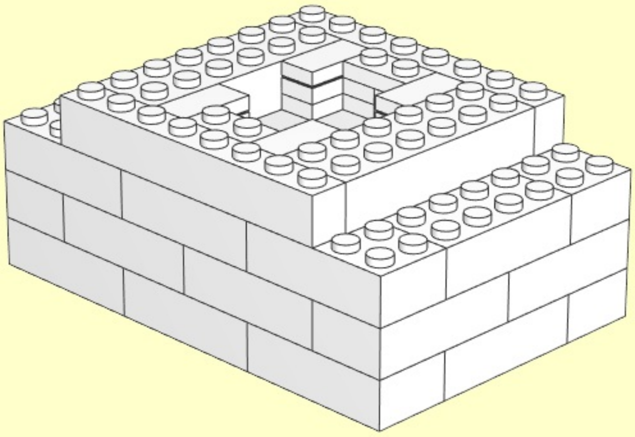

8

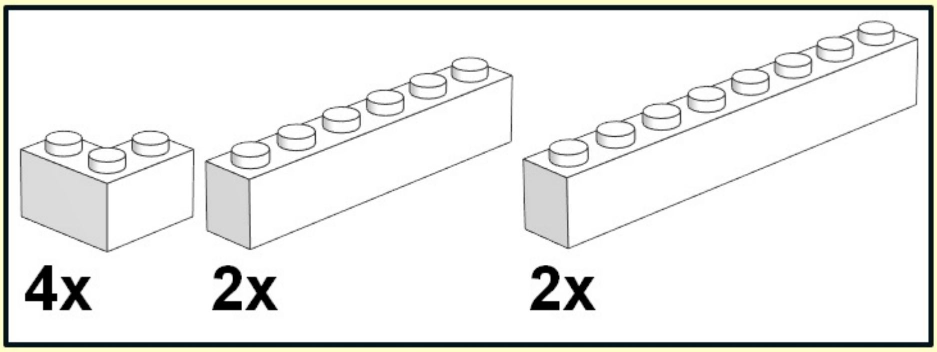


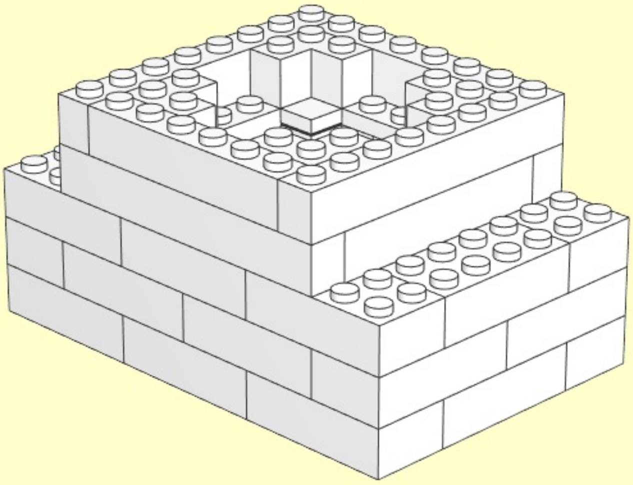

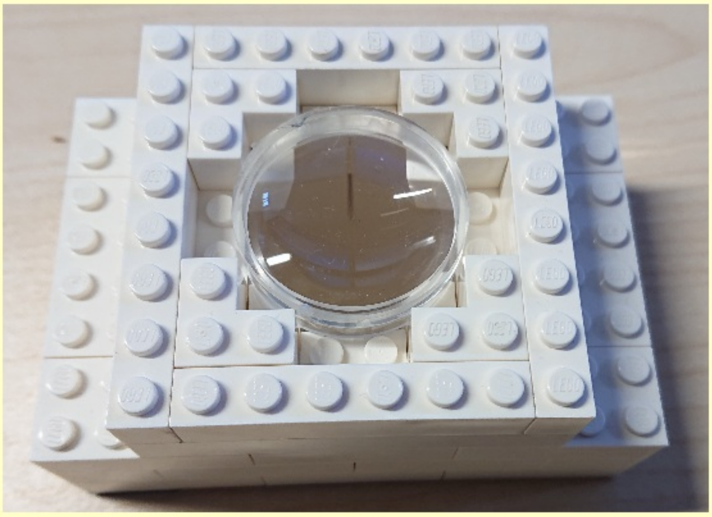

9

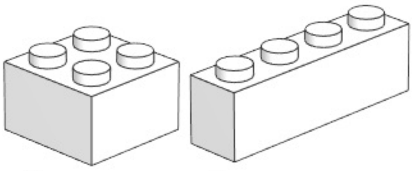

4x 4x

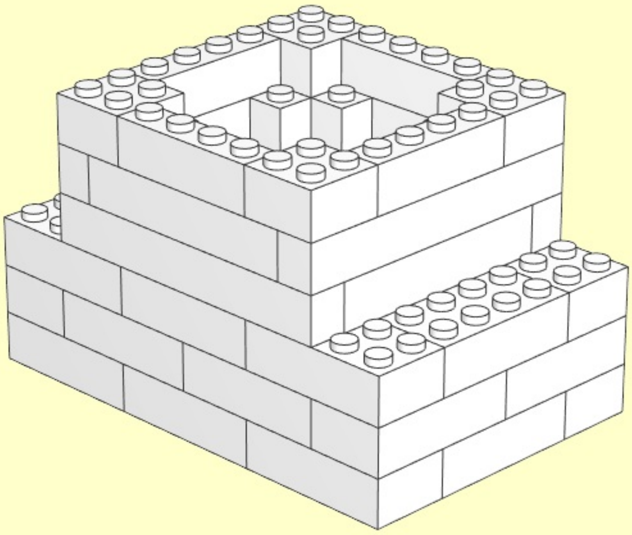

# 10

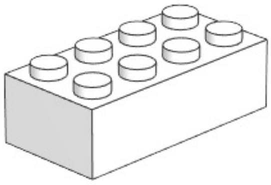

**6x**

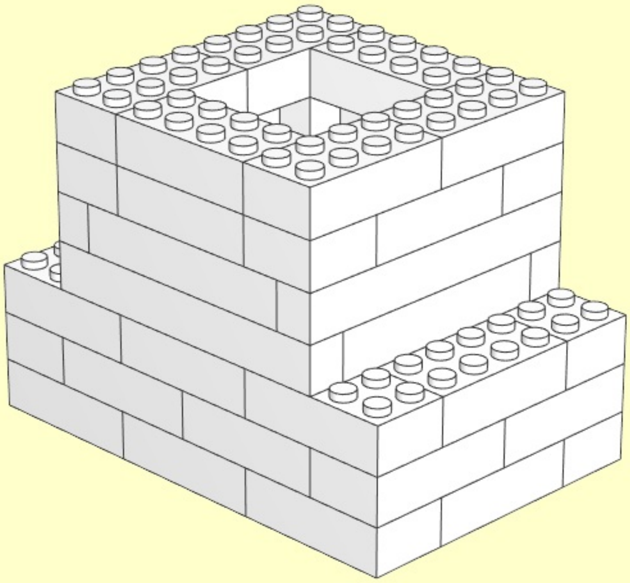

**11**

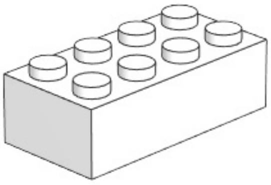

**6x**

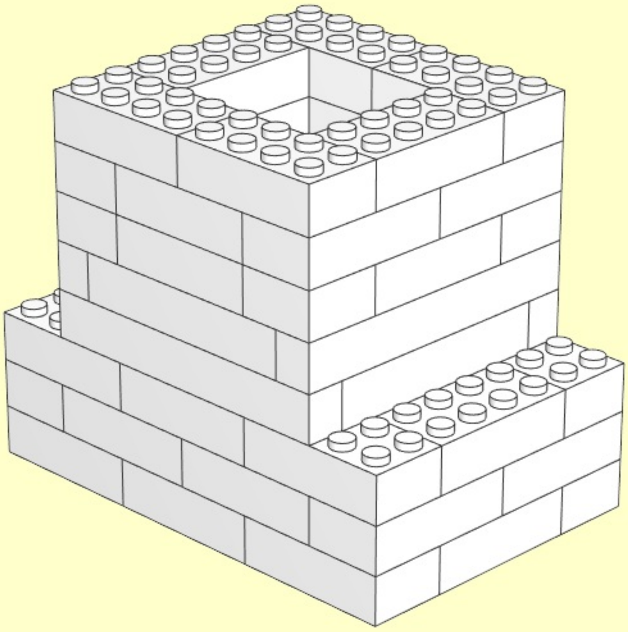

**12**

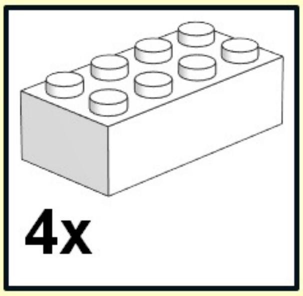


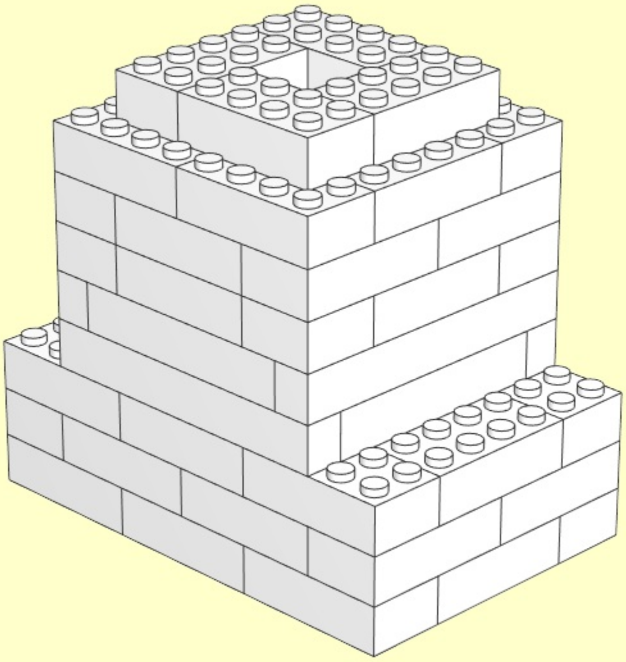

13

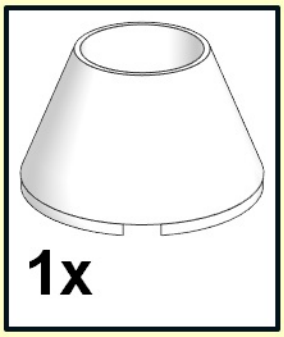


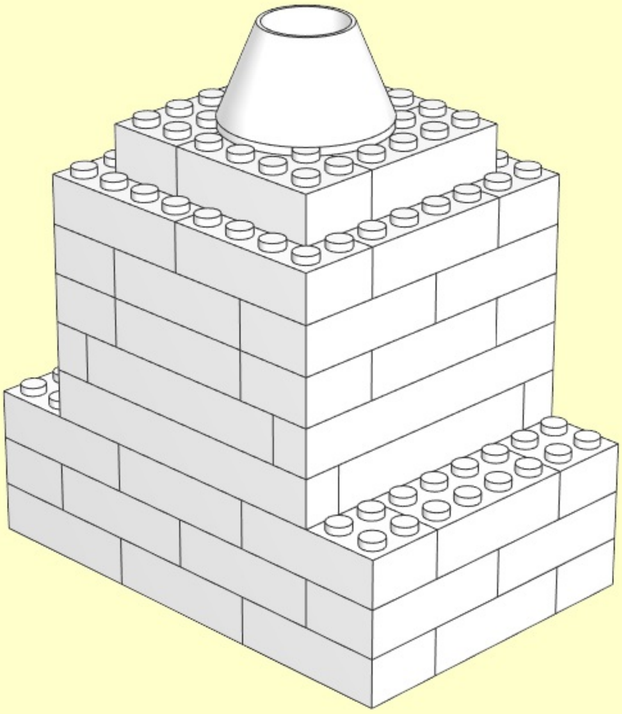

**14**

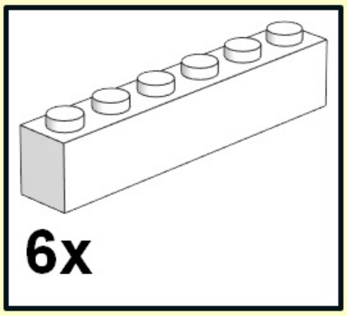


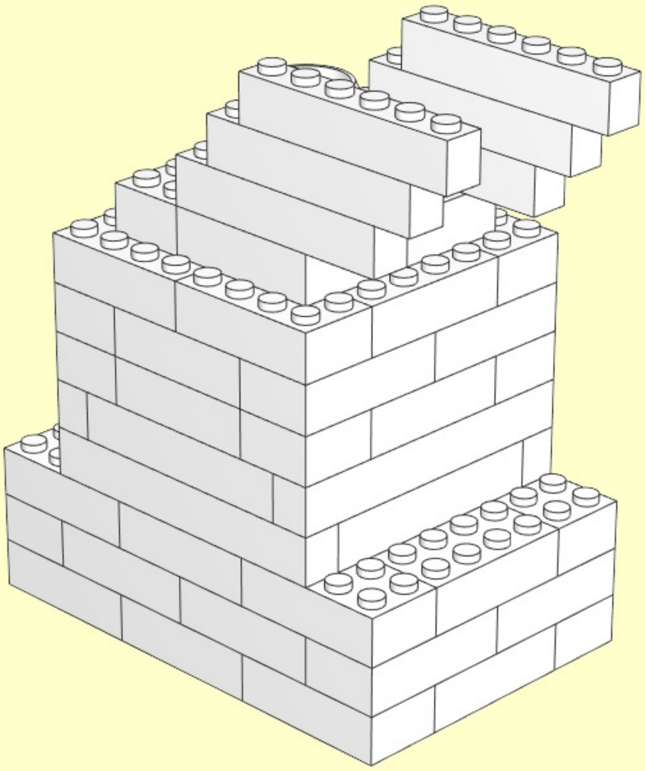

# 15

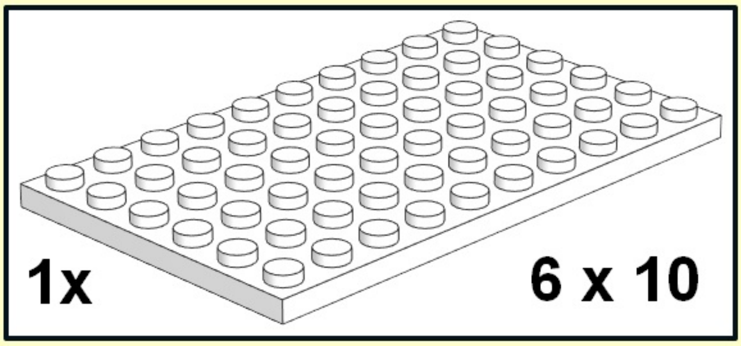


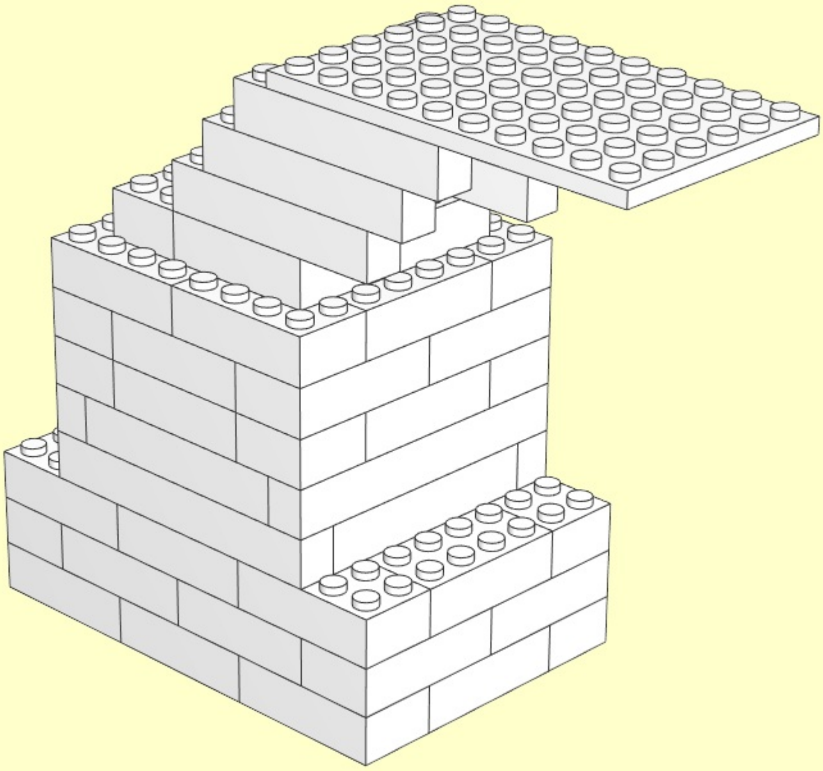

28

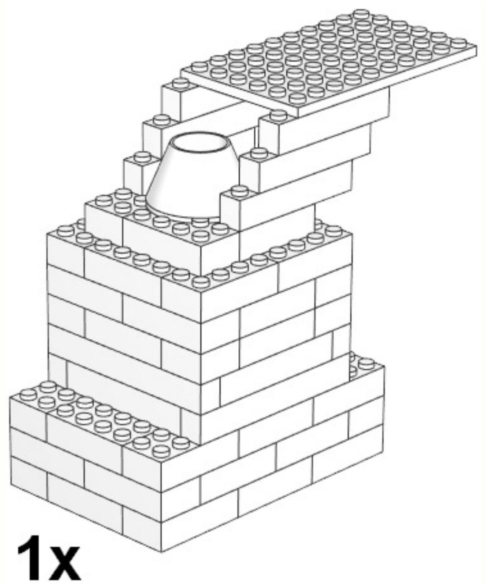

1x

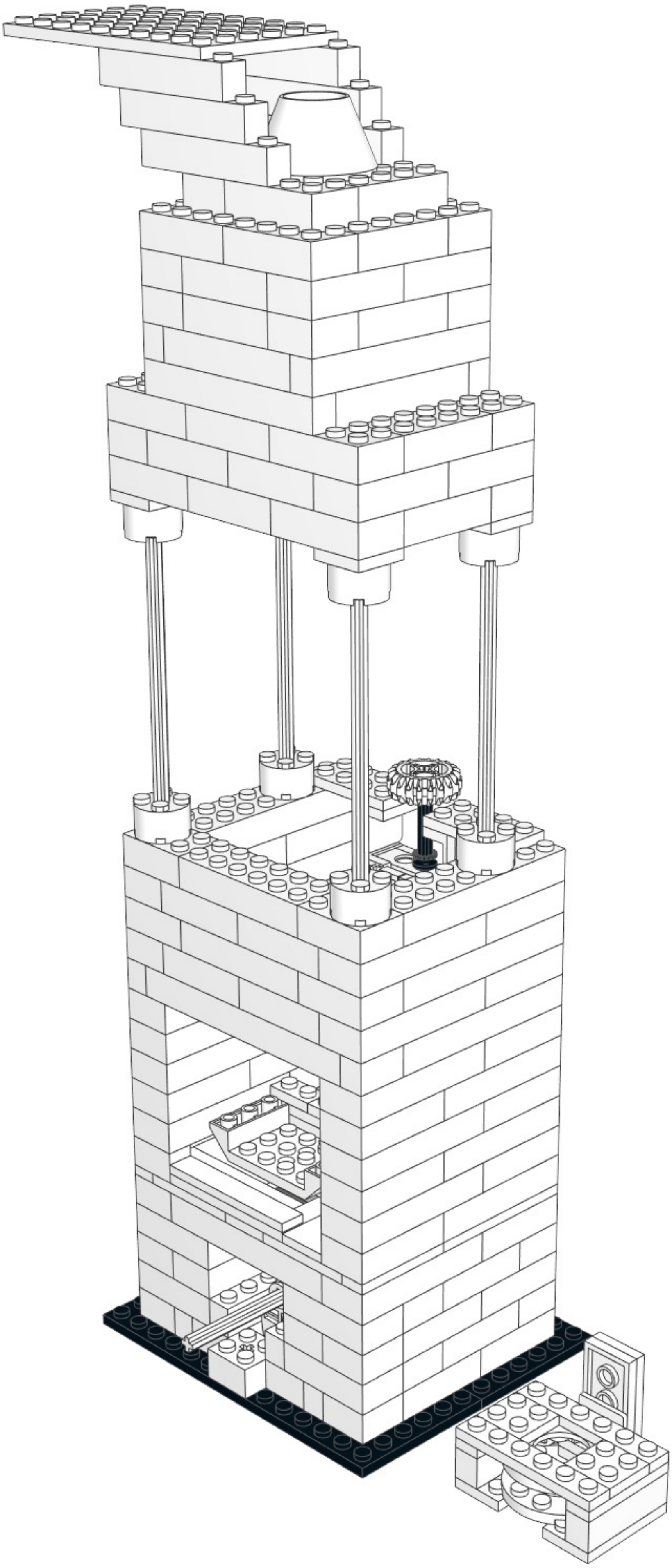

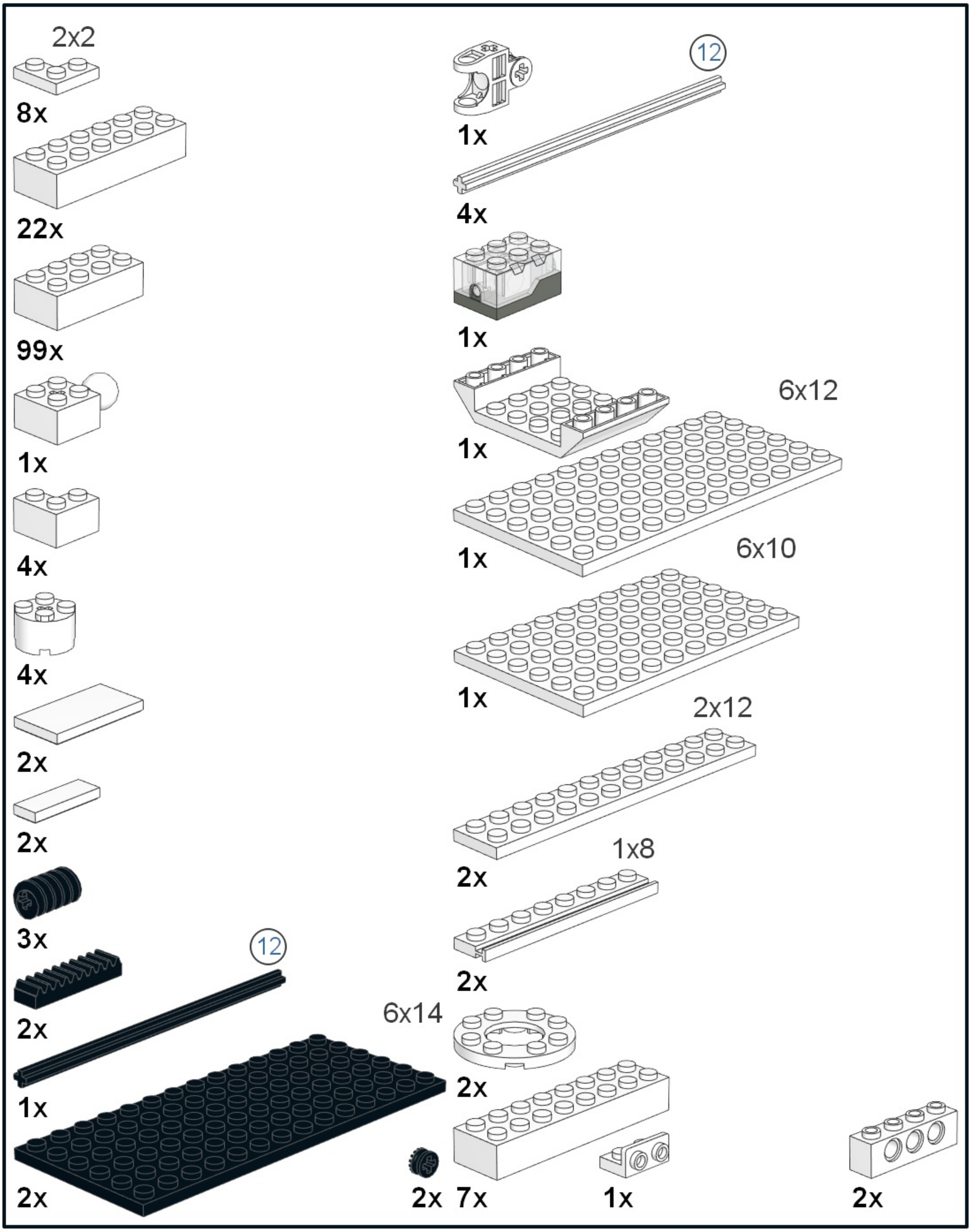
2x2
8x
22x
99x
1x
4x
4x
2x
2x
3x
12
2x
1x
6x14
2x
2x
12
1x
4x
1x
1x
6x12
1x
6x10
1x
2x12
2x
1x8
2x
2x
7x
1x
2x

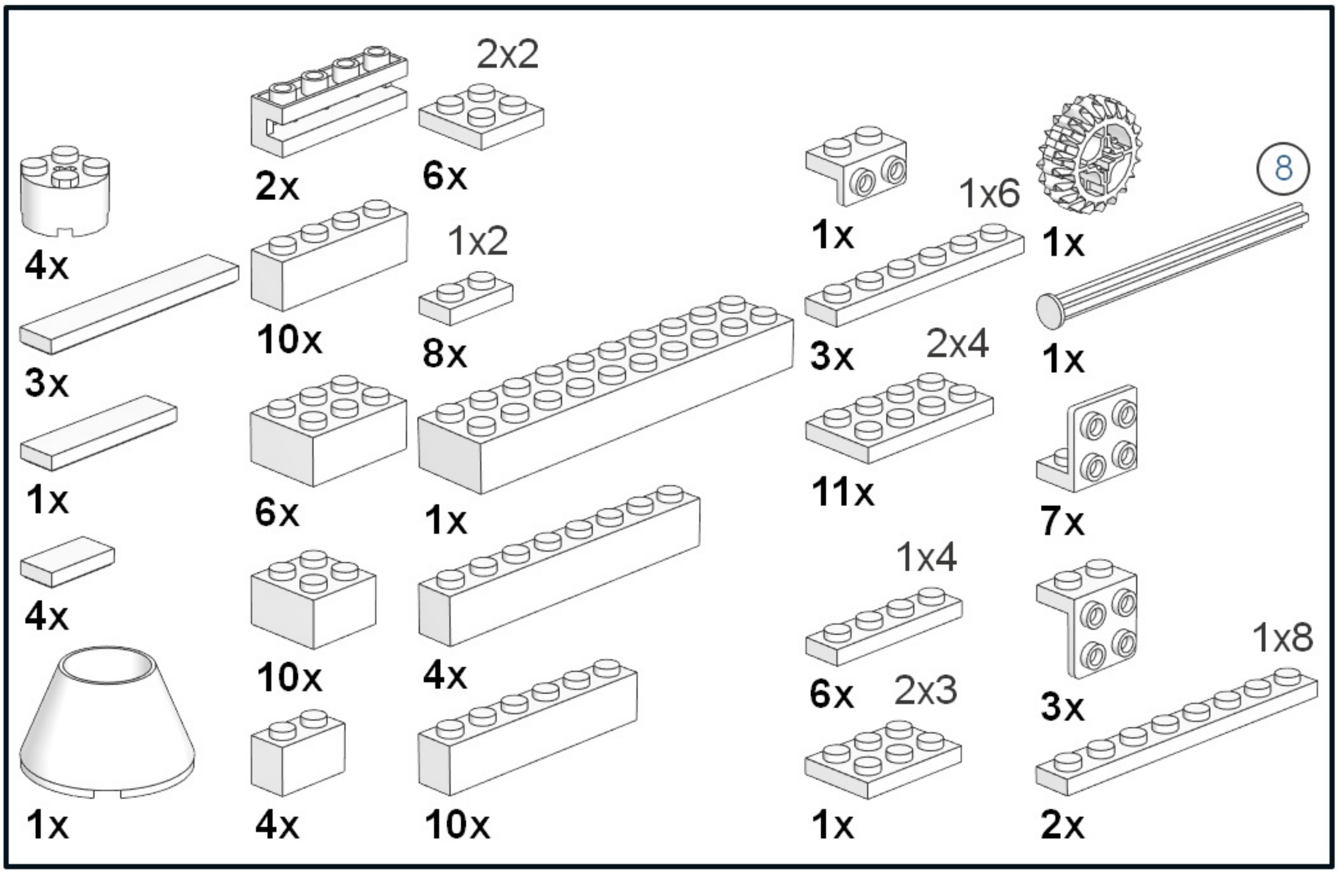
2x2
4x
2x
6x
1x
1x6
1x
8
3x
10x
1x2
8x
3x
1x
2x4
1x
6x
1x
11x
7x
4x
4x
1x4
10x
6x
2x3
3x
1x8
1x
4x
10x
1x
2x
