## Supplementary File 2 for "Designing a high-resolution, LEGO-based microscope for an educational setting"

### Parts List for Mikroscope_plan.mpd (287 parts)

| Part | Color | Quantity | Description |
|---|---|---|---|
| 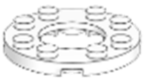 11833 | 15: White | 2 | Plate 4 x 4 Round with 2 x 2 Round Hole |
| 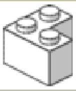 2357 | 15: White | 4 | Brick 2 x 2 Corner |
| 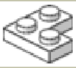 2420 | 15: White | 8 | Plate 2 x 2 Corner |
| 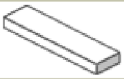 2431 | 15: White | 1 | Tile 1 x 4 with Groove |
| 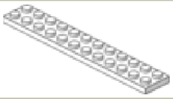 2445 | 15: White | 2 | Plate 2 x 12 |
| 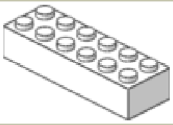 2456 | 15: White | 22 | Brick 2 x 6 |
| 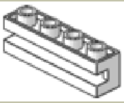 2653 | 15: White | 2 | Brick 1 x 4 with Groove |
| 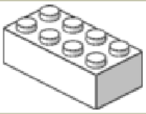 3001 | 15: White | 99 | Brick 2 x 4 |
| 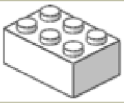 3002 | 15: White | 6 | Brick 2 x 3 |
| 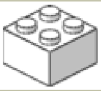 3003 | 15: White | 10 | Brick 2 x 2 |
| 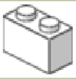 3004 | 15: White | 4 | Brick 1 x 2 |
| 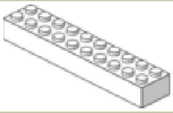 3006 | 15: White | 1 | Brick 2 x 10 |
| 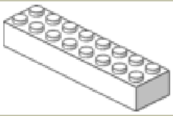 3007 | 15: White | 7 | Brick 2 x 8 |
| 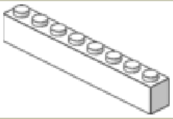 3008 | 15: White | 4 | Brick 1 x 8 |
| 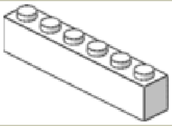 3009 | 15: White | 10 | Brick 1 x 6 |
| 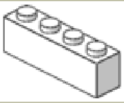 3010 | 15: White | 10 | Brick 1 x 4 |
| 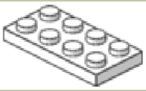 3020 | 15: White | 11 | Plate 2 x 4 |
| 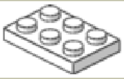 3021 | 15: White | 1 | Plate 2 x 3 |
| 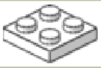 3022 | 15: White | 6 | Plate 2 x 2 |
| 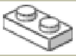 3023 | 15: White | 8 | Plate 1 x 2 |
| 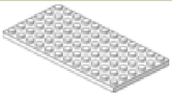 3028 | 15: White | 1 | Plate 6 x 12 |

| | | | | |
|---|---|---|---|---|
| 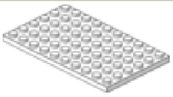 | 3033 | 15: White | 1 | Plate 6 x 10 |
| 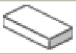 | 3069b | 15: White | 4 | Tile 1 x 2 with Groove |
| 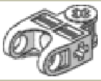 | 32174 | 15: White | 1 | Constraction Connector 3 x 2 with Single Round Ball Socket |
| 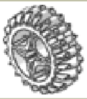 | 32269 | 15: White | 1 | Technic Gear 20 Tooth Double Bevel |
| 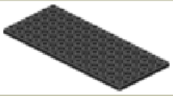 | 3456 | 0: Black | 2 | Plate 6 x 14 |
| 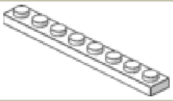 | 3460 | 15: White | 2 | Plate 1 x 8 |
| 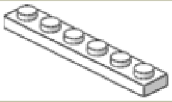 | 3666 | 15: White | 3 | Plate 1 x 6 |
| 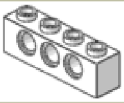 | 3701 | 15: White | 2 | Technic Brick 1 x 4 with Holes |
| 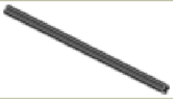 | 3708 | 0: Black | 1 | Technic Axle 12 |
| 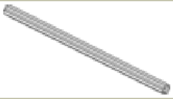 | 3708 | 15: White | 4 | Technic Axle 12 |
| 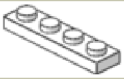 | 3710 | 15: White | 6 | Plate 1 x 4 |
| 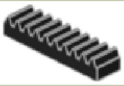 | 3743 | 0: Black | 2 | Technic Gear Rack 1 x 4 |
| 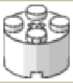 | 3941 | 15: White | 4 | Brick 2 x 2 Round |
| 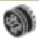 | 4265a | 0: Black | 2 | Technic Bush 1/2 Type 1 |
| 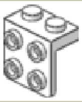 | 44728 | 15: White | 3 | Bracket 1 x 2 - 2 x 2 |
| 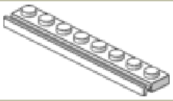 | 4510 | 15: White | 2 | Plate 1 x 8 with Door Rail |
| 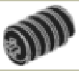 | 4716 | 0: Black | 3 | Technic Worm Gear |
| 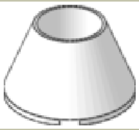 | 4742 | 15: White | 1 | Cone 4 x 4 x 2 Hollow No Studs |
| 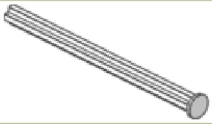 | 55013 | 15: White | 1 | Technic Axle 8 with Stop |
| 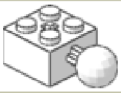 | 57909 | 15: White | 1 | Brick 2 x 2 with Ball Joint and Axlehole |
| 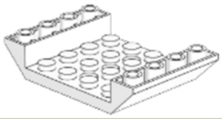 | 60219 | 15: White | 1 | Slope Brick 45 6 x 4 Double Inverted with Centre Holes |
| 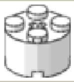 | 6143 | 15: White | 4 | Brick 2 x 2 Round Type 2 |
| 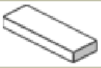 | 63864 | 15: White | 2 | Tile 1 x 3 with Groove |

| | | | | | |
|---|---|---|---|---|---|
| 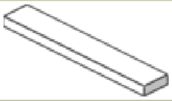 | 6636 | 15: White | | 3 | Tile 1 x 6 |
| 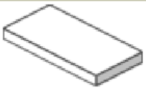 | 87079 | 15: White | | 2 | Tile 2 x 4 with Groove |
| 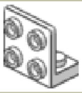 | 99207 | 15: White | | 7 | Bracket 1 x 2 - 2 x 2 Up |
| 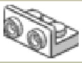 | 99780 | 15: White | | 1 | Bracket 1 x 2 - 1 x 2 Up |
| 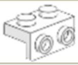 | 99781 | 15: White | | 1 | Bracket 1 x 2 - 1 x 2 Down |
| 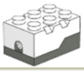 | u9158 | 15: White | | 1 | Electric Light Brick 2 x 3 x 1.333 |

This parts list was generated by LDView.
