## Supplementary File 3 for "Designing a high-resolution, LEGO-based microscope for an educational setting"

### Dear children, dear adults,

As scientists we regularly get many good questions from people of all ages. Of course we notice with how much enthusiasm you want to learn more about various phenomena of nature around us. Even though we could answer many of those questions directly, nothing goes above looking for the answer yourself. Of course the internet can be used while searching for the answer, or you can look it up in an encyclopedia, but you really learn something by trying to find the answer yourself by closely observing and thinking about experiments. That is also exactly what it means to be a scientist. Although it seems that scientists always know everything, what drives us are the gaps in knowledge. Despite the extraordinary advances that our understanding of the world has made over the past few centuries, there is still an infinite number of secrets to unravel. Of course you need a multitude of instruments to exploring nature, and in this kit we want to build one together with you. We will not tell you yet what it will be, because you should find out yourself. But maybe you already have an idea?

The instructions of this kit consist of multiple parts. There is always a basic task, often you have to build something out of Lego, think about something, or try to answer a question that we ask you. To make sure that you have understood everything that is important, we always give you hints that you can use... to avoid missing something important!

To test whether and what you have learned, you can fill in a short questionnaire. Please fill in questionnaire A before you start with the kit. Just answer what you think is the correct answer. Many questions are a little difficult on purpose because children of different ages fill out the same form. It is important that after you have done all the tasks, and perhaps also played a little, you do the second questionnaire (B). You can then compare what you did differently in questionnaire B. In that way, you can determine what you have learned. The solutions to the questions can be found online.

P.S. Some parts are very complicated to build! It’s best to let an adult do that. This holds for Object 1 and 2.

But enough talking, let’s start building something with Lego. We start on the next page.

1) Build Plan A, B and C from the LEGO bricks. In one part you have to include the plastic lens. With a picture we tried to show how that goes. We hope that the plans are clear, but we're sure that you LEGO-experts can do it easily.

2) Explore the 3 resulting LEGO-objects.

Now look at each of the finished objects from all sides. Maybe you can already do something with it?

If you want, you can write down here what you can do with the different parts.

A)

B)

C)

Now you can compare your solution with the solution on page 12.

3) Now take a closer look at part B. What can you do with it? What are the problems?
(There are hints on page 12)

4) How can we solve the problem of illumination?
(There are hints on page 13)

5) Take a look at some objects around you. Maybe a fly’s wing or a part of a plant? You can also try to magnify some text with the magnifying glass.

What happens when you change the position of your head or the distance from the magnifying glass to the object you’re looking at? (There are hints on page 13)

And an additional question: when you look at a text, are the letters normal, or are they inverted?

6) To summarize the findings so far:
   a. With A and B we can make a good magnifying glass, that even has illumination built in.
   b. Changing the distances between the object you're looking at, the lens and your eye changes the magnification.
   c. A larger distance between the magnifying glass and the object increases the magnification, but you also need to move your eye further away.

7) Now let's think about it. Unfortunately, we cannot infinitely increase the distance between the magnifying glass and your eye, to increase the magnification. At some point it just doesn't work anymore. Do you have an idea what we can do to get a higher magnification?
   (There are hints on page 13)

8) Now get object (1), that an adult built for you.

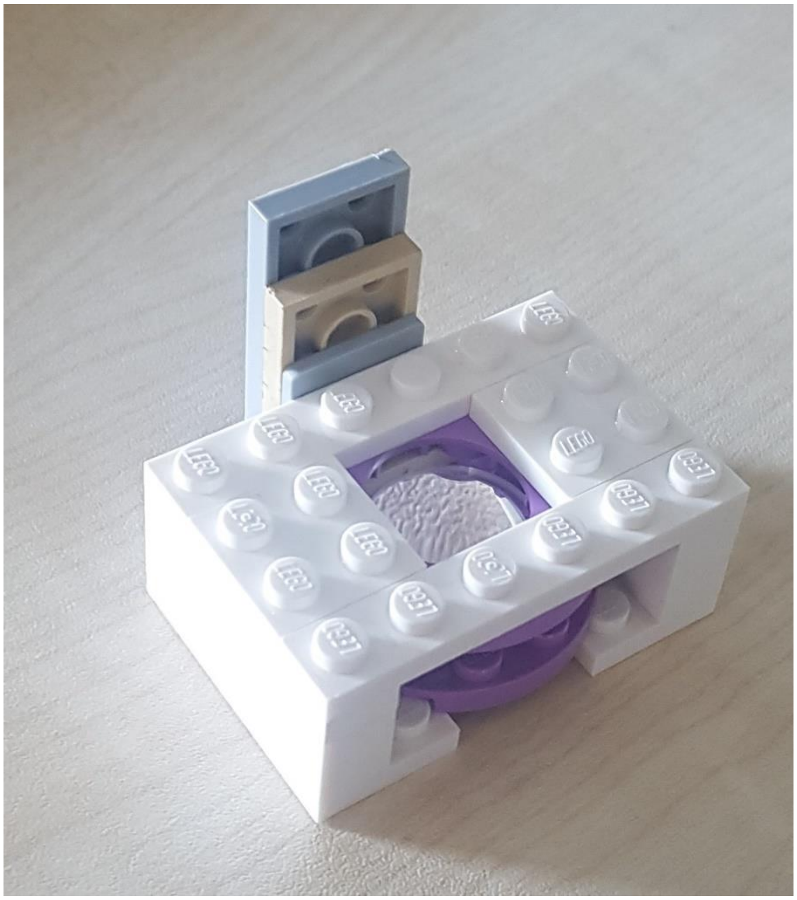

As you can see, this is again a lens built into Lego. As its diameter is smaller than that of the magnifying glass, we will call this the “small magnifying glass”. Let’s pursue the idea of making a better magnifying glass by combining two magnifying glasses. If we build the large magnifying glass on the light source, we can try to enlarge the large magnifying glass again with the small one. Give it a try.

No worries, this is really hard…

In fact, it just doesn’t work. Maybe you notice that you can actually only see an enlarged image of the large magnifying glass.

Perhaps it’s better to do it the other way around, that is, hold the small magnifying glass about 3 to 4 cm away from the object you want to look at, and then hold the large magnifying glass close to your eye. Now carefully move the large magnifying glass towards the small magnifying glass until the image becomes sharp.

Does it work? What do you see? What is difficult? What problems do you have?

Now you can try to carefully move away from the small magnifying glass with the large magnifying glass.

What do you see? It may well be that you also have to change the distance of the small magnifying glass here. Just move slowly and always try to keep the image sharp.

9) To summarize the findings so far:
   a) Great magnification if you take two magnifying glasses
   b) Very sensitive to distance and position
   c) If you increase the distance between the object and the first magnifying glass, the magnification increases!

Do you have an idea how we can solve the problem that the positions are so sensitive?

The suggested solution is to combine all the parts using Lego. It’s getting really difficult now. If you get stuck, you can have a look at the help for question 9 on page 13.

10) Congratulations! You just built a real microscope out of Lego. Even if you can hardly believe that. You have all the essential parts in your LEGO microscope that we scientists also have in the laboratory. Only your microscope is much better for playing with ☺.

11) And that is exactly what you should do now. Play a little with the microscope. We wrote some suggestions on the website and made some videos that show you what you can prepare yourself. Take a look at the hints on page 16 for some ideas.

    What do you notice when you put a text under the microscope?

    What do you have to do to make the images nice and sharp?

12) Do you have any ideas how we can improve the magnification even further? Try to use object (2).

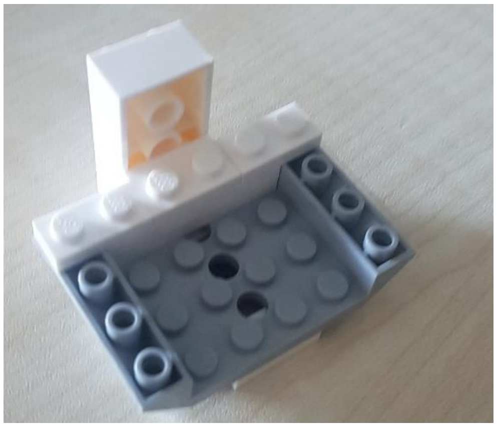

We call this the objective, because it is really like a microscope objective. By the way, the lens comes from the camera of an old iPhone 5.

Try to use the objective as a magnifying glass. Does it work? If so, what is difficult, if not, why not?

But maybe it will work better in the microscope if we remove the small magnifying glass and replace it with the objective?

Take a look at the hints on page 16 for some ideas.

13) Can we make the microscope compatible with a smartphone in order to take images and movies? (If you get stuck, look at the help on page 17)

14) Below you can see a drawing of a classic microscope. What are the similarities with our Lego microscope, which parts of the microscope have we reproduced where?

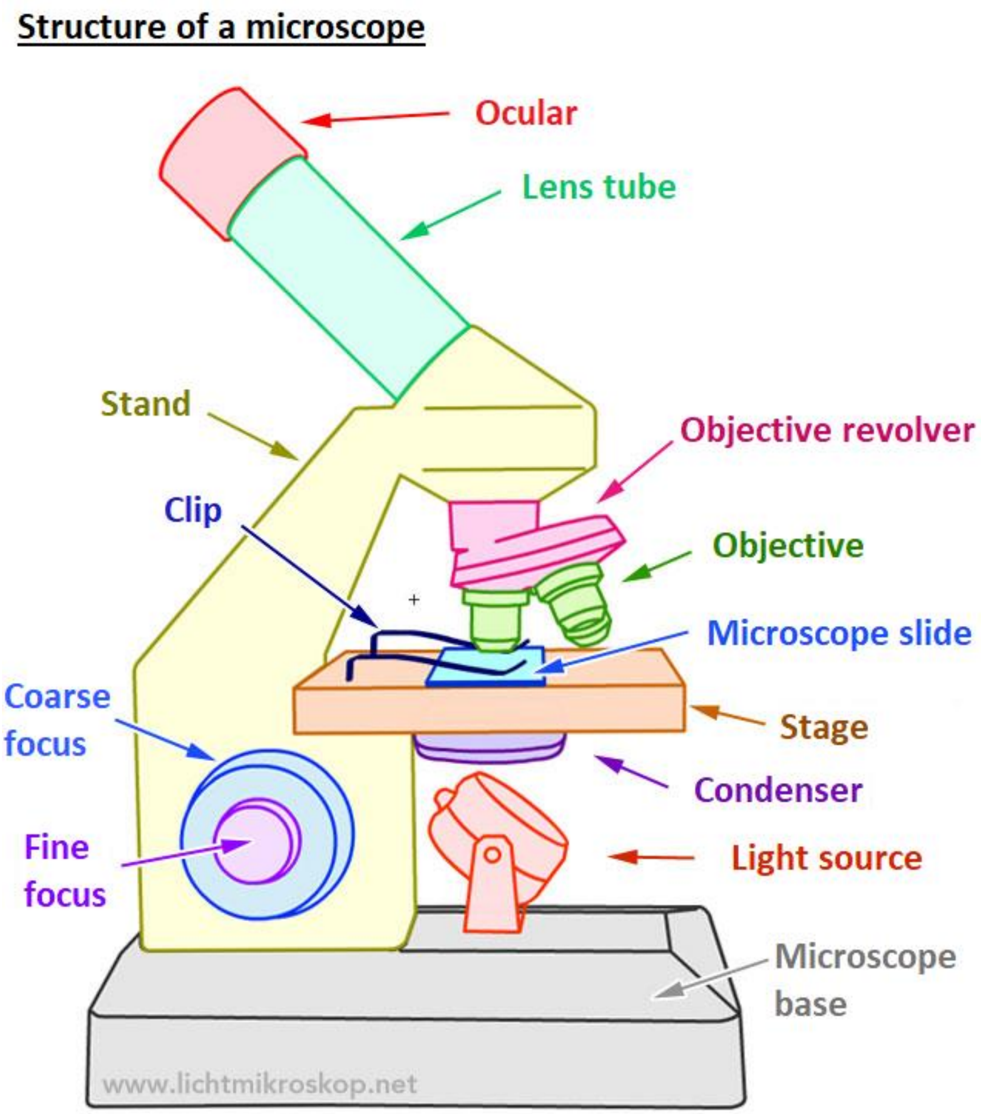


You might also notice what we did differently. Some things are missing in our microscope, which can you think of?

**Solutions to the questions:**

**Question 2: What kind of Lego objects do we have?**

A) It is important that you have understood that the object you built with Plan A gives light and can be switched on and off.

   The long stick can press on the small Lego lamp and switch the light on and off. It is important that everything is stable enough that it does not fall apart when you press it.

B) Did you also notice that you can use the object that you built with Plan B as a magnifying glass? To magnify something, you'll have to keep your head in the right position. From now on we will simply call this object the large magnifying glass.

C) Object C is probably the strangest object. Did you notice that you can move the slide vertically by turning the wheel? You can control a very precise and small movement of the slide here. That will be important later. This part is the hardest to build, so ask for help if you have any difficulties.

**Question 3: Inspection of the large magnifying glass**

You may also have noticed that you are somewhat restricted by the Lego housing, and that it is difficult to get enough light on the object you want to look at.
Thing that can be difficult:

- Distances are not so easy to control.
- When you put it down, no light falls on the object you want to look at.
- With larger distances it is difficult to hold everything still.

For the magnification: You are welcome to change the housing in which the lenses are inserted to improve the distances, or maybe to make everything more stable. Before you continue, it would be good to have everything back according to the original plan, so that we don't run into trouble later on.

**Question 4: How can we overcome the problem of illumination?**

There are, of course, many options. For example, you can remove one of the sides of part so that light can fall on the object. Another possibility that you might have come across is to put part B on part A. Then you have a lamp that you can even switch on and off.

**Question 5: How can we change the magnification?**

Take a look the magnifying glass at something simple, maybe the text of a book or a magazine. Now move your head further away from the magnifying glass, or bring it closer. What do you notice? Normally, the magnification increases as you move your head further away, but you also have to increase the distance to the object you want to look at.
But that doesn't go on forever. The further away you go, the more difficult it is to see everything well. And at some point it doesn't work anymore, right?

**Question 7: How can we further increase the magnification?**

Of course we can try to use other lenses. The shorter the focal length of the lenses, the greater the magnification. However, the lenses are also getting chunkier, so this won’t work forever.
Another suggestion could be to simply put two magnifying glasses in a row. That means that the second magnifying glass enlarges the image that the first magnifying glass has already enlarged.

**Question 9: Using two magnifying glasses.**

The alignment now becomes really difficult. Perhaps you already had the idea of building the small magnifying glass on object C yourself? Try that.
Now we only need a construction to get the big magnifying glass over the small one.

One way to build everything together is found in plan D.

To attach the small magnifying glass, we suggest the following. You can determine the rough position of the small magnifying glass by attaching it to different points on the vertical slide. Lego makes it possible 😊

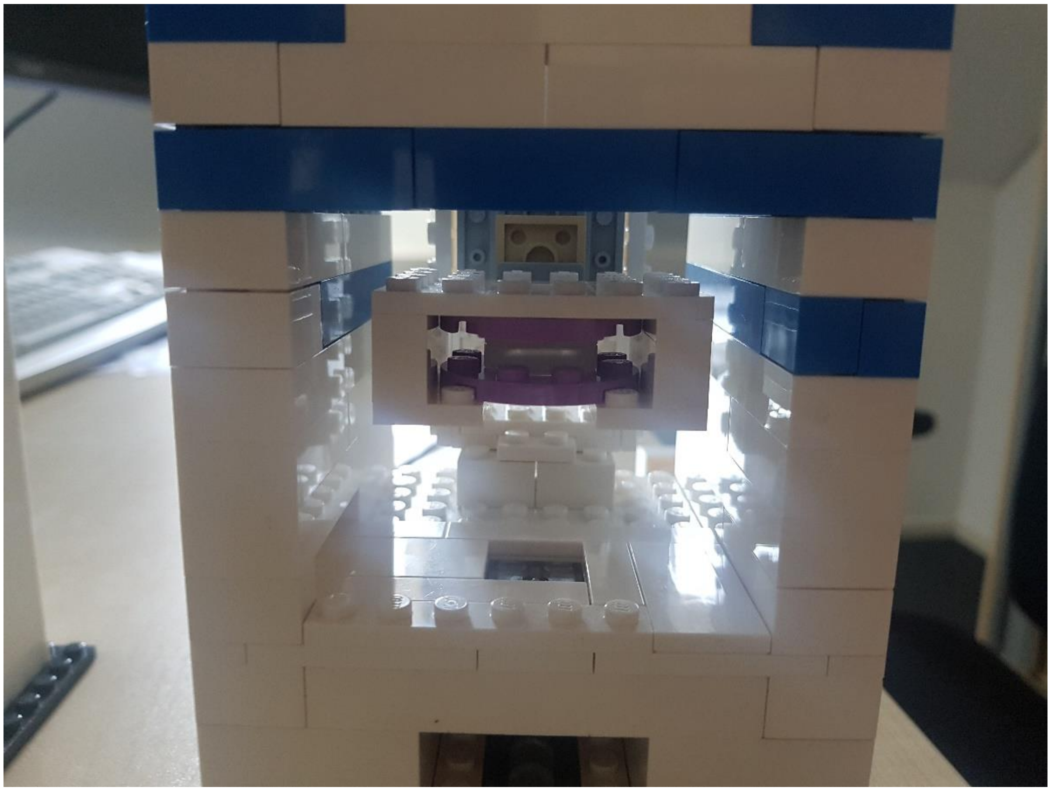

If you find the light too bright when looking through the lenses, you can cut out a piece of paper and place it on top of the lamp brick, below the object you're looking at. You can also cut out a larger piece of paper and place it on the flat Lego bricks, so that it is placed above the light.

On the next page you can see how the whole construction could look together.

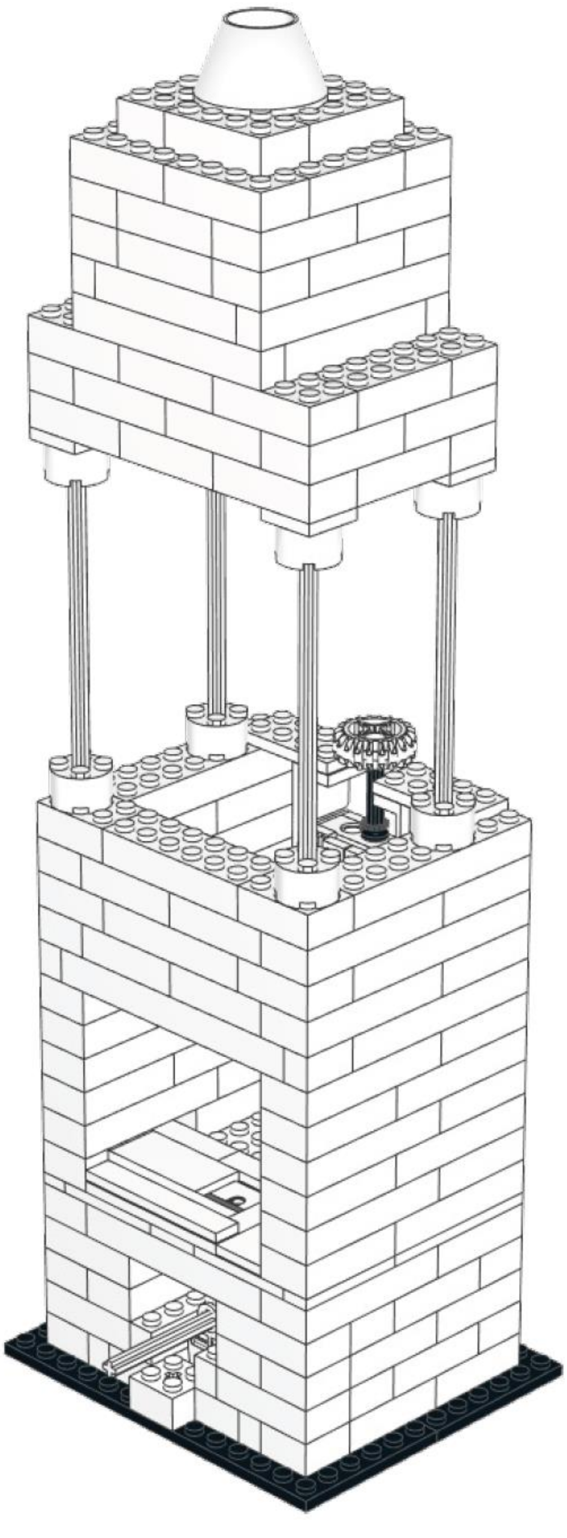

**Question 11: Video instructions for preparing samples**

The video “Preparation red onion cells” shows how to prepare red onion cells.

The video "Close sides sample" shows how to close two sides of a prepared sample with nail polish. This is what you need for the video "Osmotic shock experiment", with which you can investigate what happens when salty water flows over the onion cells. What happens to the cells? The best thing to do is to try recording it with your smartphone, then you can watch the effect again later and send the video to your friends.

Maybe you know the effect that occurs here: if you put salt on cucumber slices and wait a little, water droplets appear as the water is drawn out of the cucumber. This effect is called osmosis!

With the video "Preparation cheek cells" you can look at your own cells. You can get the cells out of your own mouth. Wipe the inside of your cheek with a cotton swab. You’ll have to use some pressure, otherwise it won't work. We need iodine to stain the cells a little, otherwise you won't see them. Caution: iodine can stain!

If you have access to brine shrimps, you can also try to prepare them. Simply dip your finger into the solution and place it on the slide. The crabs need air, so please do not leave them locked for too long.

**Question 12: High-magnification microscope with the smartphone lens.**

When you have built object 2 as objective into the microscope, you need to be a bit careful.

1) The two flat Lego pieces that were attached to the small magnifying glass also need to be removed when you take out the small magnifying glass. Otherwise the alignment is incorrect and you won’t see anything.
2) The magnification of the smartphone lens is great, but you’ll need to get very close to the object to see something. A good start is to put a piece of paper in the microscope and bring the objective so close, that you can see

the structure of the paper. If you can’t get any close, you can also attach the objective lower on object C.

3) You have a magnification of approximately 400x. That means that it’s difficult to position objects by hand since your movement is also 400x magnified! Be very careful and practice a little, then it should work. When you’ve found something exciting and want to show someone else, be careful not to move the microscope, or your object might move!

Try to take a look at a hair. What can you see?

**Question 13: SmartPhone compatibility.**

Take a look at Lego Plan E, you can definitely build that. But if you have a better idea, feel free to try it.
