## Supplementary File 4 for "Designing a high-resolution, LEGO-based microscope for an educational setting"

### Questionnaire A Name:

**1: To start a fire, you need ...**

- ☐ Oxygen (air)
- ☐ Fuel (e.g. wood)
- ☐ Heat
- ☐ All three

**2: Where is the flame of a candle at its hottest?**

- ☐ In the yellow region
- ☐ In the blue region
- ☐ Directly above the wick
- ☐ A few centimeter above the flame

**3: Which color is not part of the rainbow?**

- ☐ Yellow
- ☐ Red
- ☐ White
- ☐ Blue

**4: How does a printer produce different colors?**

- ☐ The colors are already mixed in the cartridge
- ☐ Through tiny colored dots close together
- ☐ The printer mixes the colors itself
- ☐ The colors are mixed on the paper

**5: Order from largest to smallest according to size:**

- ☐ Diameter of a hair > bacterium > virus
- ☐ Diameter of a hair > virus > bacterium
- ☐ Virus > Durchmesser eines Haares > bacterium
- ☐ Bacterium > diameter of a hair > virus

**6: If you turn a magnifying glass around and look through the other side, the image is...**

- ☐ Smaller than before
- ☐ As big as before
- ☐ As bi gas before, but mirrored
- ☐ Smaller than before and mirrored

**7: At least how many lenses do you need for a microscope?**

- ☐ 1
- ☐ 3
- ☐ 2
- ☐ 4

**8. In the picture you can see a microscope. Connect the terms with the corresponding parts on the microscope. If you can think of more terms, label them.**

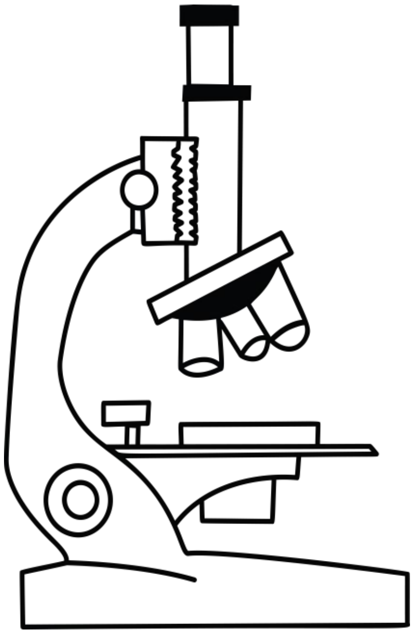

Objective

Light source

Ocular

Sample

**9: What do you change, when you focus a microscope?**

- ☐ The distance from the source to the sample
- ☐ The distance between the sample and the objective
- ☐ The distance between the ocular and the observer
- ☐ The light intensity

**10: When you look through a microscope, the image is ...**

- ☐ Mirrored left/right
- ☐ Mirrored top/bottom
- ☐ Mirrored left/right and top/bottom
- ☐ Oriented the same as the original

Draw the letter P as you would see it through a microscope:

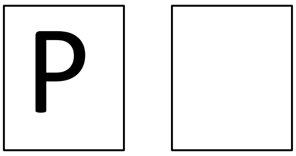
